# Towards a monograph of marine Verrucariaceae in Southern South America and maritime Antarctica: Integrative taxonomy reveals a previously unknown diversity

**DOI:** 10.1101/2025.01.22.634291

**Authors:** Daniel Fernández-Costas, Ulf Schiefelbein, Leopoldo G. Sancho, Asunción de los Ríos, Sergio Pérez-Ortega

## Abstract

The family of predominantly lichen-forming Verrucariaceae, contains several lineages that have adapted to live in the stressful conditions of the intertidal and supralittoral zones of rocky seashores in both hemispheres. The marine Verrucariaceae has been the subject of several systematic and taxonomic studies over the past few decades, although these studies have primarily concentrated on groups from the northern hemisphere. This study aims to address the paucity of taxonomic studies in the southern hemisphere by examining a substantial number of specimens collected in Southern South America and maritime Antarctica over the past two decades by the authors. We opted for an integrative taxonomic approach that joins the characterization and measurement of morpho-anatomical characters, coupled together with molecular barcoding to produce species delimitations to overcome the difficulties derived by the scarcity of taxonomic characters and the high plasticity of those characters. We barcoded a total of 301 specimens using the universal barcode un fungi (nrITS) and used single-locus species delimitation algorithms (ASAP, PTP, GMYC) to produce candidate species hypothesis that were corroborated with the morphological data. Our findings indicate that the taxonomic diversity of the Southern Hemisphere *Verrucaria* marina group has been significantly underestimated in previous studies. A total of 27 species were found in the region, 21 of which represent new taxa for science. Thorough descriptions together with illustrations showing the main characters of the species are provided for each taxon. In addition, we explored the systematics of the new taxa. We reconstructed the phylogenetic relationships of the family based on on six molecular markers (ITS, mcm7, nrSSU, nrLSU, mtSSU, RPB1). Our results showed that marine Verrucariaceae from the studied area belong to two distinct and non-related clades. A small group of species were related to the northern hemisphere species *Turgidosculum ulvae*. The second and more numerous group, which included the characteristic species *Mastodia tessellata* for a clade sister to the European genus *Verrucariopsis*. The systematic consequences of our findings are discussed. None of the previously reported species from the northern hemisphere and considered bipolar were found during our study.

## INTRODUCTION

The Verrucariaceae (Eurotiomycetes, Ascomycota), is a family of mostly lichen-forming fungi and lichenicolous fungi with 43 genera and about 950 species (Lücking *et al*. 2016). The type genus is *Verrucaria*, described by Schrader (1794). Morphologically, the species in this family are characterised by a perithecioid ascomata, bitunicate asci that dehisce through gelatinisation of the apical part of the outer wall of the ascus, hamathecium formed by periphyses located at the top of the ostiolar region and short pseudoparaphyses located below, and the lack of long intercalated sterile hyphae at the hymenial cavity at maturity, and hymenial gel usually reacting blue to potassium iodide (Gueidan *et al*. 2009; Orange *et al*. 2023; Schrader 1794)

The generic delimitation in this family has been the subject of debate over the years (Gueidan *et al*. 2007; 2009, Savić *et al*. 2008; Zahlbruckner 1903–1908, 1926; Zschacke 1933–1934). In the past, the separation between the different genera was primarily based on the use of six morphological characters: thallus structure (crustose, squamulose or foliose umbilicate), ascospore septation (simple, uniseptate, multiseptate, submuriform or muriform), the presence or absence of hymenial algae, upper cortex structure (absent, pseudocortex, eucortex, lithocortex), involucrellum (size and shape) and pycnidium structure (Dermatocarpon-type, Endocarpon-type or mucosa-type) (Gueidan *et al*. 2007; Yuzon *et al*. 2014). However, the use of molecular characters offered a new perspective, highlighting the problems with the use of these characters in generic delimitation, as most were shown to have evolved independently on multiple occasions throughout the evolutionary history of Verrucariaceae (Gueidan *et al*. 2007, 2009, 2011, 2022; Heiðmarsson *et al*. 2017; Navarro-Rosinés *et al*. 2007). In spite recent efforts towards the refinement of the systematics of the family (Gueidan *et al*. 2007, 2009, 2011, 2022; Heiðmarsson *et al*. 2017; Navarro-Rosinés *et al*. 2007; Pérez-Ortega *et al*. 2018), there is still a huge work to be done, with a number of groups still poorly studied and understood.

The family is found worldwide and can colonise a variety of substrates including soil, rock, wood, mosses and lichens, but the greatest diversity is found on rock. Verrucariaceae species are capable of thriving in a wide range of habitats, from arid and exposed areas to those completely submerged by water (Gueidan *et al*. 2009; Krzewicka 2016; Lamb 1948a; Santesson 1939; Thüs and Schültz 2009). The Verrucariaceae, together with the genera *Collemopsidium* and *Lichina*, are one of the most common groups of lichen-forming fungi in marine environments, occurring on rocky shores in temperate and cold regions of both hemispheres (Garrido-Benavent and Pérez-Ortega 2017; Hawksworth 2000, Tindall-Jones *et al*. 2023), thriving in the intertidal and supratidal zones of the rocky coasts, with one species reaching the subtidal zone (Jones *et al*. 2015, Lamb 1948a). It should be noted, however, that there are habitats that can be considered marine above the supralittoral zone, where species of this family also occur (Fletcher 1973a, b, 1980). Nevertheless, in this work, we always refer to the intertidal and supratidal zones whenever we talk about marine environments.

Marine lichen-forming fungi have been the subject of limited study throughout history (Jones *et al*. 2009). The first major classification of aquatic fungi, by Johnson and Sparrow (1961), did not consider lichenized fungi, only mentioning their occurrence in such environments. Nevertheless, there were already existing compilations lichen-forming fungi in these environments in certain areas (Lamb 1948a; Santesson 1939; Zschacke 1927).

The first descriptions of marine lichen-forming fungi were published in the beginning of the 19th century by the Swedish naturalist Wahlenberg in 1803 (in Acharius 1803). He provided descriptions of four species: *Verrucaria ceuthocarpa*, *Hydropunctaria maura* (≡ *V. maura*), *Wahlenbergiella mucosa* (≡ *V. mucosa*), and *W. striatula* (≡*V. striatula*), all collected in the province of Finnmarkin north Norway.

In addition to *Verrucaria*, several other genera of Verrucariaceae are known to occur in marine environments, namely *Hydropunctaria*, *Mastodia*, *Turgidosculum*, *Verrucariopsis* and *Wahlenbergiella* (Gueidan *et al*. 2009, 2022; Kohlmeyer and Kohlmeyer 1979; Pérez-Ortega *et al*. 2010).

The genus *Hydropunctaria* was described by Gueidan *et al*. (2009). All species within the genus are associated with aquatic ecosystems, either inhabiting freshwater habitats or marine environments. In the latter case, they typically occur in the supralittoral zone, which is exposed to water spray. The type species of the genus is *Hydropunctaria maura*, with records in rocky coasts in temperate to cold areas from both hemispheres. However, the recent description of several semi-cryptic species in Europe that were previously included in the concept of this species has cast doubt on the identity of the populations in the Southern Hemisphere (Orange 2013). The genus is characterised by the presence of crustose thalli, which are subgelatinous when wet and often interrupted by black punctae or columns. The upper cortex is weakly differentiated, while the medulla is either not differentiated or forms a black basal layer. The algal layer is composed of algal cells that are typically arranged in vertical columns. The involucrellum often exhibits an uneven or rough upper surface, and the ascospores are simple and typically range in length from 10 to 18 μm (Gueidan *et al*. 2009; Orange 2013).

The genus *Mastodia* was first described by Hooker f. et Harvey in 1847 for the type species *Mastodia tessellata* (Hooker 1847). The initial description considered the taxon as an alga, and it was not until recently that its nomenclatural status was clarified (Kohlmeyer *et al*. 2004). The genus is essentially maritime, occurring in near-shore areas that are often affected by sea spray (Kohlmeyer *et al*. 2004). It was previously classified as part of the Mastodiaceae, a monotypic family characterized by the presence of the green algae foliose photobiont of the genus *Prasiola*, unitunicate asci and septate spores (Zahlbruckner 1907). However, Pérez-Ortega *et al*. (2010) demonstrated the presence of bitunicate asci, which when mature become unitunicate due to the gelatinization of the outer layer, as well as the presence of simple ascospores. Furthermore, they presented molecular data that substantiated its classification within the family Verrucariaceae. This species is mainly found in the southern hemisphere, but populations are also known from the northern hemisphere (British Columbia and Alaska) (Garrido-Benavent *et al*. 2018). Of the 7 species in the genus, most studies have focused exclusively on the type species, particularly its distinctive photobiont and characteristic bipolar distribution pattern, which originated in the Southern Hemisphere (Garrido-Benavent *et al*. 2018, 2017; Dodge 1948).

The genus *Wahlenbergiella*, with *W. mucosa* as type species, was introduced by Gueidan *et al*. in 2009. The species is characterised by a green to olive-green subgelatinous thalli, simple ascospores and mucosa-type pycnidia (Gueidan *et al*. 2009). So far, three species have been included in the genus, all of them occurring in the intertidal zone. *Wahlenbergiella mucosa* and *W. striatula* occur in the coasts of Europe and North America in the Northern Hemisphere and they have been recorded is several areas of the Southern Hemisphere (Galloway 2007; Orange 2013; Santesson 1939; Vail and Walker 2021). The third species included in the genus was *W. tavaresiae* (Gueidan *et al*. 2011), a curious taxon restricted to the western coast of the USA which forms symbiosis with the brown seaweed *Petroderma maculiforme* (Moe 1997; Sanders 2004). However, morphologically there is a lack of useful set of characters features to characterize and circumscribe *Wahlenbergiella* (Gueidan *et al*. 2011).

The genus *Turgidosculum* was first introduced by Kohlmeyer and Kohlmeyer (1972) for the species *T. ulvae* (=*Guignardia ulvae*). It is exclusively marine, occurring in intertidal and often exposed to strong waves, distributed along the west coast of the United States and British Columbia (Kohlmeyer and Kohlmeyer 1979). The diagnostic features of this genus are the particular foliose photobiont, a green alga belonging to the genus *Blidingia*, and the size of its spores. (Kohlmeyer and Kohlmeyer 1979; Pérez-Ortega *et al*. 2018). In addition to *T. ulvae*, the species *Leptogiopsis complicatula* described by Nylander from the Beringian Strait (Nylander 1884) was combined in *Turgidosculum.* However, *T. complicatulum* is currently regarded as a synonym of *Mastodia tessellata* (Kohlmeyer *et al*. 2004). Pérez-Ortega *et al*. (2018) examined the *Turgidosculum ulvae* symbiosis studying the phylogenetic affinities of both symbionts and their interactions at the ultrastructural level. They showed that *T. ulvae* were closely related to *Verrucaria ditmarsica*, a taxon from the northern hemisphere. The two species constitute a discrete lineage, separated by an unusual long branch from the remainder of the Verrucariaceae.

The most recently described genus of marine Verrucariaceae is *Verrucariopsis* (Gueidan *et al*. 2022). The genus, which contains two species, is distinguished by the size of its spores and its plurilocular pycnidia. The type species, *V. suaedae*, is an epiphytic fungus that grows on the lower parts of *Suaeda vera* stems in estuaries. The other species of this genus, *V. halophila*, is a rock-dwelling fungus that is found in the intertidal zone (Gueidan *et al*. 2022).

Although the knowledge of marine lichen-forming has been largely biased towards the Northern Hemisphere (e.g. Erichsen 1928, 1937; Gueidan *et al*. 2022; Harada 1995; Nylander 1858, 1863; Ryan 1988; Santesson 1939), several authors have focused on the diversity of the group in the Southern Hemisphere.

William Nylander, described several taxa of marine Verrucariaceae from the Southern Hemisphere, mostly based on specimens collected by other colleagues (Ahti 1990; Nylander 1891). He described *V. microspora* from Chile and *V. tessellatula* from Kerguelen (Crombie 1875; Nylander 1855). The Finnish lichenologist Edward Vainio described several taxa of marine Verrucariaceae from the Gerlache Strait in Antarctica (*i.e. V. cylindrophora*, *V. dispartita* and *V. racovitzae*) (Vainio 1903). The Austrian-Hungarian lichenologist Alexander Zahlbruckner also interested in the diversity of this group, describing two taxa from the Southern Hemisphere. *Verrucaria chiloensis*, from the Chiloe Island in Chile, and *V. aucklandica* from Anawhata in New Zealand (Zahlbruckner 1917, 1941).

Mackenzie Lamb, who undertook several expeditions to polar areas of the Southern Hemisphere, published the first overview of marine Verrucariaceae in Antarctica, included in his review of Antarctic pyrenocapic lichens (Lamb 1948a), with the description of two species, including the first known species of lichen-forming fungi able to thrive completely submerged, below the lowest ebb-tide level, *V. serpuloides* (Lamb 1948a, 1955, 1973).

The collective findings of the research conducted over the past two decades by various authors (Garrido-Benavent *et al*. 2018; Pérez-Ortega *et al*. 2010; Černajová *et al*. 2022), and the recent systematic changes in the family (Garrido-Benavent *et al*. 2018; Gueidan *et al*. 2009, 2011, 2022; Pérez-Ortega *et al*. 2018), have collectively underscored the urgent necessity for a taxonomic and systematic reassessment of the marine Verrucariaceae group in the Southern Hemisphere.

The objective of this study is, therefore, to enhance our understanding of the actual diversity of marine species within the genus *Verrucaria* in South America and maritime Antarctica. Furthermore, this study aims to provide a systematic framework for the marine lineages within this hemisphere.

## MATERIALS AND METHODS

### Sampling

In the period between 2008 and 2024, a total of approximately 50 localities in the Southern Hemisphere in Antarctica, Livingston Island, the Falkland Islands and Chile were surveyed, resulting in the collection of more than 300 specimens of the genus Verrucaria. The specimens have been collected by Sergio Pérez Ortega (RJB, Madrid, Spain), Alan Orange (MHN, Cardiff, Wales), Ulf Schiefelbein (Botanical Garden, University of Rostock, Rostock, Germany) and Asunción de los Ríos (MNCN, Madrid, Spain).

A total of 301 specimens were subjected to a comprehensive review. Unless otherwise indicated, all specimens will be deposited in the herbarium of the Royal Botanical Garden-CSIC (MA-Lich).

### Morphological analyses and characterisation

Specimens were examined using a Nikon SMZ1000 stereomicroscope. Hand-cut sections of perithecia and thalli were observed using either an Olympus BX51 with Nomarski differential interference contrast (DIC) or a Nikon Eclipse E200 microscope fitted with a set of polarized filters. Images were captured with a Leica DMC 4500 digital camera fitted to the Olympus microscope. Macroscopic pictures were taken either with a Leica S8i stereomicroscope with a built-in digital camera of with an OM-1 digital camera fitter with and Olympus 90 mm macro lens. Colour reactions were observed using 10% KOH (K), 8% sodium hypochlorite (C) and Lugol’s iodine solution, the latter both with (K/I) and without pretreatment with K. The measurements of ascospores were made on material mounted in water and are presented as minimum – (mean ± standard deviation) – maximum values followed by the number of measurements (n) and number of specimens (s).

We characterized morphological features previously reported in the literature as relevant for the group (Gueidan *et al*. 2007, 2009, Lamb 1948a, Orange 2013). The following characteristics were observed: 1) prothallus: presence, colour, consistency; 2) thallus: colour, height, appearance; 3) thallus structures: presence of punctae, cracks, or ridges, 4) pseudocortex: presence, size, colour, K reaction; 5) mycobiont cells: size, shape, arrangement; 6) photobiont cells: size, shape, arrangement; 6) perithecia: appearance, shape, size, colour, I and K/I reaction of the hymenial gelatin, colour and shape of the ostiolar region, position, 7) involucrellum: size, colour, layer extent; 8) excipulum: size, colour, number of cell layers, 9) asci: shape, size, number of ascospores; 10) periphyses: presence, shape, and size, 11) ascospores: shape, colour and size, 12) pycnidia: arrangement on the thallus, shape, colour, size, 13) pycnidiospores: shape, size, and colour.

### Molecular methods

The preparation of the samples was carried out in a Nikon SMZ-445 binocular lens with the help of sterile dissecting forceps and a razor blade. Two to three clean areolas per sample (c. 2-3 mm^2^) were excised and placed in sterile Eppendorf tubes. After one hour of freezing, they were pulverized using TissueLyser II (Qiagen) with two glass beads. Genomic DNA was extracted using E.Z.N.A.^®^ Forensic DNA Kit (Omega Bio-Tek), following the instructions of the manufacturer, resulting in the elution of 60 µl of DNA in Elution Buffer. We amplified six loci: the nrITS region (transcribed intergenic sequences 1 and 2 and 5.8S unit of the nuclear ribosomal RNA gene), nrLSU (nuclear large ribosomal subunit), nrSSU (nuclear small ribosomal subunit), mtSSU (mitochondrial small ribosomal subunit), mcm7 (minichromosome maintenance component 7) and RPB1 (larger subunit of RNAII polymerase). The primers used for the amplification of each of these regions are available in Table 1.

**Table 1.** Primers and PCR conditions used to amplify ITS, mcm7, mtSSU, nrLSU, nrSSU, RPB1. The symbol * indicates a touchdown 0,5°C for each cycle.

| Region | Primers | Publications | PCR conditions |
| --- | --- | --- | --- |
| ITS | ITS1F | Gardes & Bruns (1993) | 2 min 95°C, 35x (1 min 92°C, 1 min 56°C, 1 min |
|  | ITS4 | White *et al.* (1990) | 72°C), 7 min 72°C. |
| mcm7 | mcm7-709for | Schmitt *et al.* (2009) | 10 min 94°C, 14x (45 sec 94°C, 50 sec 58.5°C*, |
|  | mcm7-1348rev |  | 1 min 72°C), 30x (45 sec 94°C, 50 sec 56°C, 1 min 72°C), 5 min 72°C. |
| mtSSU | mrSSU1 | Zoller *et al.* (1999) | 3 min 94°C, 9x (1 min 94°C, 1 min 56°C*, 1.3 |
|  | mrSSU3R |  | min 72°C), 30x (45 sec 94°C, 50 sec 56°C, 1 min 72°C), 5 min 72°C. |
| nrLSU | LR0R | Rehner & Samuels (1994) | 1 min 95°C, 9x (45 sec 95°C, 40 sec 56°C*, 2.3 |
|  | LR5 | Vilgalys & Hester (1990) | min 72°C), 30x (45 sec 95°C, 40 sec 52°C, 2.3 min 72°C), 10 min 72°C. |
| nrSSU | NSSU97A | Kauff & Lutzoni<br>(2002) | 5 min 94°C, 9x (1.15 min 94°C, 1.3 min 56°C*,<br>1.45 min 72°C), 30x (1.15 min 94°C, 1.3 min<br>52°C, 1.45 min 72°C), 15 min 72°C. |
|  | NSSU1088 |  |  |
| RPB1 | RPB1-Afasc | Hofstetter *et al.* (2007) | 3 min 95°C, 14x (45 sec 95°C, 1.3 min 57°C*,<br>1.3 min 72°C), 35x (45 sec 95°C, 1.3 min 52°C,<br>1.3 min 72°C), 10 min 72°C. |
|  | RPB1-6R1asc |  |  |

PCR reactions for ITS and LSU were prepared in a 15 µl final volume containing a mix made up of 6.5 µl of MyTaq™ Red Mix (Bioline®), 0.5 µl or 1 µl of each primer (10 µM), 4.5 µl or 3.5 µl of distilled water and 3 µl of DNA template. For nuSSU, mtSSU, mcm7 and RPB1, PCR beads (PuReTaq Ready-To-Go) (GE HealthCare®, UK) were used, following manufacturer’s instructions and using 3 (for nuSSU and mtSSU) or 6 (for mcm7 and RPB1) µl of DNA template, 3 µl of each primer and 19 or 16 µl of distilled water, for a total final volume of 25 µl. The amplification settings used for the PCR are presented in Table 1.

PCR products were run in 2% agarose gels stained with SYBR™ Safe DNA Gel Stain (Thermo Fisher Scientific). DNA was sequenced in Macrogen Spain (Madrid, Spain) with the same primers used in the PCRs.

### Sequence alignments

We performed multiple sequence alignment for each marker using MAFFT v.7.409 (Katoh *et al*. 2002, Katoh & Standley 2013) as implemented in Geneious® v. 2024.0.3 with the following parameters: Auto algorithm, the 200PAM / k = 2 scoring matrix, a gap open penalty of 1.53 and an offset value of 0.123. The alignment process was carried out in nucleotide mode. Once aligned, we divided the ITS into three regions, namely ITS1, 5.8S and ITS2. Ambiguous regions and introns were delimited manually and excluded from the alignment. For the six amplified regions ITS1, 5.8S, ITS2, nrLSU, nrSSU, mtSSU, mcm7, *RPB*1 region A–D, Gblocks v 0.91b (Castresana 2000) was employed to eliminate introns and ambiguously aligned positions, utilising the less stringent conditions as implemented in the web version (http://molevol.cmima.csic.es/castresana/Gblocks_server.html).

In order to phylogenetically place our group within the family Verrucariaceae, we downloaded sequences of all markers, with the exception of ITS, for several representatives of the family. The Genbank accession number was obtained from the articles by Gueidan *et al*. (2022), Heiðmarsson *et al*. (2017) and Pérez-Ortega *et al*. (2018).

Nucleotide substitution models were estimated for each separate genomic region using the Akaike Information Criterion (AIC) implemented in jModelTest v 2.1.10 (Darriba *et al*. 2012). Conflicts among partitions were eliminated by pruning out problematic sequences or taxa.

### Phylogenetic analysis

Three different datasets were constructed for the purpose of investigating phylogenetic relationships. The datasets were designated as D1 (ITS1 + 5.8S + ITS2), D2 (nuLSU + nuSSU + mtSSU + mcm7 + RPB1), and D3 (ITS1 + 5.8S + ITS2 + nuLSU + mtSSU + mcm7).

We generated an ultrametric phylogenetic tree using BEAST 1.10 (Suchard and Rambaut 2009; Drummond *et al*. 2012), which is the input for some of the species delimitation analyses (see below). This analysis was performed only with the D1 dataset. The input file for BEAST was generated using BEAUti v 1.10.4 (Drummond *et al*. 2012). The analysis used a lognormal uncorrelated relaxed clock, the GTR + I + G substitution model, and a random starting tree. We chose the “Speciation model: Birth-Death process”and a random starting tree with 50 x 10^6^ MCMC generations and sampling every 1000 generations. We used Tracer 1.7.1 (Rambaut *et al*. 2018) to check for convergence of chains. RaxML, MrBayes and BEAST inferences were carried out in CIPRES Science Gateway (Miller *et al*. 2011).

Maximum Likelihood (ML) and Bayesian approaches were used for inferring phylogenetic relationships for datasets D1, D2 and D3. Multi-locus and single-locus ML trees were calculated using RaxML (Stamatakis 2014) with GTRGAMMA model for bootstrapping phase, 1000 rapid bootstrap pseudoreplicates were performed to evaluate nodal support and the option to perform a rapid Bootstrap analysis and search for the optimal ML tree in a single program run was selected. Nodes with bootstrap values equal or higher than 70% were considered significant. Bayesian trees, for D2 and D3, were calculated using MrBayes 3.2 (Ronquist *et al*. 2012). Starting with a random tree, two simultaneous, parallel four-chain runs were executed over 50 x 10^6^ generations and sampled after every 1000th step. The first 20% of data was removed as burn-in. The 50% majority-rule consensus tree was calculated from the remaining trees. Nodes with posterior probabilities equal or higher than 95% were considered significant. ML and Bayesian single-locus phylogenies were used to check for topological incongruences.

### Species discovery strategies based on single locus datasets

We employed three species discovery algorithms to each individual marker: The Assemble Species by Automatic Partitioning (ASAP, Puillandre *et al*. 2021), the Generalized Mixed Yule Coalescent model (GMYC, Pons *et al*. 2006; Fujisawa and Barraclough 2013) and the Multi-rate Poisson tree processes for single-locus (mPTP, Kapli *et al*. 2017).

ASAP is a species discovery algorithm that computes a pairwise genetic distance matrix and attempts to find the so-called ‘barcode gap’ between the small intraspecific genetic distances and the larger interspecific ones. The algorithm generates multiple species hypotheses accompanied by a p-value and a measure of the width of the barcode gap. ASAP was run at https://bioinfo.mnhn.fr/abi/public/asap/ using the K80 model to calculate pairwise genetic distances. The required transition to transversion ratios (TS/TV) were calculated in MEGA 5.2 (Tamura *et al*. 2011).

In contrast ASAP, GMYC and mPTP are species discovery algorithms that need a phylogenetic tree as input. The primary distinctions between GMYC and mPTP lie in their respective approaches to evolutionary processes. GMYC is founded upon the differentiation between speciation and coalescence, whereas mPTP is predicated upon the modelling of mutation rates between lineages, obviating the necessity for an explicit breakpoint between processes. In essence, GMYC is a more appropriate tool for well-resolved and relatively simple trees, whereas mPTP is better suited to scenarios where there are variations in mutation rates between lineages or where the phylogenetic tree is not as robust. They estimate the branching rates in different regions of the tree and try to identify which part follows a speciation model and which part follows a coalescent model. The species hypothesis is generated by the identification of a threshold that maximizes the transition between the two branching rates. GMYC was run using the *gmyc* function of the R *splits* package (https://rdrr.io/rforge/splits/). Species hypotheses were calculated following single and multiple threshold models. As input, we used the ultrametric tree previously created by BEAST mPTP was run at https://mptp.h-its.org/#/tree with 5×10^5^ MCMC generations and a burn-in of 30%. The ML phylogenetic tree previously generated for D1 was used as input data.

## RESULTS

### Sequence data

In this study we generated 220 new sequences from 6 different regions, 144 of which correspond to the ITS region (ITS1 (0.25 kb), 5.8S (0.16 kb), ITS2 (0.23 kb), 31 to the nrLSU (1.3 kb), 2 to the nrSSU (1.6 kb), 24 to the mtSSU (1.1 kb), 18 to the mcm7 (0.5 kb) and 1 to the RPB1 (1.2 kb). The total length of the 3 different alignments concatenated multilocus alignments after using Gblocks were: 0.6 kb for D1 (ITS1 + 5.8S + ITS2), 7.8 kb for D2 (nuLSU + nuSSU + nuSSU + mtSSU + mcm7 + RPB1) and 7.6 for D3 (ITS1 + 5.8S + ITS2 + nuLSU + mtSSU + mcm7).

The best substitution models for the different regions returned by jModelTest are reported in Table 2.

**Table 2.** Substitution models return by jModelTest for the different regions.

| Region | Substitution model |
| --- | --- |
| *ITS1* | GTR + I + G |
| *5.8S* | GTR + G |
| *ITS2* | GTR + G |
| *nrLSU* | GTR + I + G |
| *nrSSU* | GTR + I + G |
| *mtSSU* | GTR + I + G |
| *mcm7* | GTR + I + G |
| *RPB1* | GTR + I + G |

### Phylogenetic relationships

The phylogenetic relationships between members of the Verrucariaceae inferred using the maximum likelihood and Bayesian approaches produced congruent topologies can be seen instead of the Figure 1. The MrBayes topology is shown. In general, both topologies showed a high support of the relationships, except for the relationships at the backbone.

**Figure 1.**
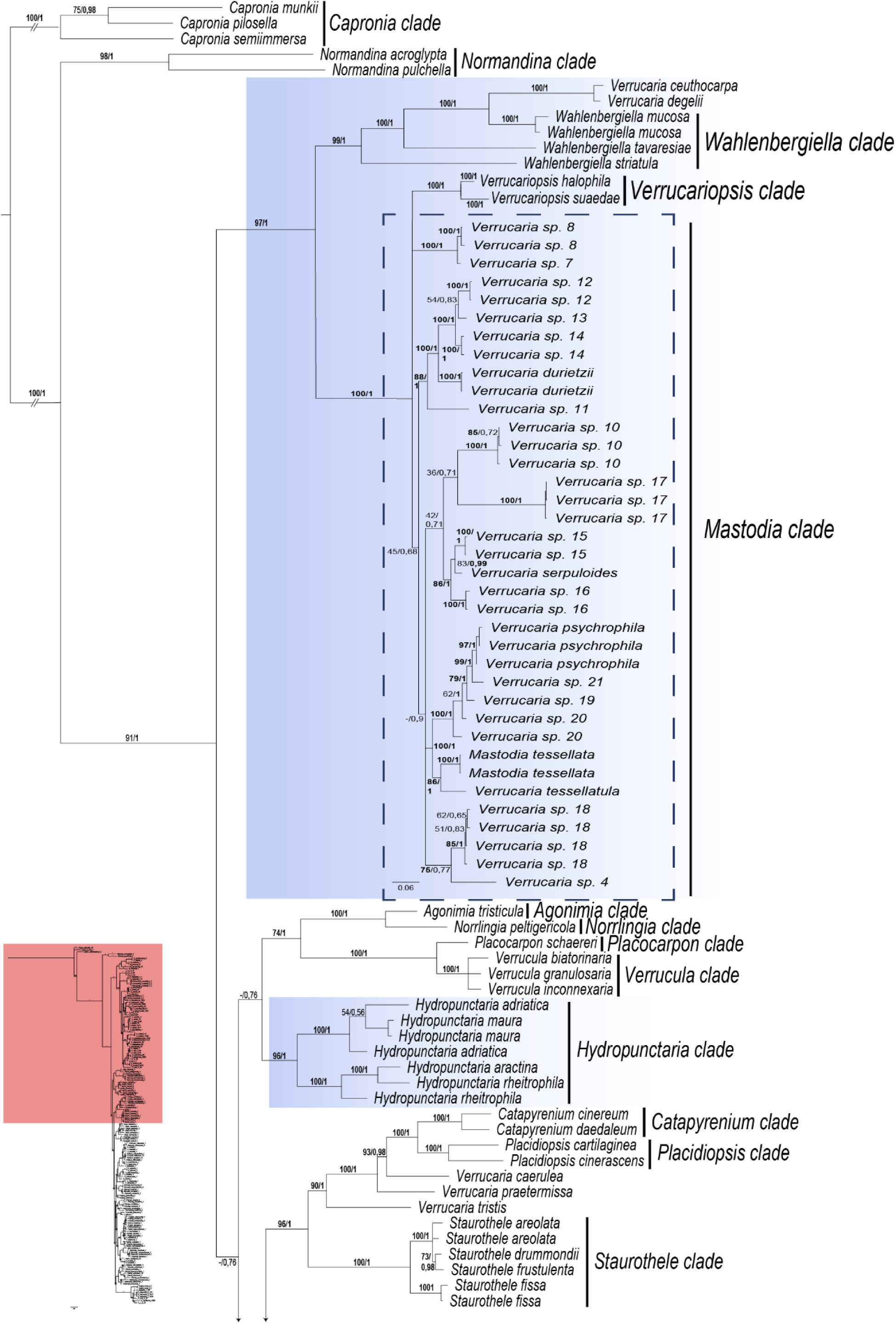
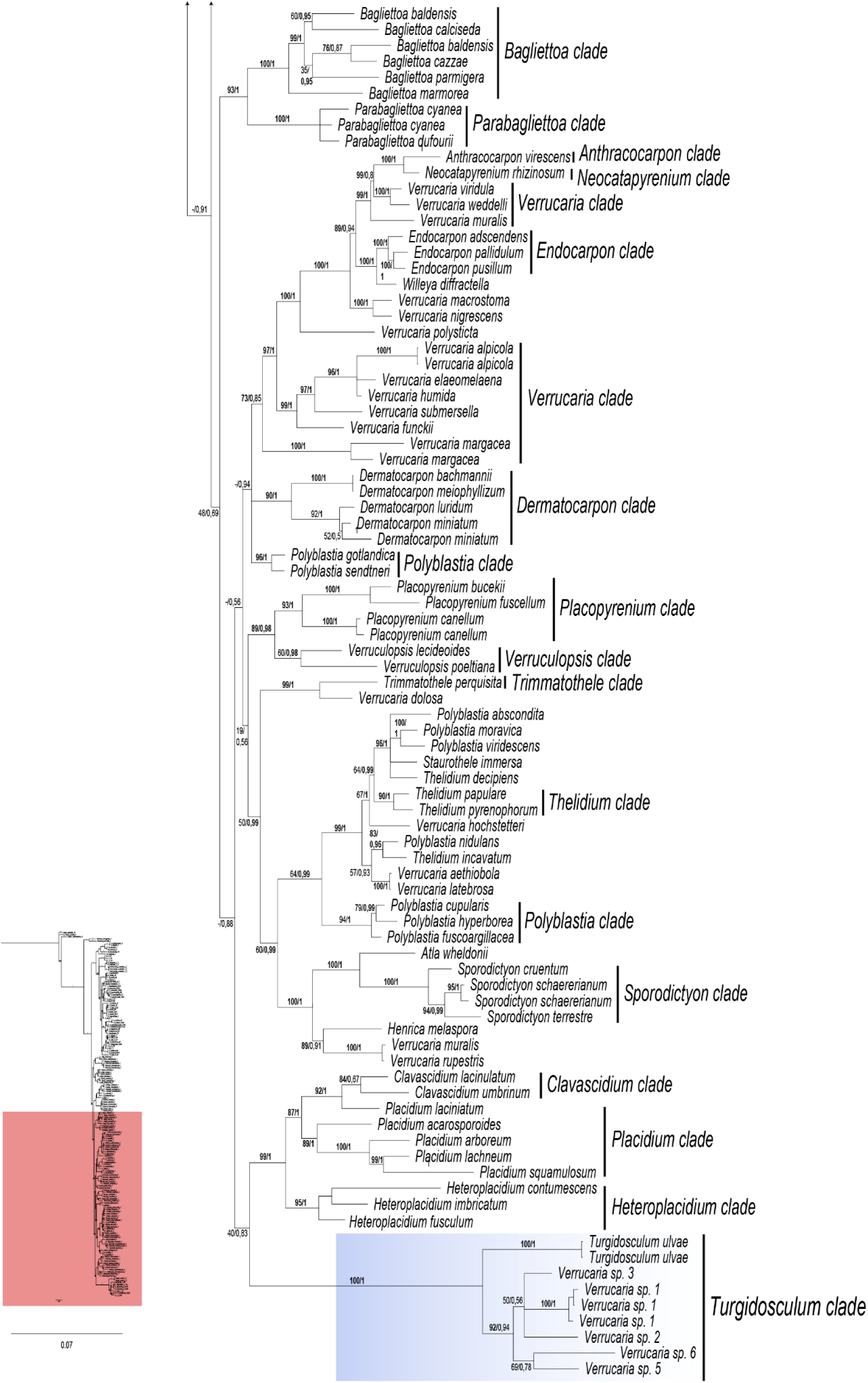
Bayesian phylogram obtained in MrBayes. Shows phylogenetic relationships among Verrucariaceae family. Branch labels in bold indicate statistical support (bootstrap(BP)≥70 and/or posterior probalities(PP)≥0.95). The general tree was constructed using dataset D2, with the exception of the clade enclosed within the dashed box, which was generated using dataset D3. The blue boxes indicate marine clades exclusively. Processed by the author with Illustrator.

The recovered phylogenetic hypotheses revealed the existence of three major exclusively marine groups supported by both BP and PP (highlighted with a blue box), which are placed in different positions of the phylogenetic tree.

The first of the marine clades corresponded to species of the genus *Hydropunctaria* (BP:96; PP:1). The second marine clade, Verrucaria species from the southern hemisphere form a clade with *Mastodia*. However, this is not supported (BP:45; PP:0,68). *Verrucariopsis* forms a supported clade (BP:100; PP:1). The *Wahlenbergiella* species, together with *V. ceuthocarpa* and *V. degelii*, form a supported clade (BP:99; PP:1), basal to the clade formed by *Verrucariopsis* and *Verrucaria s. l.* of the southern hemisphere. A third marine clade is supported (BP:100; PP:1), formed by *Verrucaria* species from the southern hemisphere together with *Turgidosculum ulvae*. The *Turgidosculum ulvae* clade (BP: 100; PP: 1) is sister to the clade of *Verrucaria* species from the southern hemisphere (BP: 92; PP: 0,94).

In the *Mastodia* and *Turgidosculum* clades the backbone relationships were poorly supported.

### Species delimitation

The single-locus species delimitation results from ASAP, mPTP and GMYC approaches are summarized in Figure 2. The ASAP and mPTP algorithms gave similar results, 26 and 24 candidate species respectively, as did GMYC single and multiple, 31 and 35 candidate species. However, ASAP and mPTP gave much more conservative results with respect to barcode changes than GMYC. Analyses using the *Turgidosculum* and *Mastodia* clades separately did not change the number of inferred putative species.

**Figure 2.**
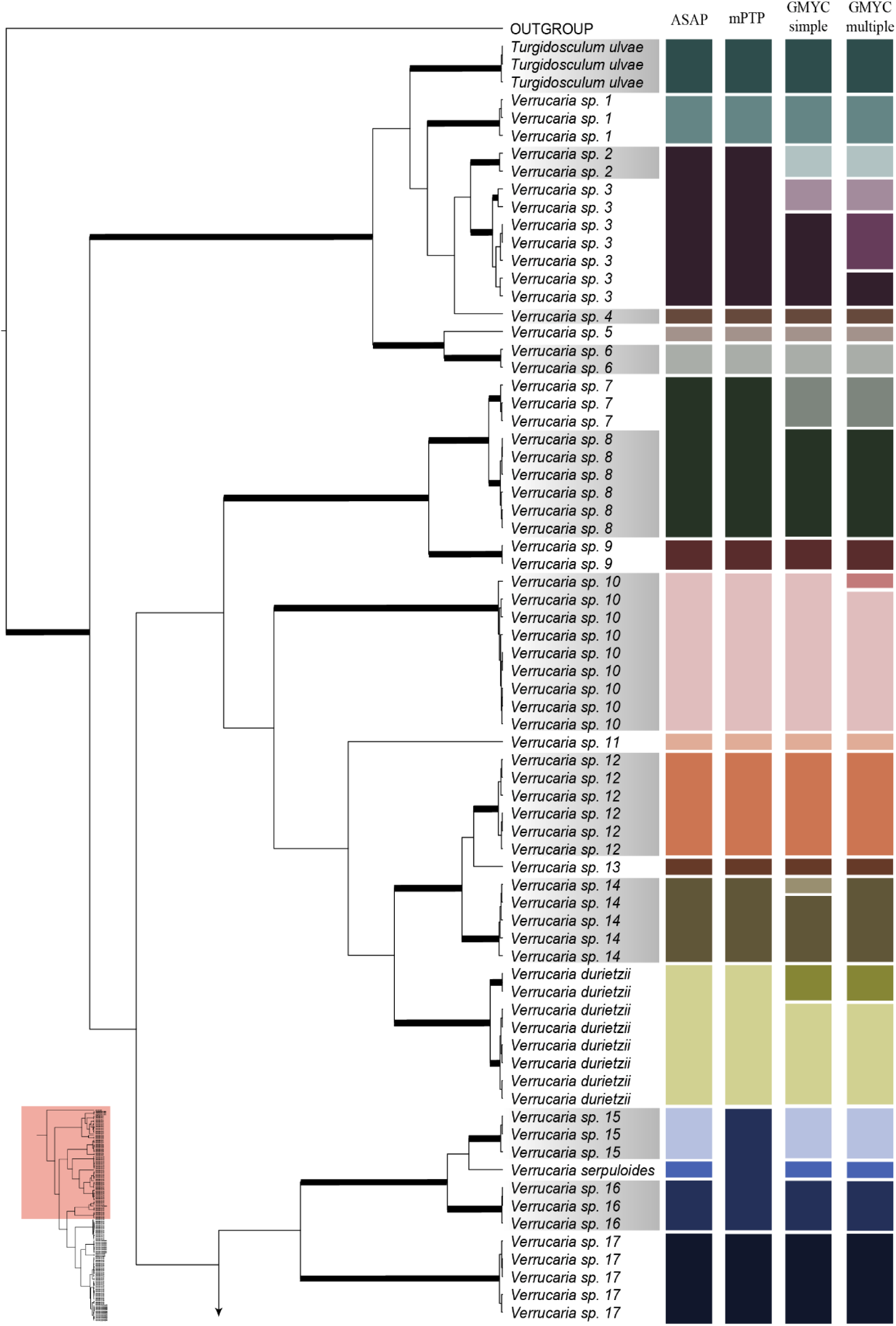
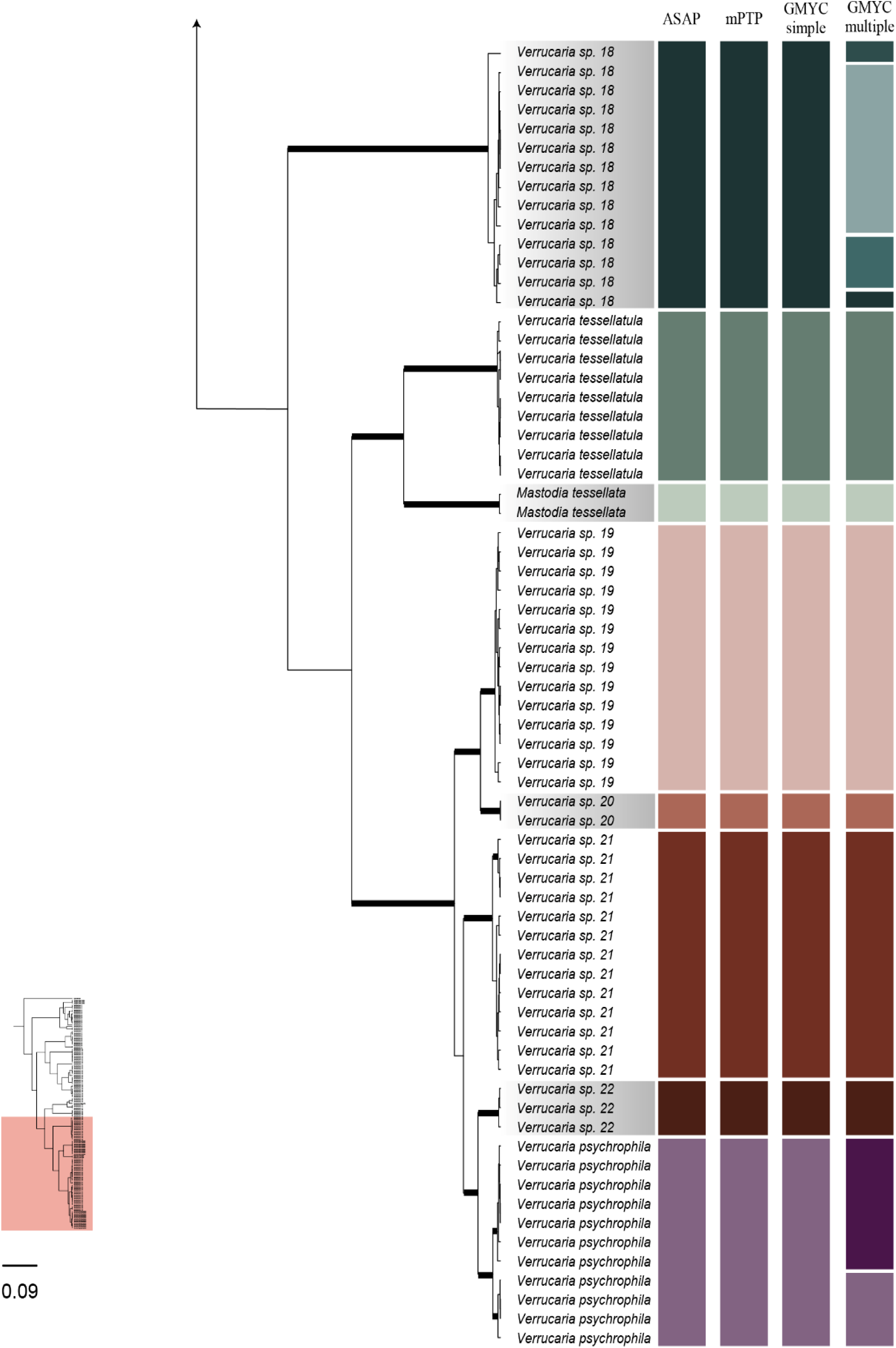
Maximum clade credibility tree from the BEAST analysis. Branches with posterior probability ≥ 0.95 or ≥ 0.70 bootstrap are highlighted in bold. The table on the right side provides further information for all accessions: more biologically plausible species hypotheses returned by ASAP (ITS, column 1), bPTP (ITS, column 2), GMYC simple (ITS, column 3) and GMYC multiple (ITS, column 4). The species finally proposed are those marked in the names of the phylogeny tips. Processed by the author with Illustrator.

In the clade formed by species 2 and 3, the ASAP and mPTP group them into a single species, while the simple GMYC gives 3 putative species, species 2 and 3 as two species. The multiple GMYC even distinguishes three species within species 3 in addition to species 2.

In the clade formed by species 7 and 8, ASAP and mPTP group them into a single species, while single and multiple GMYC separate the two species.

In the clade of species 10, ASAP, mPTP and single GMYC yield a single species, while multiple GMYC separates the two species. And in the clade formed by species 14, ASAP, mPTP and multiple GMYC yield a single species, while single GMYC separates the two species.

In the clade formed by *V. durietzii*, the ASAP and mPTP yielded a single species, whereas the single and multiple GMYC separated two species. In the clade formed by species 15, 16 and *V. serpuloides*, both the ASAP and the simple and multiple GMYC yielded three species, whereas the mPTP yielded a single species.

In the clade formed by species 18, the ASAP, mPTP and single GMYC methods yield a single species, whereas the multiple GMYC method results in the separation of three species.

In the clade formed by *V. psychrophila*, the application of ASAP, mPTP and single GMYC results in the identification of a single species. Conversely, the utilisation of multiple GMYC leads to the delineation of two distinct species.

Figure 2 depicts the results from the species delimitation analyses as well as the consensus decission based on the candidate species and the morphological and anatomical data. A total of 28 taxa were finally accepted as actual species, of which six are currently known taxa and the other 21 are new to science.

### Taxonomy

#### Mastodia tessellata

(Hook. f. & Harv.) Hook. f. & Harv., Bot. Antarc. Voy.: 499 (1847); — MycoBank MB#395110;

≡ *Ulva tessellata* Hook. f. et Harv., J. Bot. (Lond.) 4: 297, 1845 (as U. tesellata).

≡ *Prasiola tessellata* (Hook. f. et Harv.) Kütz., Species Algarum, p. 473, Leipzig, 1849.

≡ *Laestadia tessellata* (Hook.f. et Harv.) Har., Algues, p. 29 in Mission Scientifique du Cap Horn, 1882-1883, Vol. 5, Botanique, 1889.

= *Leptogiopsis complicatula* Nyl., Flora (Jena) 67: 211, 1884.

≡ *Turgidosculum complicatulum* (Nyl.) Kohlm. et E. Kohlm., Marine Mycology. The Higher Fungi, p. 361, Academic Press, New York 1979.

= *Physalospora prasiolae* Har., Journal Botanique de Paris 1:233, 1887 (nomen nudum).

= *Laestadia prasiolae* G. Winter, Hedwigia 26:16, 1887.

≡ *Guignardia prasiolae* (G. Winter) Lemmermann, Abhandlungen des Naturwissenschaftlichen Vereins Bremen 17: 199, 1901 [Note: *Guignardia prasiolae* (G. Winter) M. Reed 1902, superfluous new combination].

≡ *Plagiostoma prasiolae* (G. Winter) Clauzade, Diederich et Cl. Roux, Bulletin de la Société Linnéenne de Provence, Numéro spécial 1:47, 1989 (combinatio invalida).

=*Dermatomeris georgica* Reinsch, p. 425, in Internationale Polarforschung, Die deutschen Expeditionen und ihre Ergebnisse, Vol. II, 1890.

= *Guignardia alaskana* M. Reed, University of California, Berkeley, Publications, Botany 1:161, 1902.

≡ *Laestadia alaskana* (M. Reed) Sacc. et D. Sacc., in Saccardo, Sylloge Fungorum17: 576, 1905.

=*Mastodia mawsonii* Dodge, British and New Zealand Antarctic Research Expedition Report, Botany, 7: 57 (1948).

*Typus.* France, Kerguelen Islands, Christmas Harbour, June 1840, Anonimous *s. n.* (Herbario de Clifford, *BM ex K* 657).

The species is described and illustrated in Kolhmeyer *et al*. (2004).

#### Verrucaria durietzii

I.M. Lamb. Lilloa. 14: 205 (1948).; Fig. 3. — MycoBank MB#371531;

**Figure 3.**
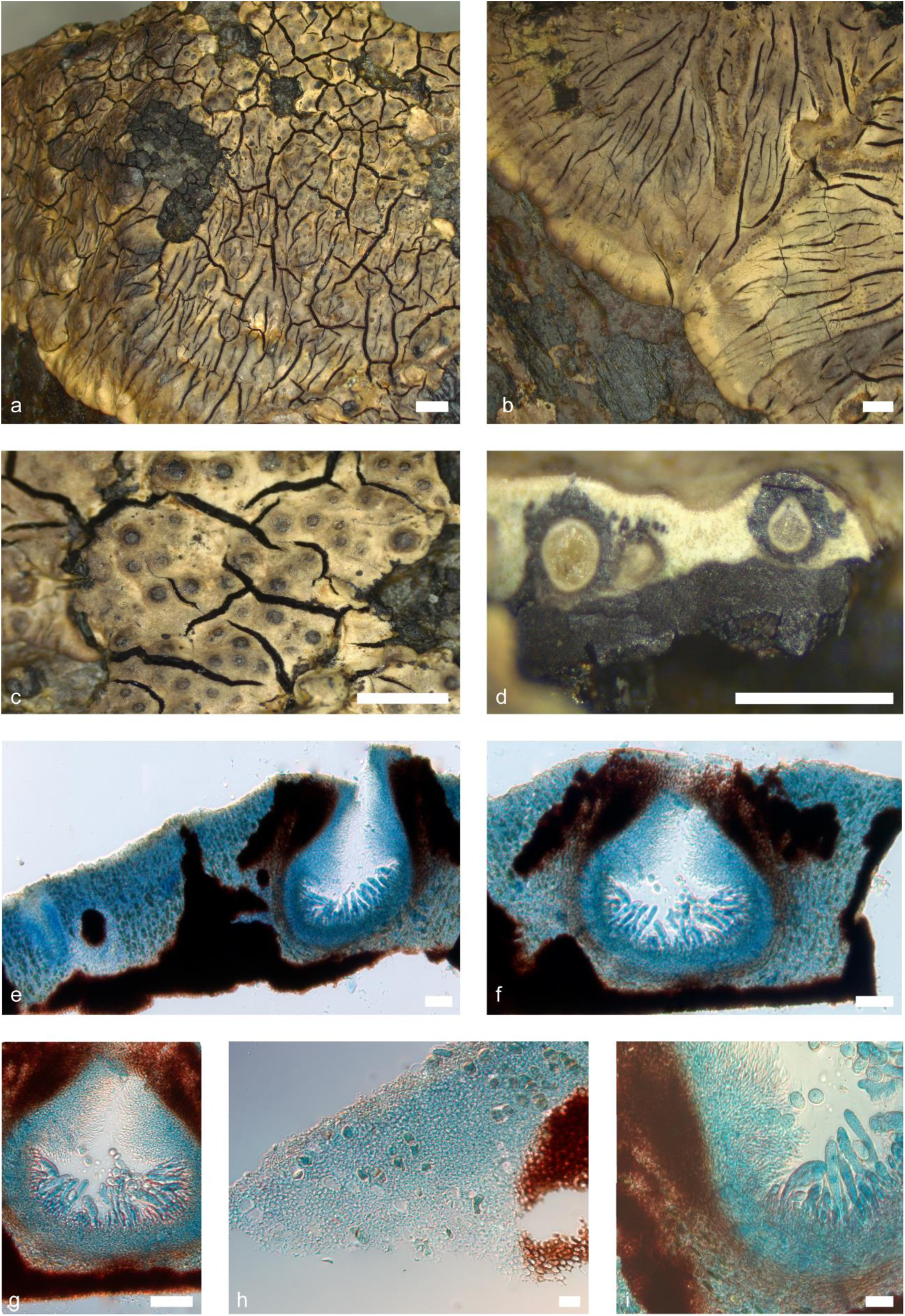
*Verrucaria durietzii* (Pérez-Ortega s.n.). Macroscopic and microscopic characters. a-b. Thallus habitus, c. Areoles bearing perithecia; d. Thallus section showing the basal carbonized layer and two perithecia section, e. Thallus section detail; f-g. Perithecium section detail; h. Structure thallus section, i. Periphyses, asci and excipulum. Scale bars: a, b: 2 mm, c, d: 1 mm, e, f, g, h, i: 50 µm. e-i: Lactophenol cotton blue. g-i: DIC.

= *Verrucaria durietzii* f. *rhabdota* I.M. Lamb. Lilloa 14: 206 (1948) [MB#479929].

*Typus.* Subantarctc Islands of New Zealand, Auckland Islands, Port Ross, boulder shore on the interior side of the innermost small peninsula in Laurie Harbour, dominant just above the pure Verrucaria maura-belt, 27 March 1927, *G. E. Du Rietz*, s.n. (Swedish Museum of Natural History Department of Botany (S), S-L125).

*Thallus* epilithic, crustose, continuous to areolate, usually effigurate up to 10 cm in diam, thick, 1000–1500(–2000) µm (*n*=18, *s*=12) in height; from white to pale cream or straw-yellowish in colour, with dark brownish shades visible towards the centre of the thallus and at the edges of the lobes. Areole surface flat, delimited by black margins, from 0.9 to 2.3 mm in diam. Hypothallus not present. Surface smooth, with long fissures with black margins. Thallus in section paraplectenchymatous, with two clear layers, mycobiont cells irregularly globose, 2–4(–5) µm (*n*=15, *s*=12) in diam; upper layer, 85–360 µm thick (*n*=14, *s*=12), consists of hyaline mycobiont cells and contain the photobionts cells, lower layer, 400–750(–900) µm thick (*n*=14, *s*=12) occasionally protruding towards the thallus surface, consists of carbonised melanised mycobiont cells. Phaenocortex present, 10–15(–16) µm thick (*n*=13, *s*=12). Pigmented brown layer between the Phaenocortex and the upper photobiont cells present, K-. Photobiont cells ±randomly arranged to more rarely distributed in columns, irregularly ellipsoidal to globose, (7–)9–12(–14) × (4–)5–7(–8) µm (*n*=20, *s*=12).

*Perithecia* immersed to semi-immersed up to 1/4, glossy black in colour, sometimes, when the thallus covers the involucrellum, it becomes matt, usually in groups of up to three perithecia; ellipsoidal to pyriform in section, (245–)252–460(–470) × (155–)162–300(–310) µm (*n*=14, *s*=12); ostiolar region non-papillate, typically paler, ostiolar opening visible at 10×. Invollucrellum present, (30–)45–70(–80) µm thick (*n*=13, *s*=12), typically extending up to the melanised basal layer. Excipulum hyaline, but it may darken slightly in the upper and lower parts, (10–)15–25 µm thick (*n*=15, *s*=12), composed of c. 11 layers of flattened rectangular cells. Periphyses present, septate, simple, 15–20 × 1–2 µm (*n*=17, *s*=12). Paraphyses are absent. Hamathecial gelatine: K-, I+ red, K/I+ blue. Asci bitunicate, exotunica disappears rapidly with maturity, clavate, 8-spored, occurring at the lower and lateral parts of the ascoma, (40–)50–80(–85) × (9–)10–14(–17) µm (*n*=14, *s*=11). *Ascospores* simple, hyaline, ellipsoidal to oval in shape, (11–)12–15(–17) × (6–)7–9 µm (*n*=35, *s*=12), without halonate perispore.

*Pycnidia* typically present, immersed, randomly distributed on the thallus, sometimes forming groups of up to four; narrowly ellipsoid in section, (90–)120–210(–215) × (35–)45–87(–90) µm (*n*=10, *s*=9), pycnidial wall hyaline. Conidiospores bacillar, hyaline, 4–6 × 1 µm (*n*=18, *s*=9).

Ecology & Distribution – *V. dutietzii* is a marine lichen, not exactly amphibious, but characteristic of the zone just above high tide level (lower hygrohaline), where it’s exposed to the salt spray from wave action, on coastal rocks, often associated with Candelariella, Caloplaca, Rinodina, *Verrucaria* sp. and *V. tessellatula*. (Labm 1948b; Galloway 2007). *V. durietzii* is geographically restricted to the Southern Hemisphere and the circumpolar Antarctic region. The species is found in southern Chile, the Magallanes Region, and Tierra del Fuego, as well as in the sub-Antarctic Islands, including the Falkland Islands, Kerguelen, South Georgia, Marion and Prince Edward Islands, and Macquarie Island (Lamb 1948b; McCarthy 1991; Øvstedal & Gremmen 2001; Øvstedal & Lewis Smith 2001). It is also known from New Zealand (Galloway 2007).

Notes – The species *V. durietzii* can be readily distinguished from other members of the genus by the distinctive continuous areolate thallus, which can reach a diameter of up to 10 cm, and the characteristic creamy brown coloration with yellowish tones. However, it also exhibits a typically thick thallus, measuring up to 2 mm in thickness. Nevertheless, upon microscopic examination, the perithecia may appear similar to those of other species, such as *V.* sp. 15 or even, in less developed perithecia, to those of *V*. sp. 16. However, this group of species, along with *V*. sp. 13, forms a clade, as illustrated in Fig. 1. However, a clear difference in size is observed when the mature perithecia of the three are compared. *V. durietzii* has the largest perithecia (252-460 × 162-300 µ). m) In comparison to *V*. sp. 12 (150-200 × 125-180 µm) and *V*. sp. 14 (170-240 × 150-220 µm). However, the involucrellum of the three do have slightly similar sizes: 45-70 µm for *V. durietzii*, 60-80 µm for *V*. sp. 12, and 50-60 µm for *V*. sp. 14.

Conversely, the forma rhabdobata, described from East Falkland Islands (Berkeley, Port Louis, 1946, I. M. Lamb 2939) is characterized by the presence of black striae (juga) approximately 80 µm wide, parallel to the thallus cracks. In our opinion, this character lacks taxonomic relevance., located at some distance from the centre.

##### Additional specimens examined

Chile, Magallanes and Antarctica Region, Navarino Island, Cove near Puerto Navarino in front of Hoste Island, 54°55’48’S, 68°20’45’W, on rock. 0-15 m alt. 27 January 2008. *S. Pérez-Ortega* s. n. (MA-Lich) – Chile, Isla Grande de Tierra de Fuego, Southwest arm of Seno Parry, Blocks near the beach, 54°40’32’S, 69°26’25’W, on rock. 0-15 m alt. 10 December 2009. *S. Pérez-Ortega* s. n. (MA-Lich) – Chile, Magallanes and Chilean Antarctica Region, Isla Grande de Tierra del Fuego, west arm of the Pía glacier, 54°42’36’S, 69°42’01’W, on rock. 0-5 m alt. 16 December 2009. *S. Pérez-Ortega s. n.* (MA-Lich) – Chile, Magallanes and Chilean Antarctica Region, Tierra del Fuego, Beagle Channel, Chair Island, Darwin Bay, 54°53’59’S, 70°00’48’W, on rock. 0-5 m alt. 16 December 2009. *S. Pérez-Ortega s. n.* (MA-Lich) – Chile, Magallanes and Chilean Antarctica Region, Tierra del Fuego, Beagle Channel, Chair Island, 54°53’52’S, 70°2’12’W, on rock. 0-5 m alt. 16 December 2009. *S. Pérez-Ortega s. n.* (MA-Lich) – Chile, Magallanes and Chilean Antarctica Region, Tierra del Fuego. Isla Basket. Intermareal, 54°42’13’S, 71°34’53’W, on rock. 0-1 m alt. 17 December 2009. *S. Pérez-Ortega s. n.* (MA-Lich) – West Falkland, Roy Cove, 51°32’05’S 60°23’05’W, on rocks. 15 m alt. 17 January 2011. *A. Orange* 19927 (Welsh National Herbarium NMW) – French Southern and Antarctic Lands, Kerguelen, Ile Foch, along SE side of Baie Phillips, 48°57’00’S, 69°19’6’E, on basaltic rock. 0-1 alt. 22 February 1999. *I.M. Lamb* CL 76756 (Museum Botanicum Hauniense C-L-76756).

#### Verrucaria psychrophila

M. Lamb, Discovery Reports 25: 18 (1948); Fig. 4. — MycoBank MB 371775

**Figure 4.**
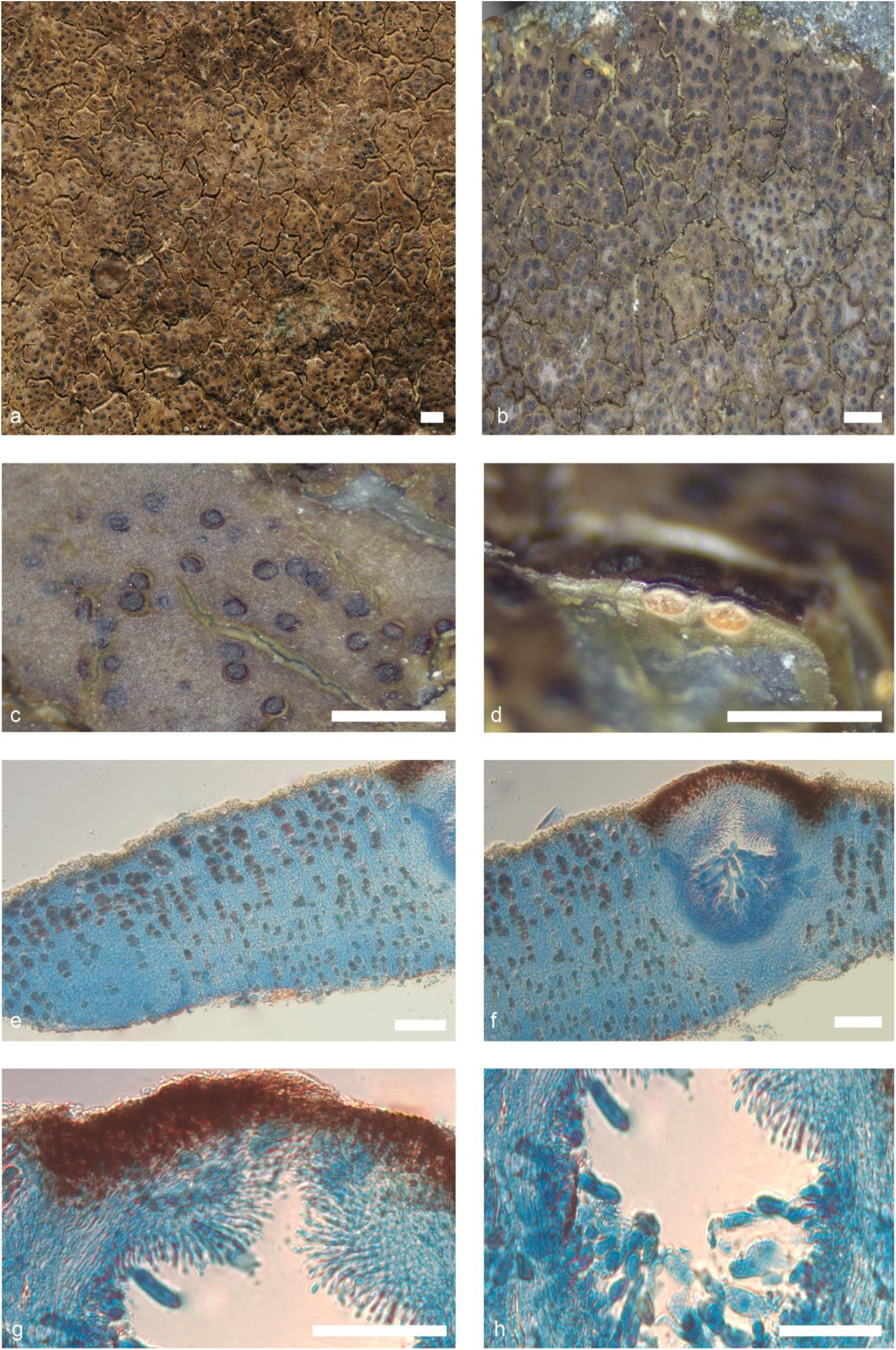
*Verrucaria psychrophila* (de Los Ríos *s. n.*). Habitus and anatomical features. a-b. Thallus habitus, c. Detail of perithecia; d. Thallus section showing perithecia, e. Thallus section; f. Thallus section showing a perithecium; g. Detail of the ostiolar region showing the periphyses detail; h. Asci and excipulum. Scale bars: a: 2 mm, b, c, d: 1 mm, e, f, g, h: 50 µm. e-h: Lactophenol blue. g-h: DIC.

*Typus.* Falkland Islands (Malvinas), W. Graham Land, Palmer Archipelago, Port Lockroy, Goudier Islet, on granodiorite rocks, 25 March 1944, Anon., s.n. Natural History Museum (BM), BM001107134!.

*Thallus* epilithic, crustose, rimose, up to 4 cm in diam, and 200–250 µm (*n*=12, *s*=6) in height; from dark brownish-brown to light brown to greyish shades, slightly shiny, waxy appearance. Thallus surface smooth, some specimens heavily fissured, forming irregular polygonal areoles of 1.5 to 3.1 mm. Areole surface flat to slightly convex, bordered by concoloured margins. Hypothallus not present. Punctae or ridges not present. Thallus in section paraplectenchymatous. Mycobiont cells irregularly globose to polygonal, up to 4–5 × 3 µm (*n*=35, *s*=6). A translucent refringent layer with cells with lipid content, basal layer is present, KI-. Phaenocortex absent, sometimes a thin, discontinuous layer 5–10 µm thick is visible (*n*=4, *s*=3). Brown pigment layer above photobiont layer present, K-. Photobiont cells arranged in vertical columns, irregularly ellipsoidal to globose, (8–)10–15(–16) × (5–)6–10(–12) µm (*n*=24, *s*=6).

*Perithecia* immersed, groups of up to three per areole, shiny blackish in colour, globose to slightly pyriform in shape, (160–)170–220(–225) × (145–)150–210(–215) µm (*n*=12, *s*=6); ostiolar region non-papillate, typically concolour, ostiolar opening visible at 10×. Involucrellum present, 20–30 µm thick (*n*=12, *s*=6), extending to the upper third of the perithecium. Excipulum hyaline, 15–20 µm thick (*n*=8, *s*=6), composed of c. 7 layers of flattened rectangular cells. Periphyses present, septate, simple, 15–20(–30) × 1–2 µm (*n*=8, *s*=6). Paraphyses are absent. Hamathecial gelatine: K-, I+ red, K/I+ blue. Asci bitunicate, exotunica disappears rapidly with maturity, clavate, 8-spored, located in the lower and lateral parts of the perithecial cavity, (35–)40–50(–53) × (7–)10–14(–15) µm (*n*=17, *s*=6). *Ascospores* simple, hyaline, ellipsoidal to slightly oval in shape, (9–)10–12 × 6–9 µm (*n*=33, *s*=6), without halonate perispore.

*Pycnidia* common, immersed, randomly distributed in the thallus, light brown spots, usually unilocular, but multilocular when very mature, (80–)90–120(–125) × (45–)50–80(–85) µm (*n*=8, *s*= 4), pycnidial wall hyaline. Conidiospores bacillar, hyaline, 4–5 × 1(–2) µm (*n*=22, *s*=3).

Ecology & Distribution – *V. psichrophila* is a marine lichen, amphibious, intermittently submerged by the tide, occurring at low level in the littoral zone (Lamb 1948b). On coastal rocks, sometimes associated with other species of *Verrucaria* like *V*. gr. *dispartita* or *V*. sp. 20. *V. psichrophila* is geographically restricted to the Southern Hemisphere, with a particularly notable presence in maritime Antarctica and the sub-Antarctic islands. The species is predominantly found on Livingston Island, the Palmer Archipelago, and Gould Island (West Graham Land) (Dodge 1973; Lamb 1948a).

Notes – *V. psychrophila* was formerly confused with *V. ceuthocarpa* from the Northern Hemisphere, and even with some specimens of *V. tessellatula* (Lamb 1948a). However, this study shows that northern hemisphere species such as *V. ceuthocarpa* are not present in the southern hemisphere (as shown in Fig. 1). However, it may be that the species Lamb (1948a) cites as *V. ceuthocarpa* from the southern hemisphere may actually be referring to species such as *V*. sp 19. which would be very similar to *V. ceuthocarpa* (H.N) and *V. psychrophila*. A phylogenetic analysis reveals that the sister species of *V. psychrophila* is *V*. sp. 21. However, this species can be readily distinguished from *V*. psychrophila due to the presence of distinct black cracks and a magenta-blue KI+ reaction in the basal layer, which can extend up to the middle of the thallus. In contrast, *V. psychrophila* lacks this reaction and exhibits craks that are similar in colour to the surrounding thallus. Conversely, *V*. sp. 19 exhibits considerable morphological similarity to *V. psychrophila*, displaying brown and occasionally greyish tones and perithecia immersed in the thallus. However, in the case of *V*. sp. 19, the craks are also darker than the rest of the thallus, reaching a black colour and exhibiting slightly smaller perithecia (140-185 × 130-160 µm) than *V. psychrophila* (170-220 × 150-210 µm).

##### Additional specimens examined

Antarctica, Livingston Island, Caleta Española, 62°39’24’S, 60°21’57’W, on rock. 0-5 m alt. 21 January 2014. *A. de los Ríos* 2.3 – Antarctica, Livingston Island, Sally Rocks, 62°42’07’S 60°25’44’W, on rock. 0-5 m alt. 25 January 2014. *A. de los Ríos s.n.* – Antarctica, Livingston Island, Punta Polaca, 62°39’08’S 60°22’16’W, on rock. 0-5 m alt. 30 January 2014. *A. de los Ríos s.n.* – Antarctica, Livingston Island, Caleta Argentina, 62°39’57’S 60°23’49’W, on rock. 0-5 m alt. 11 March 2019. *A. de los Ríos s.n.* – Antarctica, Livingston Island, Punta Barnard, 62°45’27’S 60°20’20’W, on rock. 0-5 m alt. 20 August 2023. *A. de los Ríos 075* – Chile, Magallanes and Chilean Antarctica Region, Holger Islet. Rocky seashores, intertidal zone., 54°56’28’S, 67°15’03’W, on rock. 0-5 m alt. 22 January 2008. *S. Pérez-Ortega s. n.* (MA-Lich).

#### Verrucaria serpuloides

M. Lamb, Discovery Reports 25: 18 (1948); Fig. 5. — MycoBank MB#371820

**Figure 5.**
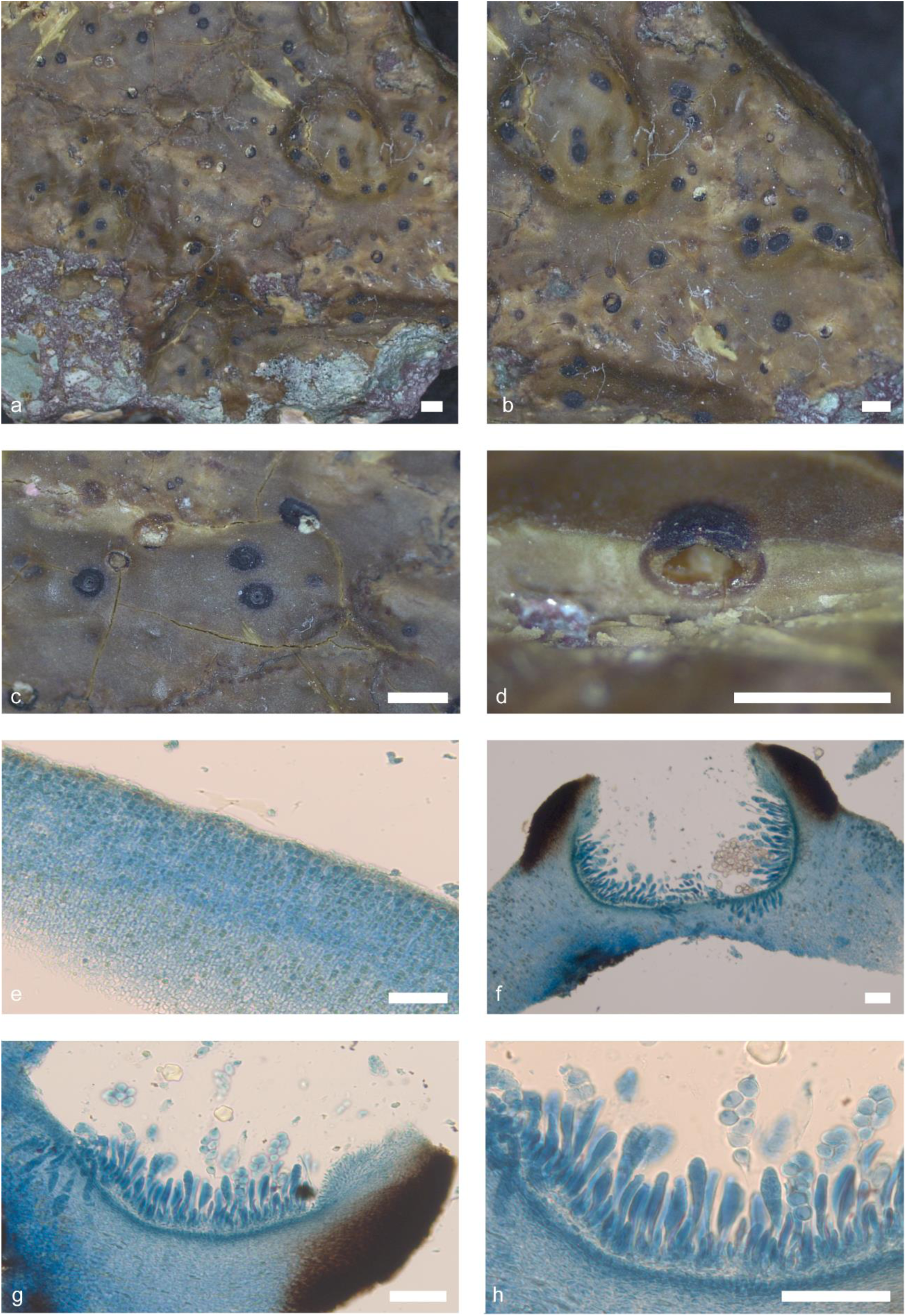
*Verrucaria serpuloides* (A. Peters *s. n*.). Thallus habitus and microscopic features. a-b. Thallus habitus, c. Detail of perithecia; d. Section of the thallus showing perithecia, e. Thallus section with a visible brown layer; f. Thallus in section showing perithecia; g. Periphyses detail; h. Perithecium, i. Asci and excipulum. Scale bars: a, b: 2 mm, c, d: 1 mm, e, f, g, h, i: 50 µm. e-i: Lactophenol cotton blue. g-i: DIC.

*Typus.* Falkland Islands (Malvinas), E. Graham Land, trinity Peninsula, Hope Bay, on granodiorite rocks, below the level of the lowest spring tides and therefore permanently submerged, 21 November 1945, Anon. 2565 Natural History Museum (BM), BM001107134!.

*Thallus* epilithic, crustose, continuous, up to 4 cm in diam, thick, 300–350 µm (*n*=6, *s*=1) in height; from dark olive green to yellowish-brown ochre, occasionally olive black, matt, waxy appearance. Thallus surface smooth, sometimes with fissures forming flat irregular areoles. Hypothallus not present. Punctae or ridges not present. Thallus in section paraplectenchymatous. Mycobiont cells from irregularly globose to slightly rectangular, 4–7 × 3–6 µm (*n*=8, *s*=1). A translucent refringent layer with cells with lipid content, basal layer is present, KI-. Phaenocortex absent. Brown pigment layer above photobiont layer present, K-. Photobiont cells arranged in vertical columns, from irregularly ellipsoidal to globose, (10–)13–15(–16) × (4–)5–7(–9) µm (*n*=5, *s*=1).

*Perithecia* immersed, slightly protruding with age, dull blackish in colour, globose to pyriform in shape, (300–)320– 340(–350) × (265–)270–290(–295) µm (*n*=5, *s*=1); ostiolar region non-papillate, typically concolour with the rest of perithecia, ostiolar opening visible at 10×. Involucrellum present, 50–60 µm thick (*n*=6, *s*=1), extending to the upper third of the perithecium. Excipulum hyaline, 10–12 µm thick (*n*=3, *s*=1) composed of c. 5 layers of flattened rectangular cells. Periphyses present, septate, simple, 30–35(–45) × 1–2 µm (*n*=5, *s*=1); Paraphyses absent. Hamathecial gelatine: K-, I+ red, K/I+ blue. Asci bitunicate, exotunica disappears rapidly with maturity, clavate, 8-spored, located in the lower and lateral parts of the perithecial cavity, (35–)45–60(–65) × (7–)8–12(–14) µm (*n*=6, *s*=1). *Ascospores* simple, hyaline, ellipsoid to slightly oval, (13–)14–15 × 8–9(–10) µm (*n*=23, *s*=1), without halonate perispore.

*Pycnidia* common, immersed, randomly distributed on the thallus, hardly visible, ostiole concolour with the thallus, narrowly ellipsoid, unilocular, (130–)140–150(–155) × (55–)60–70(–80) µm (*n*=4, *s*=1), pycnidial wall hyaline. Conidiospores bacillar, hyaline, 5–7 × 1 (–2) µm (*n*=15, *s*=1).

Ecology & Distribution – *V. serpuloides* is a marine lichen that is found in a permanently submerged state below the level of the lowest spring tides. It is sometimes found on rocks, in association with other species of encrusting calcareous algae (Lamb 1948a). *V. serpuloides* is geographically restricted to the Southern Hemisphere, in maritime Antarctica (the Palmer Archipelago and Goudier Islet), although it is also known from New Zealand (Galloway 2007; Lamb 1948a).

Notes – Previously*, V. serpuloides* was considered to be a potential misidentification of some specimens of *W. mucosa* (Galloway 2007; Lamb 1948a). However, it has been demonstrated that the species *W. mucosa* is not present in the southern hemisphere (Fig. 1), suggesting that the specimens in question may actually represent V. sp. 15 or 16, which are species that are present in this hemisphere. An examination of the phylogeny (Fig. 1) reveals that V. serpuloides forms a clade with two species with which it can be morphologically assimilated, namely V. sp. 15 and 16. The perithecia of *V. serpuloides* are slightly larger than those of *V*. sp. 15 and 16, measuring 320-340 × 270-290 µm and 200-300 × 250-300 µm, respectively. However, in consideration of the species’ ecology, *V. serpuloides* is the sole species capable of complete and permanent submergence in marine environments at depths of 4 to 10 meters below mean low tide level (Lamb 1970, 1973).

##### Additional specimens examined

Antartica, on rocks. 16 December 1997. A. Peters *s. n*.

#### Verrucaria tessellatula

Nyl., J. Bot., Lond. 13: 335 (1875); Fig. 6. — MycoBank MB 409864

**Figure 6.**
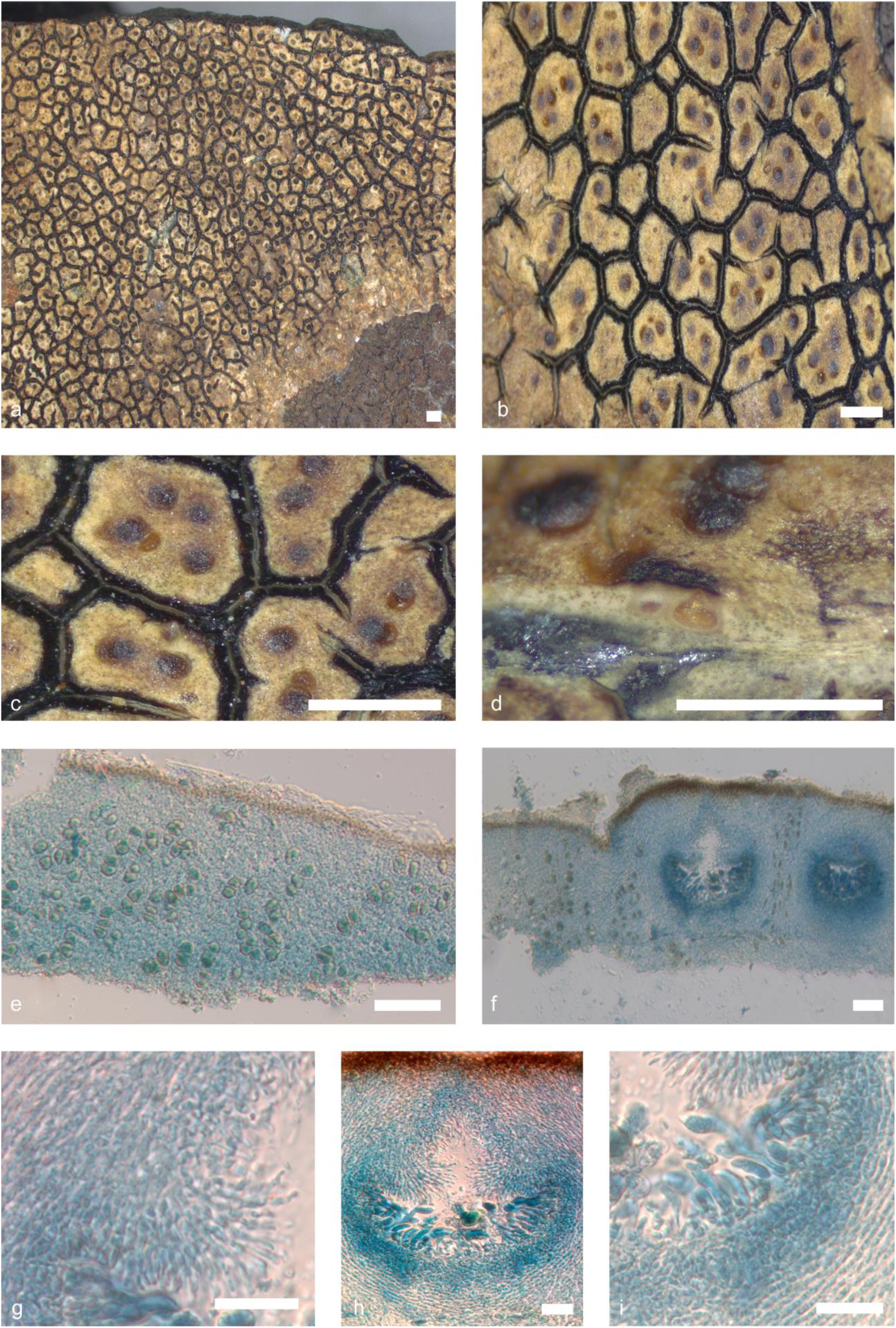
*Verrucaria tessellatula* (num.: Pérez-Ortega *s. n.*). Thallus habitus and microscopic features. a-b. Thallus habitus, c. Detail of perithecia in the areoles; d. Section of the thallus showing perithecia, e. Thallus section with a visible brown layer; f. Thallus in section showing perithecia; g. Periphyses detail; h. Perithecium, i. Asci and excipulum. Scale bars: a, b: 2 mm, c, d: 1 mm, e, f, g, h, i: 50 µm. e-i: Lactophenol cotton blue. g-i: DIC.

= *Verrucaria tessellatula* f. *dermoplaca* (Nyl.) I.M. Lamb, Discovery Reports 25: 22 (1948) [MB#480860].

*Typus.* French Southern Territories, Swan’s Bay, Kerguelen Land, on rocks, January 1875, A. E. Eaton *s. n.* (Natural History Museum (BM), BM001107160).

*Thallus* crustose, epilithic, areolate, up to 6 cm in diam, thick, 150–280(–300) µm (*n*=20, *s*=17) in height; from pale brown to yellowish cream, occasionally slightly dark brownish or even slightly maroon, matt. Thallus surface smooth to slightly rough, with strong fissures towards the centre, forming irregular polygonal areoles, from 0.9 to 1.9(–3.5) mm in diam. Areole surface flat to slightly convex, usually delimited by black margins that often slightly protrude from the surface of the aureole. Hypothallus usually present, whitish to cream-coloured. Punctae or ridges not present. Thallus in section paraplectenchymatous. Mycobiont cells irregularly globose, up to 4–5 µm in diameter (*n*=35, *s*=17). A translucent refringent layer with cells with lipid content, basal layer is present, KI-. Phaenocortex present, (12–) 15–25 (–30) µm thick (*n*=30, *s*=17). Brown pigment layer present, between Phaenocortex and photobiont layer, K-. Photobiont randomly arranged throughout the thallus, although photobiont columns can be occasionally observed. Photobiont cells irregularly ellipsoidal to globose, (10–)13–20(–22) × (4–)5–15(–17) µm (*n*=38, *s*=17).

*Perithecia* immersed to semi-immersed up to 1/4, from 1 to 8 per areole, glossy black in colour, irregularly globose to oval in shape, (130–)140–275(–285) × (130–)140–250(–260) µm (*n*=22, *s*=16); ostiolar region typically concolour, ostiolar opening visible at 10×. Involucrellum present, (12–)15–25(–27) µm thick (*n*=18, *s*=16), extending to the upper third of the perithecium. Excipulum hyaline, (13–)15–25(–30) µm thick (*n*=20, *s*=17), composed of c. 8 layers of flattened rectangular cells. Periphyses present, septate, simple, 10–15 (–20) × 2–3 µm (*n*=17, *s*=16). Paraphyses are absent. Hamathecial gelatine K-, I + red, K/I + blue. Asci bitunicate, exotunica disappears rapidly with maturity, clavate, 8-spored, located in the lower and lateral parts of the perithecial cavity, (30–)35–56(–60) × (6–)8–17(–19) µm (*n*=19, *s*=17). *Ascospores* simple, hyaline, ellipsoidal to ovoid, (7–)10–15 × 6-8(–10) µm (*n*=45, *s*=16), without halonate perispore.

*Pycnidia* common, immersed, irregularly distributed on the thallus, with clusters occurring only in certain areas, although more mature ones may become multilocular; irregularly globose to narrowly ellipsoid, (90–)100–180(–190) × (35–)45–87(–92) µm (*n*=14, *s*=12). Conidiospores bacillar, hyaline, 4–7 × 1–2 µm (*n*=19, *s*=12).

Ecology & Distribution – *V. tessellatula* is not strictly amphibious, but is characteristic of the area just above high tide level, where it is exposed to salt spray from wave action on coastal rocks. It is often found in association with other *Verrucaria* species, *Hydropunctaria maura* or *Caloplaca* sp. (Galloway 2007; Lamb 1948a). The species is geographically restricted to the Southern Hemisphere, occurring in maritime Antarctica and the sub-Antarctic islands, southern South America, and New Zealand. It is found in Tierra del Fuego, the Magellan Region, Livingston Island, Kerguelen (type locality), the Falkland Islands, South Orkney Island, South Shetland Island, South Georgia, Marion and Prince Edward Islands, among others (Galloway 2007).

Notes – The species *V. tessellatula* is readily identifiable, characterised by a brown areolate crustose thallus with yellowish or cream tones and deep black cracks. This distinctive combination of traits renders it a rather striking species. However, in some specimens that are not yet fully developed, there may be some confusion with *V. psychrophila*. However, the latter has the cracks of the thallus that are light brown or the same colour as the rest of the thallus, which is distinct from the colour of *V. tessellatula*. In some cases, thallus is not so areolate and have a more continuous appearance, separated by some cracks, giving rise to irregular areas of variable size. This type of thallus would correspond to *V. tessellatula* f.*dermoplaca* Nyl. From a phylogenetic perspective, this species is closely related to *Mastodia tessellata* (Fig. 1), which exhibits a markedly different morphology. The latter possesses a foliaceous thallus due to its association with the green foliaceous alga of the genus *Prasiola*. The formation of this clade by these two species gives rise to a debate concerning the genus *Mastodia* and the question of whether the entire clade constituted by the *Verrucaria* of the southern hemisphere should be considered as a genus *Mastodia*, or whether new genera should be created.

##### Additional specimens examined

Antartida, Livingston Island, Caleta Española, 62°39’24’S, 60°21’57’W, on rock. 0-5 m alt. 21 January 2014. *A. de los Ríos s.n.* – Antartida, Livingston Island, Sally Rocks, 62°42’07’S, 60°25’44’W, on rock. 0-5 m alt. 25 January 2014. *A. de los Ríos s.n.* – Antartida, Livingston Island, Punta Polaca, 62°39’08’S, 60°22’16’W, on rock. 0-5 m alt. 30 January 2014. *A. de los Ríos s.n.* – Antartida, Livingston Island, Punta Barnard, 62°45’27’S, 60°20’20’W, on rock. 0-5 m alt. 20 August 2023. *A. de los Ríos s.n.* – Falkland Islands, East Falkland, Darwin Harbour, 51°48’68’S, 58°57’60’W, on rock. 0-5 m alt. 13 January 2011. *A. Orange* 19769 (Welsh National Herbarium NMW) – Chile, Aysén Region, Aysén province, Chonos Archipelago, coast NNW of the village Puerto Aguirre, 73°31’53’W, 45°09’29’S, on coastal beadrocks. 0-5 m alt. 16 February 2016. *U. Schiefelbein* 4328 (Herbarium Musei Britannici) – Chile, Magallanes and Chilean Antarctic Region, Province of Magallanes, Punta Arenas, N of Puerto del Hambre, 70°57’33”W, 53°29’57”S, on sedimentary rock. 0-5 m alt. 8 January 2023. *U. Schiefelbein* 6476 – Chile, Magallanes and Chilean Antarctic Region, Province of Última Esperanza, Natales, Seno Última Esperanza, coast S of Puerto Natales, 72°29’05”W, 51°45’27”S, on sedimentary rock. 0-5 m alt. 17 January 2023. *U. Schiefelbein* 6565 – Chile, Magallanes and Chilean Antarctic Region, Picton Island, Intertidal, 54°59’29”S, 67°3’11”W, on rock. 1-5 m alt. 23 January 2008. *S. Pérez-Ortega s.n.* (MA-Lich) – Chile, Magallanes and Chilean Antarctic Region, Snook Bay, Marchant Island, intertidal zone, 54°56’3”S, 67°40’30”W, on rock. 0 m alt. 23 January 2008. *S. Pérez-Ortega s.n.* (MA-Lich) – Chile, Magallanes and Chilean Antarctic Region, Drymis winteri forest near Puerto Navarino, 54°55’57”S, 63°20’59”W, on rock. 27 m alt. 27 January 2008. *S. Pérez-Ortega s.n.* (MA-Lich) – Chile, Magallanes and Chilean Antarctic Region, Navarino Island, Cove near Puerto Navarino in front of Hoste Island, 54°55’48”S, 68°20’45”W, on rock. 0-15 m alt. 27 January 2008. *S. Pérez-Ortega s.n.* (MA-Lich) – Chile, Magallanes and Chilean Antarctic Region, Isla Grande de Tierra del Fuego, Seno del Almirantazgo, Gabriel’s Channel, 54°25’55”S, 70°7’48”W, on rock. 4 m alt. 12 December 2009. *S. Pérez-Ortega s.n.* (MA-Lich) – Chile, Magallanes and Chilean Antarctic Region, Isla Grande de Tierra del Fuego, Seno d’Agostini, Angelitos Cove, Peat bogs, 54°24’12”S, 70°26’24”W, on rock. 20 m alt. 13 December 2009. *S. Pérez-Ortega s.n.* (MA-Lich) – Chile, Magallanes and Chilean Antarctic Region, Tierra del Fuego, Basket Island, 54°44’25”S, 71°34’31”W, on rock. 0-5 m alt. 17 December 2009. *S. Pérez-Ortega s.n.* (MA-Lich).

#### Verrucaria pseudodispartita

Fernández-Costas & Pérez-Ort.; Fig. 7. sp. nov.— (**sp. 1**)

**Figure 7.**
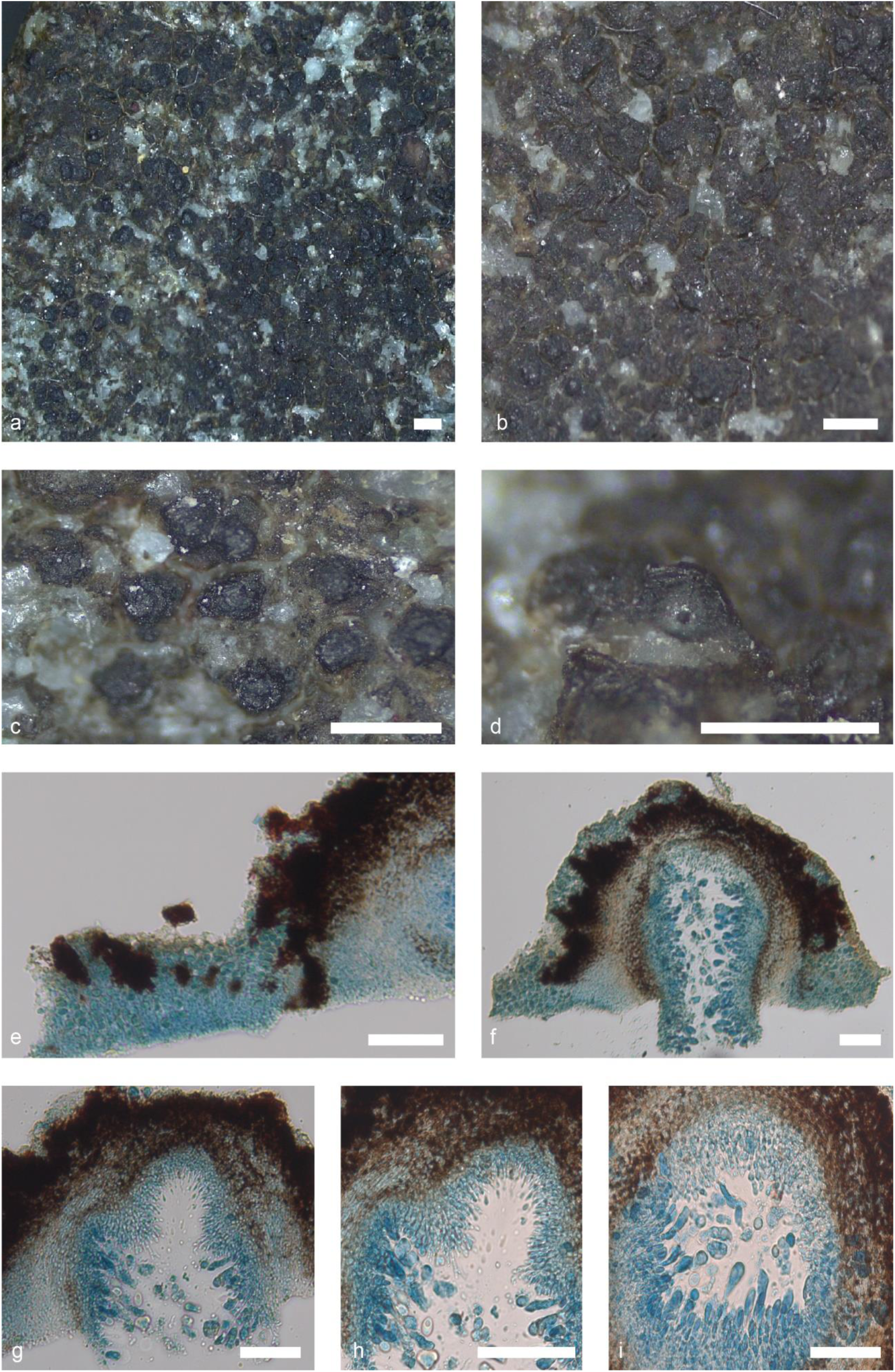
*Verrucaria pseudodispartita* (De Los Ríos *s. n.*). Thallus habitus and microscopic features. a-b. Habitus, c. Detail of the perithecia; d. Section of a perithecium, e. Thallus section showing the punctae; f. Thallus section with perithecia; g, h, i. Perithecia section detail. Scale bars: a, b, c, d: 1 mm, e, f, g, h, i: 50 µm. e-i: Lactophenol cotton blue. g-i: DIC.

*Etymology.* The specific epithet ‘*pseudodispartita*’, refers to the fact that it is very similar to *V. dispartita*.

*Typus.* Antartida, Livingston Island, Punta Polaca, 62°39’08’S 60°22’16’W, on rock. 0-5 m alt. 30 January 2014*. A. de los Ríos s. n.* (MA-Lichen).

*Thallus* epilithic, crustose, areolate, tiny, up to 0.4 cm in diam, and 60–80 µm in height (*n*=6, *s*=2); from black to olive green and brownish brown, shiny. Thallus surface rough, cracks forming polygonal areoles, 0.4 × 1.1 mm. Surface flat to slightly convex, usually bordered concolour with the rest of the thallus, punctae present, numerous, throughout thallus, giving it a rough appearance, 25–50 × 10–40 µm. Hypothallus not present. Thallus in section paraplectenchymatous. Mycobiont cells from irregularly globose to polygonal, 4–5 × 2–3 µm (*n*=12, *s*=2). A translucent refringent layer absent. Phaenocortex absent. A brownish pigment layer, K-, is located above photobiont layer. Photobiont randomly arranged throughout the thallus, from irregularly ellipsoidal to globose, (7–)8–10(–11) × (4–)5–9(–10) µm (*n*=16, *s*=2).

*Perithecia* sessile, shiny black colour, irregularly oval in shape. In section, from globose-ellipsoidal to pyriform, (290–)200–250(–255) × (150–)200–275(–280) µm (*n*=8, *s*=2); ostiolar region typically concolour with the rest of perithecia, ostiolar opening barely visible at 10×. Involucrellum present, scabrid, (55–)60–70(–75) µm thick (*n*=8, *s*=2), extending up to the perithecial base. Excipulum from light to drak brown, 10–15(–20) µm thick (*n*=9, *s*=2), composed of 7–8 layers aprox. of flattened rectangular cells. Periphyses present, septate, simple, (9–)10–13 × 1–1.5. Paraphyses are absent. Hamathecial gelatine K-, I + red, K/I + blue. Asci bitunicate, exotunica disappears rapidly with maturity, clavate, 8-spored, located in the lower and lateral parts of the perithecial cavity, (25–)30–45(–50) × 10–15 µm (*n*=10, *s*=2). *Ascospores* simple, hyaline, from ellipsoidal to ovoid, (7.5–)8–10(–10,5) × (4–)5–6(–6.5) µm (*n*=40, *s*=2).

*Pycnidia* present, immersed, irregularly distributed on the thallus, inconspicuous, globose, (40–)50–60(–65) × (35–)40–50(–60) µm (*n*=3, *s*=1), pycnidial wall hyaline. Conidiospores not observed.

Ecology & Distribution – *V*. sp.1 is a marine lichen, amphibious, intermittently submerged by the tide, occurring at low level in the littoral zone. On coastal rocks, sometimes associated with other species of *Verrucaria* like *V*. psychrophila or *V*. sp. 19. This species is geographically restricted to the southern hemisphere, in particular it has only been found in marine Antarctica.

Notes – *V*. sp. 1 is distinguished by its diminutive, strongly areolate thallus and diminutive perithecia with a scabrous involucrellum. However, this species is readily mistaken for *V*. sp. 3, *V*. sp. 2, or *V*. sp. 7. *V*. sp. 2 and 3 form a clade together with *V*. sp. 1 (Fig. 1). These species exhibit significant morphological similarities, forming a distinctive group that could be designated as gr. dispartito. However, it should be noted that perithecia of *V*. sp. 2 and 3 are smaller (120-180 × 110-140 µm) than those of *V*. sp. 1 (200-250 × 200-275 µm). Furthermore, the spores of *V*. sp. 1 (8-10 × 5-6 µm) are slightly larger than those of *V*. sp. 2 and 3 (8-9 × 4-5 µm). In contrast, *V*. sp. 7 exhibits perithecia (235-250 × 180-200 µm) and ascospores (8-10 × 5-6 µm) that are more similar in size to those of *V*. sp. 1 (perithecia (200-250 × 200-275 µm), ascospores (8-10 × 5-6 µm)). However, *V*. sp. 7 exhibits a markedly rougher thallus, characterized by a greater number of more protruding punctae that may even form ridges. In contrast to *V*. sp. 1, which exhibits a rougher thallus with a smaller number of punctae that do not protrude from the thallus in such an erumpent manner. Furthermore, the photobiont cells of *V*. sp. 1 are observed to be slightly larger (8-10 × 5-9 µm) in comparison to those of *V*. sp. 7 (5-8 × 4-6 µm).

##### Additional specimens examined

Antartida, Livingston Island, Punta Polaca, 62°39’07’S 60°22’14’W, on rock. 0-5 m alt. 30 January 2014. *A. de los Ríos*.

#### Verrucaria cuncorum

Fernández-Costas & Pérez-Ort.; Fig. 8. sp. nov. — (**sp. 2**)

**Figure 8.**
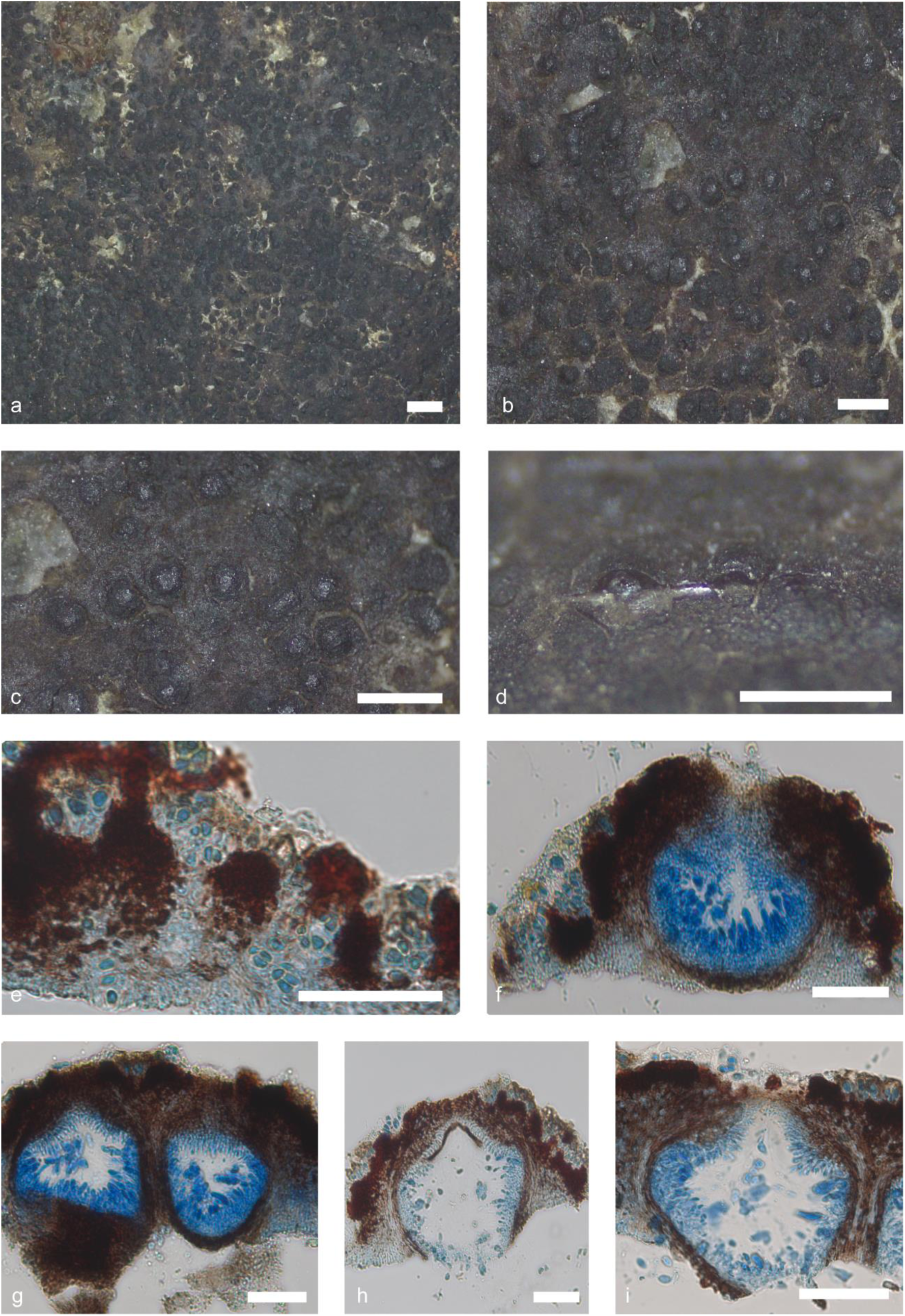
*Verrucaria sp.* 2 (num.: Pérez-Ortega *s. n.*). Thallus habitus and microscopic features. a-b. Habitus, c. Detail of the perithecia; d. Section of a perithecium, e. Thallus section showing the punctae; f. Thallus section with perithecia; g, h, i. Perithecia section detail. Scale bars: a, b, c, d: 1 mm, e, f, g, h, i: 50 µm. e-i: Lactophenol cotton blue. g-i: DIC.

*Etymology.* The specific epithet ‘*cuncorum*’, refer to the poorly known subgroup of Huilliche people native to coastal areas of southern Chile.

*Typus.* Chile, Lakes Region, Achao, Chiloé Island. Playa de Achao, rocks on the supralittoral, 42°28’9”S, 73°29’19”W, on rock. 0-5 m alt. 3 December 2009. *S. Pérez-Ortega s. n.* (MA-Lich).

*Thallus* epilithic, crustose, continuous, up to 3.9 cm in diam, and 25–30 µm in height (*n*=5, *s*=1); from black to blackish brown, shiny. Thallus surface rough. Surface flat to slightly convex; punctae present, numerous, throughout thallus, giving it a rough appearance, 10–20 × 20–30 µm. Hypothallus not present. Thallus in section paraplectenchymatous. Mycobiont cells from irregularly globose to polygonal, 3–5 × 2–3 µm (*n*=10, *s*=1). A translucent refringent layer absent. Phaenocortex absent. A blackish brown pigment layer, K-, is located above photobiont layer. Photobiont randomly arranged throughout the thallus, from irregularly ellipsoidal to globose, (4–)5–8(–9) × (3–)4–5(–6) µm (*n*=10, *s*=1).

*Perithecia* sessile to semi-immersed up to 2/3, shiny black colour, from irregular pyriform to oval in shape with flattened tip. In section, from ellipsoidal to pyriform, (155–)160–180(–185) × (115–)125–140(–145) µm (*n*=10, *s*=1); ostiolar region typically concolour with the rest of perithecia, ostiolar opening barely visible at 10×. Involucrellum present, scabrid, (25–)30–40(–45) µm thick (*n*=8, *s*=1), extends over the entire perithecium. Excipulum from brownish to blackish, 5–10(–12) µm thick (*n*=5, *s*=1), composed of 3–4 layers aprox. of flattened rectangular cells. Periphyses present, septate, simple, 10–13(–14) × 1–1.5. Paraphyses are absent. Hamathecial gelatine K-, I + red, K/I + blue. Asci bitunicate, exotunica disappears rapidly with maturity, clavate, 8-spored, located in the lower and lateral parts of the perithecial cavity, (22–)25–30(–33) × (9.5–)10–11 µm (*n*=8, *s*=1). They are located on the lower and lateral part of the ascoma. *Ascospores* simple, hyaline, from ellipsoidal to ovoid, (7–)8–9(–9.5) × (4.5–)4–5(– 5.5) µm (*n*=43, *s*=1).

*Pycnidia* not present.

#### Verrucaria dispartita

Vain. Résultats du Voyages du S.Y. Belgica en 1897-1898-1899 - Lichens: 38 (1903); Fig. 9. sp. nov. — MycoBank MB#408990 **(sp. 3)**

**Figure 9.**
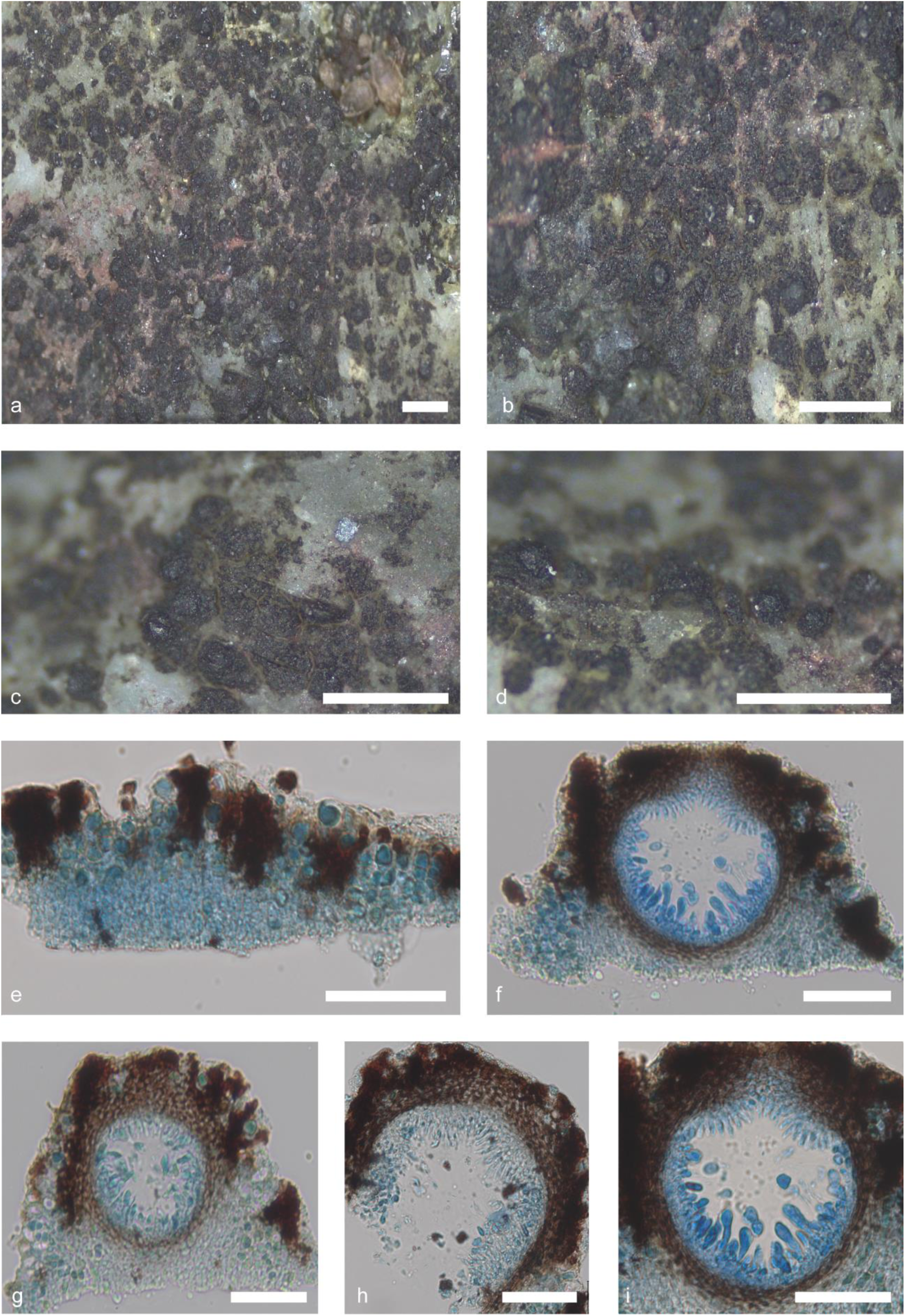
*Verrucaria dispartita* (*De Los Rios s. n.*). Thallus habitus and microscopic features. a-b. Habitus, c. Detail of the perithecia; d. Section of a perithecium, e. Thallus section showing the punctae; f. Thallus section with perithecia; g, h. Perithecia section detail; i. Asci, periphyses and excipulum. Scale bars: a, b, c, d: 1 mm, e, f, g, h, i: 50 µm. e-i: Lactophenol cotton blue. g-i: DIC.

*Typus.* Antartica, Gerlache Strait, Danco coast, Cape Anna, on fine grained non-calcareous rock, E. G. Racovitza 199.

*Thallus* epilithic, crustose, discontinuous, forming scattered small patches, up to 0.2 cm in diam, and 60–85 µm in height (*n*=6, *s*=2); from black to blackish brown with some greenish tones, shiny. Thallus surface rough, cracks forming polygonal areoles, 0.15 × 0.90 mm. Surface flat to slightly convex; punctae present, numerous, throughout thallus, occasionally forming ridges, giving it a rough appearance, 20–60 × 15–40 µm. Hypothallus not present. Thallus in section paraplectenchymatous. Mycobiont cells from irregularly globose to polygonal, 2–5 × 2–3 µm (*n*=12, *s*=2). A translucent refringent layer absent. Phaenocortex absent. Pigmented layer on top to the algal cells absent. Photobiont randomly arranged throughout the thallus, sometimes look like irregular columns, from irregularly ellipsoidal to globose, (8–)9–12(–14) × (3–)4–7(–11) µm (*n*=14, *s*=2).

*Perithecia* sessile, shiny black colour, oval in shape with flattened tip. In section, from globose-ellipsoidal to pyriform, (112–)120–160(–170) × (105–)110–140(–150) µm (*n*=11, *s*=2); ostiolar region typically concolour with the rest of perithecia, ostiolar opening barely visible at 10×. Involucrellum present, scabrid, (25–)30–45(–50) µm thick (*n*=11, *s*=2), extends over the entire perithecium. Excipulum from brownish to blackish, 10–15(–17) µm thick (*n*=6, *s*=2), composed of 4–5 layers aprox. of flattened rectangular cells. Periphyses present, septate, simple, 10–12(– 15) × 1–1.5. Paraphyses are absent. Hamathecial gelatine K-, I + red, K/I + blue. Asci bitunicate, exotunica disappears rapidly with maturity, clavate, 8-spored, located in the lower and lateral parts of the perithecial cavity, (25–)30–45(– 50) × (10–)12–14 µm (*n*=10, *s*=2). They are located on the lower and lateral part of the ascoma. *Ascospores* simple, hyaline, from ellipsoidal to ovoid, (7–)8–9(–10) × (4.5–)5(–6) µm (*n*=33, *s*=2).

*Pycnidia* not present.

##### Additional specimens examined

Antartica, Livingston Island, Sally Rocks, 62°42’07’S 60°25’44’W, on rock. 0-5 m alt. 25 January 2014. *A. de los Ríos s. n.* Antartica, Livingston Island, Punta Barnard, 62°45’27’S 60°20’20’W, on rock. 0-5 m alt. 20 August 2024. *A. de los Ríos s. n*.

#### Verrucaria orangei

Fernández-Costas & Pérez-Ort.; Fig. 10. sp. nov. — (**sp. 6**)

**Figure 10.**
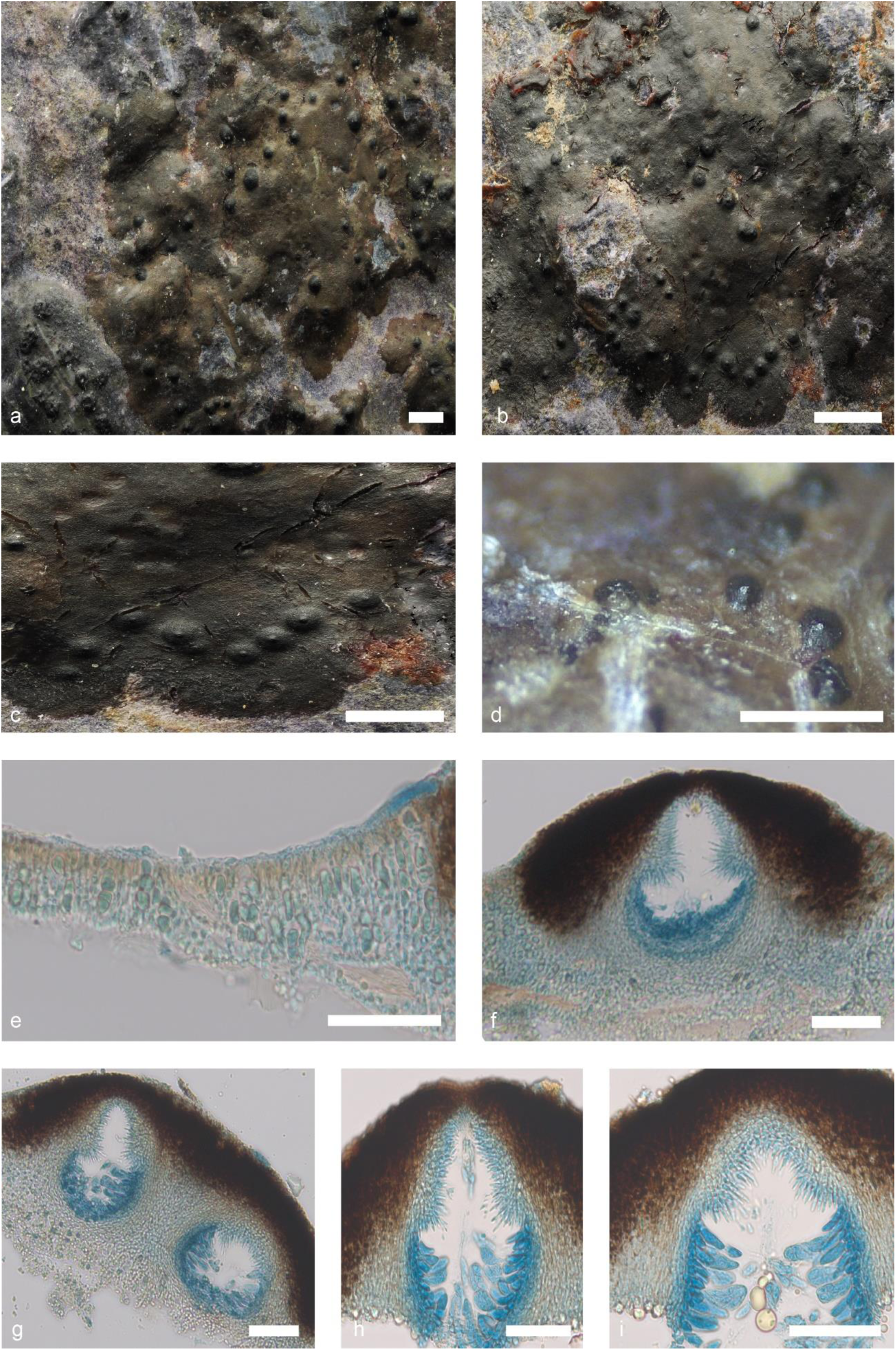
*Verrucaria orangei* (*Orange* 22632). Thallus habitus and microscopic features. a-b. Habitus, c. Detail of the perithecia; d. Section of a perithecium, e. Thallus section showing the punctae; f. Thallus section with perithecia; g, h. Perithecia section detail; i. Asci, periphyses and excipulum. Scale bars: a, b, c, d: 1 mm, e, f, g, h, i: 50 µm. e-i: Lactophenol cotton blue. g-i: DIC.

*Etymology. Verrucaria orangei* is named after the late Dr. Alan Orange, who undertook extensive research on the marine Verrucariaceae and Verrucariacae over the course of several decades. He made significant contributions to the field and was responsible for collecting the type material of this taxon.

*Typus.* Falkland Islands, East Falkland, Stanley Harbour, north side, 51°68’39’S 57°88’82’W, on shells of living

*Mytilus edulis*. 0 m alt. 1 February 2015. *A. Orange* 22632 (Welsh National Herbarium NMW).

*Thallus* epilithic, crustose, continuous, up to 2 cm in diam, thin, 80–90(–100) µm in height (*n*=5, *s*=2); from light ochraceous brown to blackish grey-brown, shiny. Thallus surface smooth; some fissures along the thallus. Hypothallus not present, sometimes darker line. Thallus in section paraplectenchymatous. Mycobiont cells irregularly spherical, 3–5 × 2–4 µm (*n*=14, *s*=2). A translucent refringent layer absent. Phaenocortex absent. Brown pigment between Phaenocortex and photobiont layer present, K-. Photobiont randomly arranged throughout the thallus, although photobiont columns can be occasionally observed; from irregularly ellipsoidal to globose, (7–)8–11(–13) × 4–9(–10) µm (*n*=13, *s*=2).

*Perithecia* semi-immersed up to 1/3–1/4, shiny blackish in colour, upper part broadly conical in shape, somewhat flattened at the apex. In section, from globose ellipsoid to pyriform, (120–) 130–150(–155) × (95–)100–125(–130) µm (*n*=6, *s*=2); ostiolar region often papillose, typically concolour with the rest of perithecia, ostiolar opening visible at 10×. Involucrellum present, (35–)40–50(–55) µm thick, extending to the middle of the perithecium. Excipulum hyaline, 10–15 µm thick (*n*=4, *s*=2), composed of c. 6 layers of flattened rectangular cells. Periphyses present, septate, simple, 10–15 × 1 µm (*n*=4, *s*=2). Paraphyses absent. Hamathecial gelatine K-, I + red, K/I + blue. Asci bitunicate, exotunica quickly disappears as ascus matures, clavate, 8-spored, located in the lower and lateral parts of the perithecial cavity, (25–)30–35(40–) × (6–)7–10(–12) µm (*n*=4, *s*=2). *Ascospores* simple, hyaline, from ellipsoidal to laterally elongated, (8–)9–12 × 4(–5) µm (*n*=43, *s*=2), without halonate perispore.

#### *Pycnidia* not observed

Notes – Occasionally different parts of the thallus can be observed to take on a bluish or even purplish colour, but this is due to the pigment of the shells on which it develops.

##### Additional specimens examined

Falkland Islands, West Falkland, Port Howard, NE of settlement, 51°61’61’S 59°52’0’W, on shells of living *Mytilus edulis*. 0 m alt. 25 January 2015. *A. Orange* 22449 (Welsh National Herbarium NMW).

#### Verrucaria delosriosii

Fernández-Costas & Pérez-Ort.; Fig. 11. sp. nov. — **(sp. 7)**

**Figure 11.**
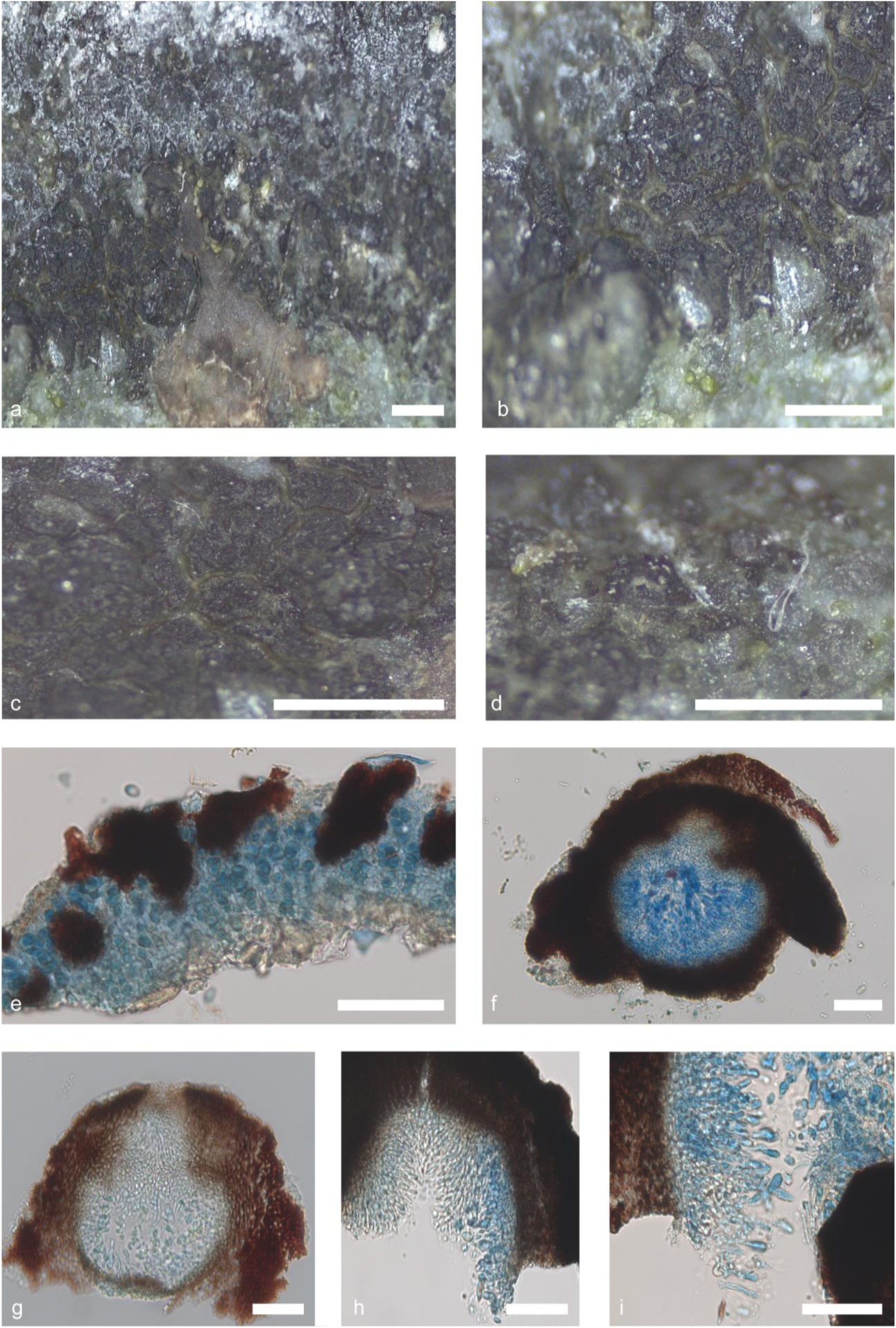
*Verrucaria delosriosi* (De Los Rios *s. n.*). Thallus habitus and microscopic features. a-b. Habitus, c. Detail of the perithecia; d. Section of a perithecium, e. Thallus section showing the punctae; f. Thallus section with perithecia; g. Perithecia section detail; h. Periphyses detail; i. Asci. Scale bars: a, b, c, d: 1 mm, e, f, g, h, i: 50 µm. e-i: Lactophenol cotton blue. g-i: DIC.

*Etymology. Verrucaria delosriosii* is named after Dr. Asunción de los Ríos, who has undertaken extensive collections of marine Verrucariaceae in Antarctica and the southern hemisphere during the last decades. She also collected the type material of this taxon.

*Typus.* Antartica, Livingston Island, Punta Polaca, 62°39’08’S 60°22’16’W, on rock. 0-5 m alt. 30 January 2014. *A. de los Ríos s. n* (MA-Lichen).

*Thallus* epilithic, crustose, areolate, up to 0.3 cm in diam, and 60–90 µm in height (*n*=5, *s*=1); from black to blackish brown, shiny. Thallus surface rough, cracks forming polygonal areoles, 0.2 × 0.8 mm. Surface flat to slightly convex, usually bordered by indistinct black margins, punctae present, numerous, throughout thallus, forming ridges, giving it a rough appearance, 25–55 × 10–40 µm. Hypothallus not present. Thallus in section paraplectenchymatous. Mycobiont cells from irregularly globose to polygonal, 3–4 × 2–3 µm (*n*=8, *s*=1). A translucent refringent layer absent. Phaenocortex absent. Brown pigment between Phaenocortex and photobiont layer present, K-. Photobiont randomly arranged throughout the thallus, from irregularly ellipsoidal to globose, (4–)5–8(–9) × (3–)4–6(–7) µm (*n*=10, *s*=1).

*Perithecia* sessile, shiny black colour, irregularly oval in shape. In section, from globose-ellipsoidal to pyriform, (230–)235–250(–255) × (170–)180–200(–205) µm (*n*=7, *s*=1); ostiolar region typically concolour with the rest of perithecia, ostiolar opening no visible at 10×. Involucrellum present, scabrid, (50–)60–70(–75) µm thick (*n*=6, *s*=1), extends over the entire perithecium. Excipulum from brownish to blackish, barely distinguishable from the excipulum, 10–15(–20) µm thick (*n*=6, *s*=1), composed of c. 5 layers aprox. of flattened rectangular cells. Periphyses present, septate, simple, (10–) 13–15 × 1–1.5. Paraphyses are absent. Hamathecial gelatine K-, I + red, K/I + blue. Asci bitunicate, exotunica disappears rapidly with maturity, clavate, 8-spored, located in the lower and lateral parts of the perithecial cavity, (35–)40–50(–55) × (10–)12–14 µm (*n*=8, *s*=1). They are located on the lower and lateral part of the ascoma. *Ascospores* simple, hyaline, from ellipsoidal to ovoid, (7–)8–10(–11) × (4–)5–6(–6,5) µm (*n*=41, *s*=1).

*Pycnidia* not present.

#### Verrucaria austroditmarsica

Fernández-Costas, Schiefelbein & Pérez-Ort.; Fig. 12. sp. nov. — **(sp. 8)**

**Figure 12.**
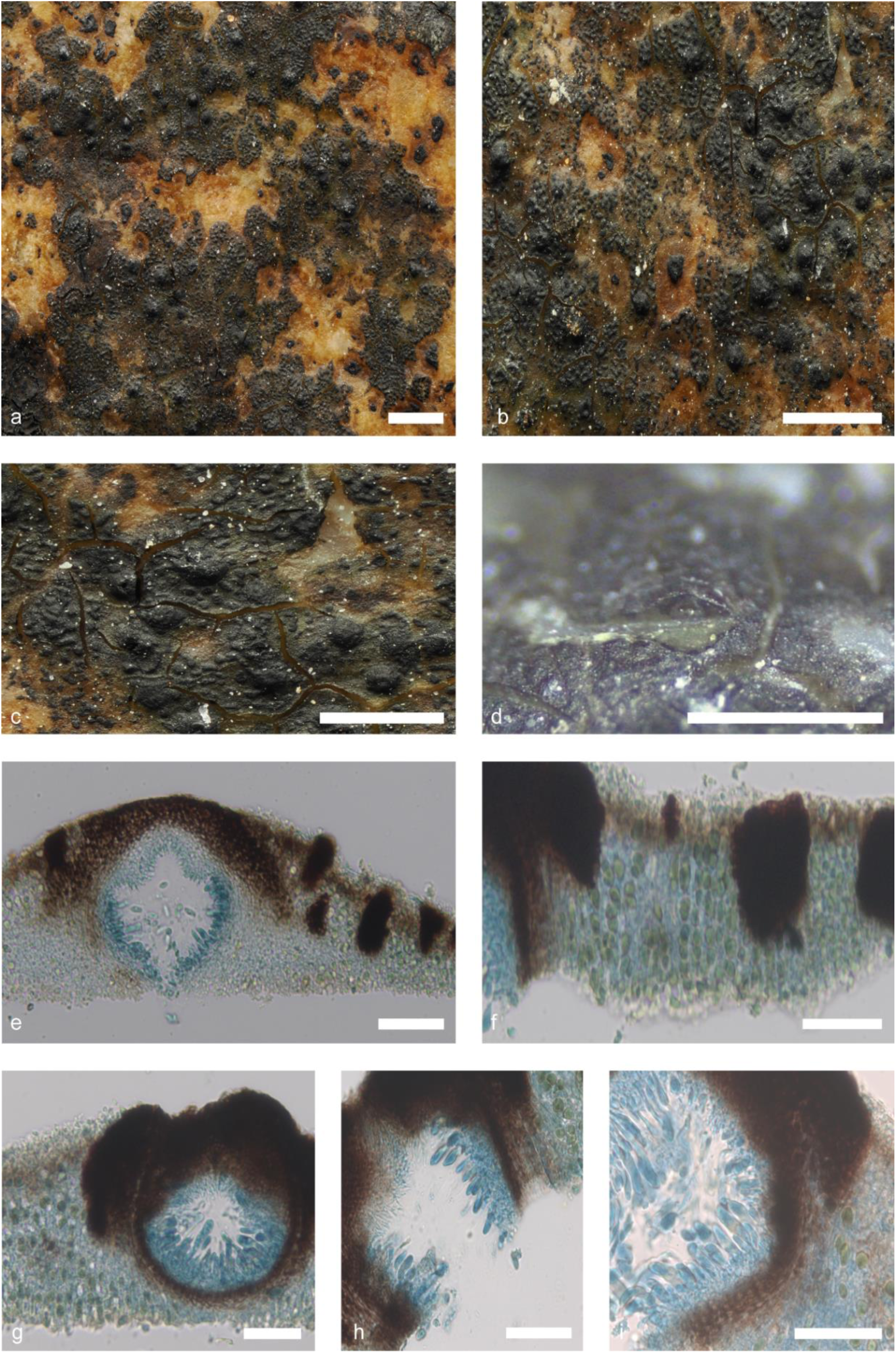
*Verrucaria austroditmarsica* (*Schiefelbein* 6628). Thallus habitus and microscopic features. a-b. Habitus, c. Detail of the perithecia; d. Section of a perithecium, e. Thallus section with perithecia; f. Thallus section showing the punctae; g-h. Perithecia section detail; i. Asci and excipulum. Scale bars: a, b, c, d: 1 mm, e, f, g, h, i: 50 µm. e-i: Lactophenol cotton blue. g-i: DIC.

*Etymology.* The specific epithet *austroditmarsica* refers to the marked similarity of the species to *V. ditmarsica*, and the fact that the newly discovered species is endemic to the Southern Hemisphere.

*Typus.* Chile, Magallanes and Chilean Antarctic Region, Province of Magallanes, San Gregorio, Strait of Magallanes, coast E of Puerto Sara, 70°06’39”W, 52°34’38”S, on sedimentary rock. 0 m alt. 20 January 2023. *U. Schiefelbein* 6628 (MA-Lichen).

*Thallus* epilithic, crustose, continuous, up to 3 cm in diam, thin, 75–110(–120) µm (*n*=9, *s*=4) in height; from black to greenish-black greenish-brown in colour, shiny. Thallus surface rough, with some cracks and fissures that form patches, large number of punctae scattered throughout the thallus, (30–)40–60(–70) × (15–)20–35(–40) µm (*n*=12, *s*=4), sometimes punctae can be connected to give the appearance of ridges. Hypothallus not present. Thallus in section paraplectenchymatous. Mycobiont cells slightly globose, 4–5 × 2–4 µm (*n*=24, *s*=4). A translucent refringent layer absent. Phaenocortex absent, sometimes a thin, discontinuous layer 5–10 µm thick is visible in some areas (*n*=5, *s*=4). Brown pigmented layer above the photobionts present, K-. Photobiont cells arranged in irregular vertical columns, sometimes totally unordered, from irregularly ellipsoidal to globose, (8–)9–13(–14) × (4–)5–9(–10) µm (*n*=17, *s*=4).

*Perithecia* semi-immersed up to 1/3–1/2, sometimes grouped in pairs, shiny or matt blackish in colour, from pyriform to oval, (100–)110–150(–155) × (85–)90–150(–160) µm (*n*=11, *s*=4); ostiolar region non-papillate, typically concolour, ostiolar opening visible at 10×. Involucrellum present, 25–35(–40) µm thick (*n*=10, *s*=4), extending to the middle of the perithecium. Excipulum brownish, 10–15 µm thick (*n*=8, *s*=4), composed of c. 6 layers of flattened rectangular cells. Periphyses present, septate, simple, 15–20 × 1–1.5 µm (*n*=9, *s*=4). Paraphyses absent. Hamathecial gelatine K-, I + red, K/I + blue. Asci bitunicate, exotunica quickly disappears as ascus matures, clavate, 8-spored, located in the lower and lateral parts of the perithecial cavity, (25–)30–40(–45) × 8–10(–12) µm (*n*=10, *s*=4). *Ascospores* simple, hyaline, from ellipsoidal to laterally elongated, (10–)12–17(–20) × 4–5 µm (*n*=70, *s*=4), without halonate perispore.

*Pycnidia* rare, immersed, irregularly distributed on the thallus, inconspicuous in dark thallus; from narrowly ellipsoid to globose, (50–)60–75(–82) × (37–)40–50(–55) µm (n=3, s=1), pycnidial wall hyaline. Conidiospores bacillar, hyaline, 3–4 × 1 µm (n=21, s=1).

Notes – Perithecia often go unnoticed because of the dark colours of the thallus, they hardly protrude from the thallus and even with a magnifying glass the ostiolar region is not visible. The ostiolar region is not visible under magnification, it is coloured with the involucre and the rest of the thallus, except in the lighter and brighter thalli where the region is slightly higher and more distinct from the rest of the involucrellum.

##### Additional specimens examined

Chile, Magallanes and Chilean Antarctic Region, Intertidal and *Nothofagus* Forest, 54°56’28”S, 67°15’3”W, on rock. 0-4 m alt. 22 January 2008. S. *Pérez-Ortega s. n.* (MA-Lich) – Chile, Magallanes and Chilean Antarctic Region, Province of Magallanes, San Gregorio, Strait of Magallanes, coast E of Puerto Sara, 70°06’39”W, 52°34’38”S, on sedimentary rock. 0 m alt. 20 January 2023. *U. Schiefelbein* 6628 – Falkland Islands, East Falkland, Stanley Harbour, 51°41’13’S 57°48’15’W, on roks. 0-5 m alt. 9 January 2011. *A. Orange* 19457 (Welsh National Herbarium NMW) – Falkland Islands, West Falkland, Roy Cove, 51°32’05’S 60°23’05’W, on roks. 15 m alt. 17 January 2011. *A. Orange* 19927 (Welsh National Herbarium NMW).

#### Verrucaria yagani

Fernández-Costas, Schiefelbein & Pérez-Ort.; Fig. 13. sp. nov. — **(sp. 10)**

**Figure 13.**
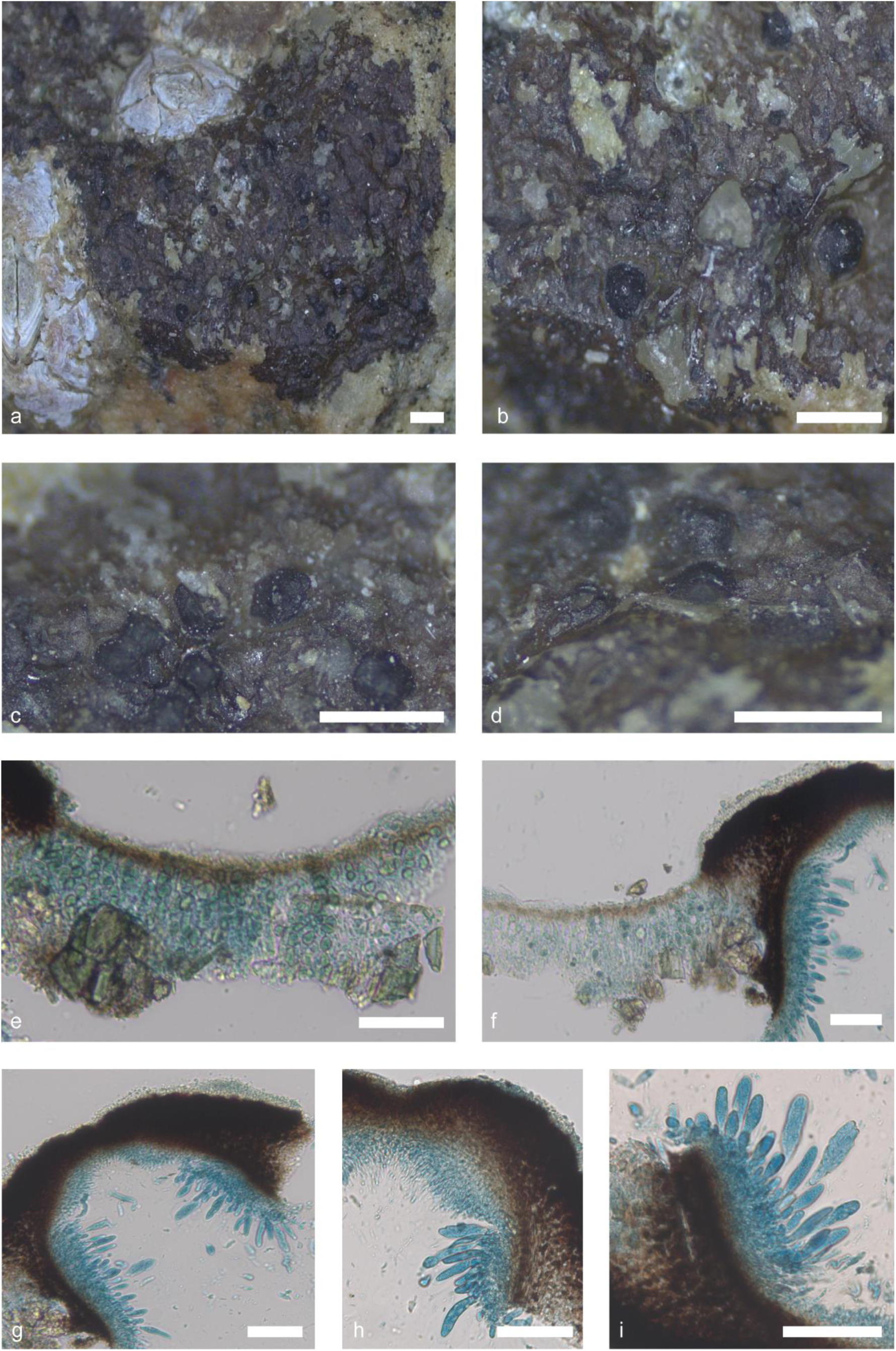
*Verrucaria yagani* (Pérez-Ortega *s. n.*). Thallus habitus and microscopic features. a-b. Habitus, c. Detail of the perithecia; d. Section of a perithecium; e. Thallus section showing the punctae; f. Thallus section with perithecia; g. Perithecia section detail; h. Asci, excipulum and periphyses; i. Asci and excipulum. Scale bars: a, b, c, d: 1 mm, e, f, g, h, i: 50 µm. e-i: Lactophenol cotton blue. g-i: DIC.

*Etymology.* The specific epithet *yagani* refers to the Yahgan, an indigenous people of the Southern Cone of South America. Their traditional territory encompasses the islands south of Isla Grande de Tierra del Fuego and extends to Cape Horn, marking them as the world’s southernmost indigenous population.

*Typus.* Chile, Magallanes and Chilean Antarctic Region, Navarino Island, Cove near Puerto Navarino in front of Hoste Island, 54°55’48”S, 68°20’45”W, on rock. 0-15 m alt. 27 January 2008*. S. Pérez-Ortega s. n.* (MA-Lich).

*Thallus* epilithic, crustose, continuous, up to 3 cm in diam, and 60–90(–110) µm in height (*n*=20, *s*=15); from olive to blackish brown, slightly greenish brown, shiny. Thallus surface smooth; some fissures along the thallus. Hypothallus not present, sometimes darker line. Thallus in section paraplectenchymatous. Mycobiont cells from irregularly globose to polygonal, 3–5 × 2–4 µm (*n*=36, *s*=15). A translucent refringent layer absent. Phaenocortex absent. A brown pigment layer, K-, is located above photobiont layer. Photobiont cells arranged in irregular vertical columns, sometimes appearing not to form true columns, from irregularly ellipsoidal to globose, (5–)8–12(–14) × (4–)5–7(–10) µm (*n*=39, *s*=15).

*Perithecia* semi-immersed to sessile, groups of up to three per areole, shiny black colour, from irregularly oval to pyriform in shape. In section, from globose-ellipsoidal to pyriform, (150–)180–200(–240) × (110–)150–200(–212) µm (*n*=24, *s*=15); ostiolar region typically concolour with the rest of perithecia, papillose, ostiolar opening barely visible at 10×. Involucrellum present, (25–)30–50(–65) µm thick (*n*=26, *s*=15), extending to the middle of the perithecium and may even surround it. Excipulum brownish, 10–15 µm thick (*n*=20, *s*=15), composed of 5 layers aprox. of flattened rectangular cells. Periphyses present, septate, simple, (10–)15–20 × 1–1.5. Paraphyses absent. Hamathecial gelatine K-, I + red, K/I + blue. Asci bitunicate, exotunica disappears rapidly with maturity, clavate, 8-spored, located in the lower and lateral parts of the perithecial cavity, (25–)40–50(–60) × (8–)10–15 µm (*n*=25, *s*=15). They are located on the lower and lateral part of the ascoma. *Ascospores* simple, hyaline, from ellipsoidal to laterally elongated, (10–)12–17(–19) × (3–)4–5(–6) µm (*n*=154, *s*=15).

*Pycnidia* common, immersed, irregularly distributed on the thallus, inconspicuous, from bacilliform to globose, (30–)45–70(–75) × (30–)35–45(–50) µm (*n*=13, *s*=7), pycnidial wall hyaline, Conidiospores bacillar, hyaline, 3–4 × 1–2 µm (*n*=60, *s*=7).

*Additional specimens examined*: Falkland Islands, East Falkland, Darwin Harbour, 51°48’68’S, 58°57’60’W, on rock. 0-5 m alt. 13 January 2011. *A. Orange* 19769 (Welsh National Herbarium NMW) – Chile, Aysén Region, Aysén province, Chonos Archipelago, coast NNW of the village Puerto Aguirre, 73°31’08”W, 45°09’26”S, on rocks. 0-5 m alt. 17 February 2016. *U. Schiefelbein* 4330 (Herbarium Musei Britannici) – Chile, Region Los Lagos, Province of Palena, Huinay, Comau Fjord, Intertidal rocks at Punta Llonco, 42°20’36”S 72°27’24”W, on rock. 0 m alt. 19 December 2014. *S. Pérez-Ortega & A. de los Ríos* 3300 – Chile, Region Los Lagos, Province of Palena, Huinay, Comau Fjord, Intertidal rocks at Punta Llonco, 42°20’36”S 72°27’24”W, on rock. 0 m alt. 19 December 2014. *S. Pérez-Ortega & A. de los Ríos* 3301 – Chile, Region Los Lagos, Province of Palena, Huinay, Comau Fjord, Intertidal rocks at Punta Llonco, 42°20’36”S 72°27’24”W, on rock. 0 m alt. 19 December 2014. *S. Pérez-Ortega & A. de los Ríos* 3314 – Chile, Region Los Lagos, Province of Palena, Huinay, Comau Fjord, Intertidal rocks at Punta Llonco, 42°20’36”S 72°27’24”W, on rock. 0 m alt. 19 December 2014. *S. Pérez-Ortega & A. de los Ríos* 3394 – Chile, Los Lagos Region, Llanquihue Province, Los Muermos, pacific coast at Estaquilla, 73°50’10”W, 41°23’32”S, on rocks. 0-5 m alt. 19 February 2016. *U. Schiefelbein* (Herbarium Musei Britannici) – Chile, Los Rios Region, Valdivia Province, Corral, pacific coast at Chaihuin, 73°35’13”W, 39°56’22”S, on rocks. 0-5 m alt. 21 February 2016. *U. Schiefelbein* 4356 (Herbarium Musei Britannici) – Chile, Magallanes and Chilean Antarctic Region, Province of Magallanes, San Gregorio, Strait ofMagellanes, coast E of Puerto Sara, 70°06’39”W, 52°34’38”S, on sedimentary rock. 0 m alt. 20 January 2023. *U. Schiefelbein* 6629 – Chile, Magallanes and Chilean Antarctic Region, Province of Magallanes, San Gregorio, Strait ofMagellanes, coast E of Puerto Sara, 70°06’39”W, 52°34’38”S, on sedimentary rock. 0 m alt. 20 January 2023. *U. Schiefelbein* 6631 – Chile, Región de Magallanes y la Antártida Chilena, Holger Islet. Intertidal and Nothofagus forest., 54°56’28”S, 67°15’3”W, on rock. 0-5 m alt. 22 January 2008. *S. Pérez-Ortega s. n.* (MA-Lich) – Chile, Región de Magallanes y la Antártida Chilena, Native coigües forest in front of Snipe Island, 54°57’41”S, 67°11’36”W, on rock. 1-4 m alt. 23 January 2008. *S. Pérez-Ortega s. n.* (MA-Lich) – Chile, Magallanes and Chilean Antarctic Region, Isla Grande de Tierra de Fuego, Admiralty Sound, Filton Fjord, Area of the Thousand Waterfalls, 54°36’50”S, 70°28’45”W, on rock. 2 m alt. 12 December 2009. *S. Pérez-Ortega s. n.* (MA-Lich).

#### Verrucaria hydropunctarioides

Fernández-Costas & Pérez-Ort.; Fig. 14. sp. nov. — (**sp. 11**)

**Figure 14.**
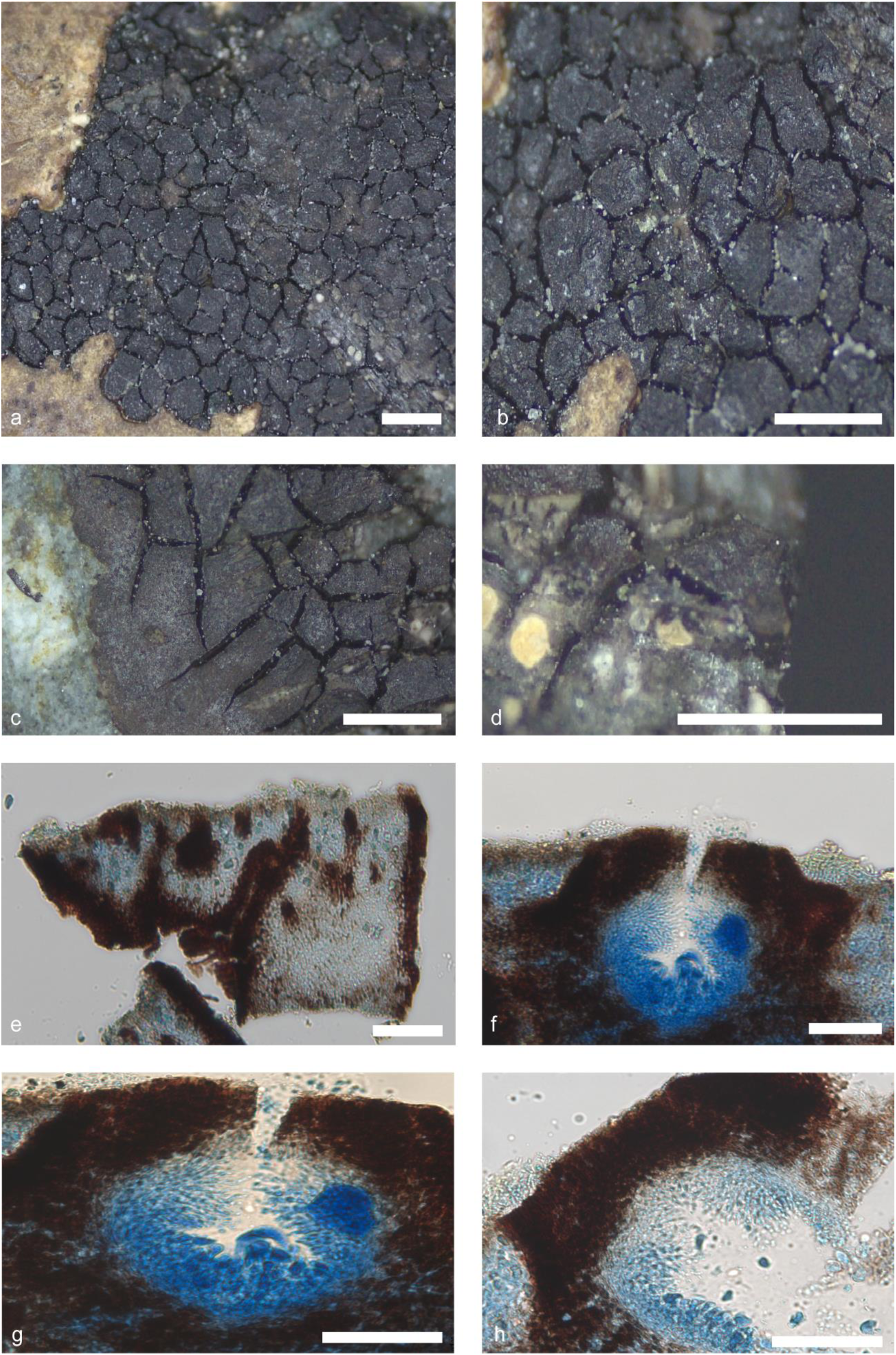
*Verrucaria hydropunctarioides* (Pérez-Ortega *s. n.*). Thallus habitus and microscopic features. a, b, c. Habitus; d. Section of a perithecium; e. Thallus section showing the punctae; f. Thallus section with immature perithecia; g-h. Immature perithecia section detail. Scale bars: a, b, c, d: 1 mm, e, f, g, h: 50 µm. e-h: Lactophenol cotton blue. g-h: DIC

*Etymology.* The specific epithet ‘*hydropunctarioides*’, refers to its similarity to species of the genus *Hydropunctaria*.

*Typus.* Chile, Magallanes and Chilean Antarctic Region, Tierra del Fuego, Basket Island, 54°44’25”S, 71°34’31”W, on rock. 0-5 m alt. 17 December 2009. *S. Pérez-Ortega s. n.* (MA-Lich).

*Thallus* epilithic, crustose, areolate, up to 1.5 cm in diam, and 160–190 µm in height (*n*=5, *s*=1); from black to blackish brown, shiny. Thallus surface rough, cracks form polygonal areolas, 0.2 × 0.8 mm. Areole surface flat to slightly convex, usually delimited by black margins, punctae present, scattered in thallus, giving it a rough appearance, 15–100 × 10–30 µm. Hypothallus not present. Thallus in section paraplectenchymatous. Mycobiont cells from irregularly globose to polygonal, 3–5 × 2–3 µm (*n*=12, *s*=1). A translucent refringent layer absent. Phaenocortex absent. A blackish brown pigment layer, K-, is located above photobiont layer. Photobiont randomly arranged throughout the thallus, from irregularly ellipsoidal to globose, (4–)5–6(–7) × (3–)4–5(–6) µm (*n*=12, *s*=1).

*Perithecia* immersed, immature, not visible on the thallus.

*Pycnidia* not observed.

#### Verrucaria labyrinthica

Fernández-Costas & Pérez-Ort.; Fig. 15. sp. nov. — **(sp. 12)**

**Figure 15.**
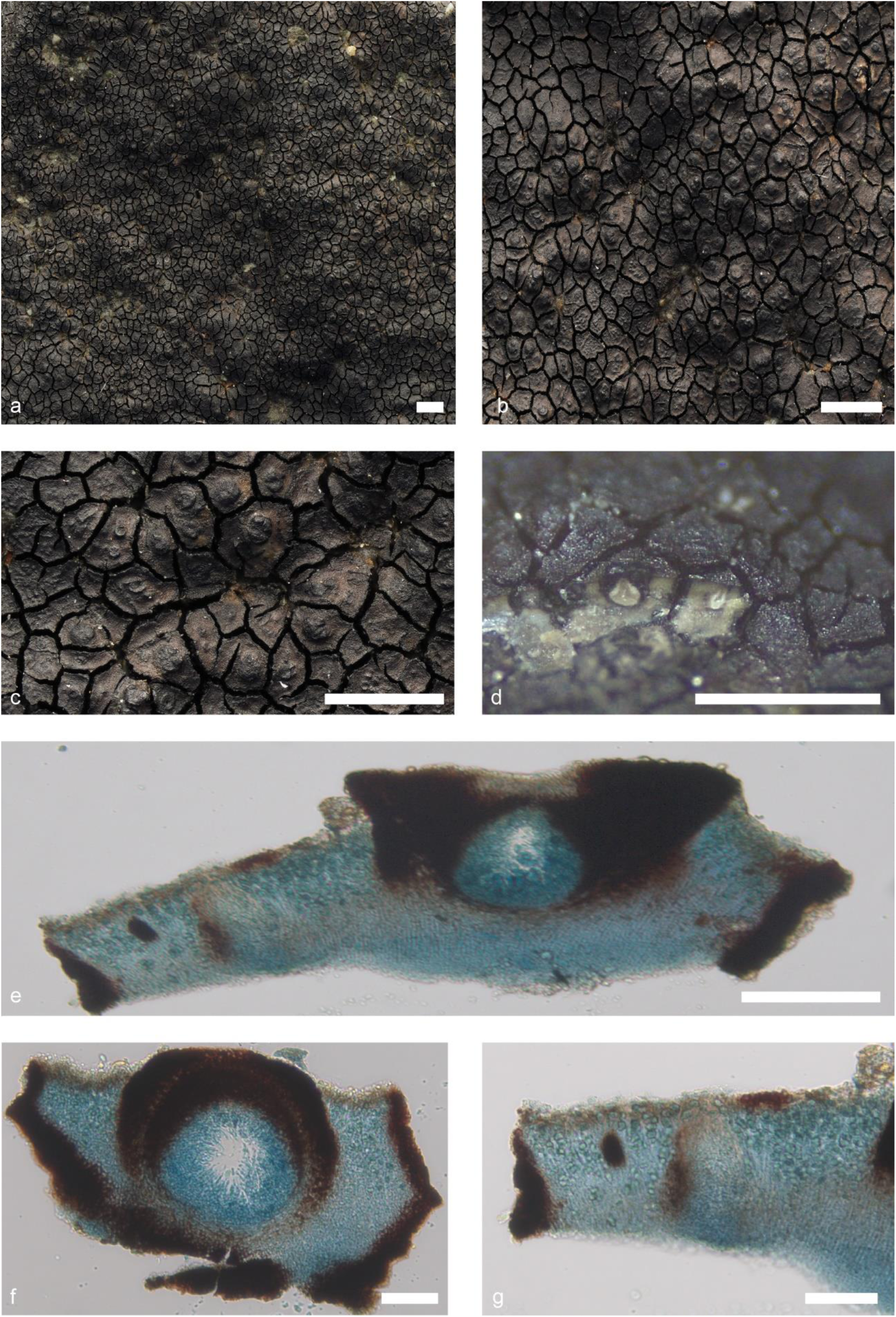
*Verrucaria labyrinthica* (Pérez-Ortega *s. n.*). Thallus habitus and microscopic features. a-b. Thallus habitus, c. Detail of perithecia in the areoles; d. Section of the thallus showing perithecia, e. Thallus section with perithecia with a visible brown layer; f. Perithecia section detail; g. Thallus section showing punctae. Scale bars: a, b: 2 mm, c, d: 1 mm, e, f, g: 50 µm. e-g: Lactophenol cotton blue. g-i: DIC.

*Etymology.* The specific epithet *labyrinthica*,’ derived from the Latin root *labyrinthus*, highlights the resemblance of the species thallus to a labyrinth.

*Typus.* Chile, Magallanes and Chilean Antarctic Region, Navarino Island, Cove near Puerto Navarino in front of Hoste Island, 54°55’48”S, 68°20’45”W, on rock. 0-15 m alt. 27 January 2008. S. *Pérez-Ortega s.n*. (MA-Lich).

*Thallus* epilithic, crustose, areolate, up to 6 cm in diam. thick, 150–200 µm (*n*=12, *s*=5) in height; from dark brown to olive-brown colour, shiny. Thallus surface rough, cracks form polygonal areolas, 0.15 × 1.10 mm. Areole surface flat to slightly convex, usually delimited by black margins, punctae present, scattered in thallus, giving it a rough appearance, 40–110 × 20–50 µm. Hypothallus not present. Thallus in section paraplectenchymatous. Mycobiont cells irregularly globose, 3–6 × 2–5 µm (*n*=24, *s*=5). A translucent refringent layer with cells with lipid content, basal layer is present, KI-. Phaenocortex absent. Brown pigmented layer above the photobiont layer present, K-. Photobiont cells arranged in irregular vertical columns, sometimes more irregularly distributed, from irregularly ellipsoidal to globose, (8–)9–12(–13) × 4–10(–12) µm (*n*=16, *s*=5).

*Perithecia* immersed to semi-immersed up to 1/4, shiny black colour, from irregularly oval to hemispherical in shape. In section, from globose-ellipsoidal to pyriform, (140–)150–200(–205) × (120–)125–180(–185) µm (*n*=12, *s*=5); ostiolar region non-papillate, typically concolour with the rest of perithecia, ostiolar opening barely visible at 10×. Involucrellum present, (50–)60–80(–85) µm thick (*n*=10, *s*=5), usually extending to the middle of the perithecium, and more rarely surrounding it. Excipulum slightly brown, 10–15 µm thick (*n*=11, *s*=5), composed of c. 4 layers of flattened rectangular cells. Periphyses present, septate, simple, 17–22(–25) × 1–1.5. Paraphyses absent. Hamathecial gelatine K-, I + red, K/I + blue. Asci bitunicate, exotunica quickly disappears as ascus matures, clavate, 8-spored, located in the lower and lateral parts of the perithecial cavity, (30–)35–50(–55) × (9–)10–15 µm (*n*=13, *s*=5). *Ascospores* simple, hyaline, ellipsoidal, (10–)11–14(–15) × 5–6 µm (*n*=40 *s*=5).

*Pycnidia* common, immersed, distributed on the thallus without any discernible pattern, with clusters of up to three, inconspicuous; from narrowly ellipsoid to globose, unilocular although more mature ones may become multilocular, (90–)100–150(–155) x (45–)50–90(–95) µm (*n*=13, *s*=5), pycnidial wall hyaline. Conidiospores bacillar, hyaline, 3– 4 x 1–2 µm (*n*=41, *s*=4).

##### Additional specimens examined

Chile, Magallanes and Chilean Antarctic Region, Holger Islet. Intertidal and supralittoral zone. 54°56’28”S, 67°15’3”W, on rock. 0-5 m alt. 22 January 2008. *S. Pérez-Ortega s.n.* (MA-Lich) – Chile, Magallanes and Chilean Antarctic Region, Picton Island, Intertidal, 54°59’29”S, 67°3’11”W, on rock. 1-5 m alt. 23 January 2008. *S. Pérez-Ortega s.n.* (MA-Lich) – Chile, Magallanes and Chilean Antarctic Region, Tierra del Fuego, Basket Island, 54°44’25”S, 71°34’31”W, on rock. 0-5 m alt. 17 December 2009. *S. Pérez-Ortega s.n.* (MA-Lich) – Falkland Islands, Weddell Island, N side of bay N of settlement, 51°89’38’S 60°90’22’W, on rock. 0-5 m alt. 22 January 2015. *A. Orange* 22330 (Welsh National Herbarium NMW).

#### Verrucaria rimosoareolata

Fernández-Costas, Schiefelbein & Pérez-Ort.; Fig. 16. sp. nov. — **(sp. 14)**

**Figure 16.**
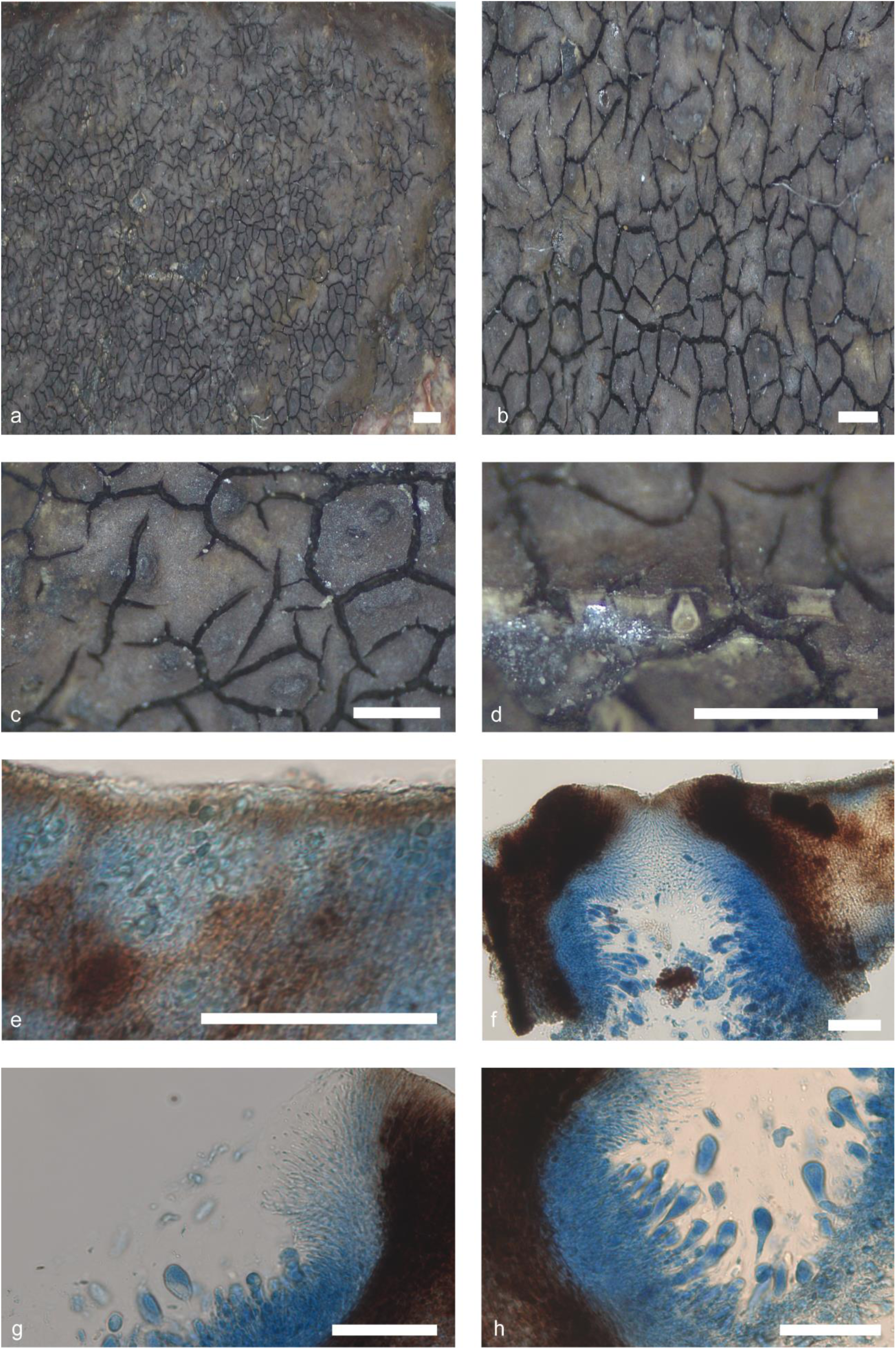
*Verrucaria rimososareolata* (Pérez-Ortega *s. n*.). Thallus habitus and microscopic features. a-b. Habitus, c. Detail of the perithecia; d. Section of a perithecium, e. Thallus section; f. Thallus section with perithecia; g. Periphyses; h. Asci. Scale bars: a, b, c, d: 1 mm, e, f, g, h: 50 µm. e-h: Lactophenol cotton blue. g-h: DIC.

*Etymology.* The specific epithet *rimosoareolata*, refers to the thallus of the species which structure vary from rimose to areolate.

*Typus.* Chile, Magallanes and Chilean Antarctic Region, Isla Grande de Tierra del Fuego, Seno del Almirantazgo, Gabriel’s Channel, 54°25’55”S, 70°7’48”W, on rock. 4 m alt. 12 December 2009. *S. Pérez-Ortega s. n.* (MA-Lich).

*Thallus* epilithic, crustose, continuous to areolate, up to 4 cm in diam, and 80–200 µm in height (*n*=5, *s*=2); from dark brown to dark yellowish brown, matt. Thallus surface smooth, cracks form polygonal areolas, 0.5 × 1.0 mm, punctae present, few, throughout thallus, barely visible, 40–70 × 15–30 µm. Areole surface flat to slightly convex, usually delimited by black margins. Hypothallus not present. Thallus in section paraplectenchymatous. Mycobiont cells from irregularly globose to polygonal, 4–6 × 2–4 µm (*n*=13, *s*=2). A translucent refringent layer absent. Phaenocortex absent. A brown pigment layer, K-, is located above photobiont layer. Photobiont arranged in columns throughout the thallus, from irregularly ellipsoidal to globose, (6–)7–10(–11) × (3–)4–6(–7) µm (*n*=12, *s*=2).

*Perithecia* immersed to semi-immersed, up to 1/3, shiny black colour, irregular ovoid in shape. In section, from globose-ellipsoidal to pyriform, (165–)170–240(–250) × (140–)150–220(–225) µm (*n*=8, *s*=2); ostiolar region concolour with the rest of the perithecium, ostiolar opening barely visible at 10×. Involucrellum present, (40–)50– 60(–65) µm thick (*n*=9, *s*=2), extends from the middle or to the base of the perithecium mixing with the excipulum. Excipulum blackish brown, 15–20) µm thick (*n*=6, *s*=2), composed of 5 layers aprox. of flattened rectangular cells. Periphyses present, septate, simple, 20–25 × 1–1.5. Paraphyses are absent. Hamathecial gelatine K-, I + red, K/I + blue. Asci bitunicate, exotunica disappears rapidly with maturity, clavate, 8-spored, located in the lower and lateral parts of the perithecial cavity, (30–)35–45(–47) × (9–)10–12(–13) µm (*n*=6, *s*=2). They are located on the lower and lateral part of the ascoma. *Ascospores* simple, hyaline, ellipsoidal, (12–)13–16(–17) × (5,5–)6–7(–8) µm (*n*=71, *s*=2).

*Pycnidia* present, immersed, irregularly distributed on the thallus, brownish, inconspicuous, ostiolar region concolour with the rest of pycnidia; from globose to bacilliform, (60–)70–90(–95) × (30–)35–45(–50) µm (*n*=5, *s*=2), pycnidial wall hyaline. Conidiospores from bacillar, hyaline, 3–4 × 1–1.5 µm (*n*=54, *s*=2).

##### Additional specimens examined

Chile, Magallanes and Chilean Antarctic Region, Province of Magallanes, Punta Arenas, Protected National Asset Cabo Froward, N of Cabo San Isidrio, 70°58’27”W, 51°45’27”S, on sedimentary rock. 0 m alt. 17 January 2023. *U. Schiefelbein* 6566.

#### Verrucaria pseudomucosa

Fernández-Costas & Pérez-Ort.; Fig. 17. sp. nov. — (**sp. 15**)

**Figure 17.**
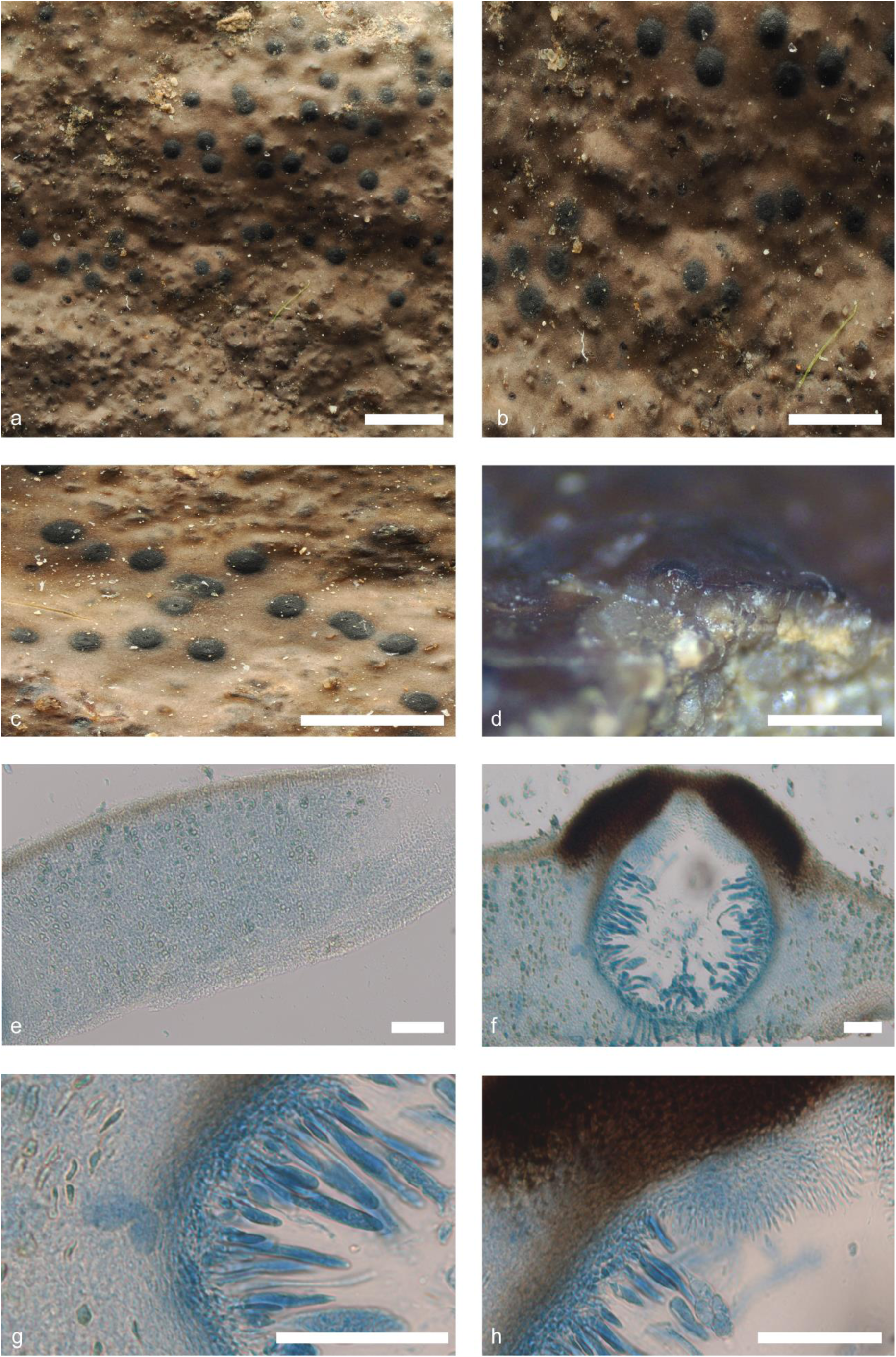
*Verrucaria pseudomucosa* (Pérez-Ortega *s. n.*). Thallus habitus and microscopic features. a-b. Habitus, c. Detail of the perithecia; d. Section of a perithecium, e. Thallus section; f. Thallus section with perithecia; g. Asci and excipulum; h. Periphyses. Scale bars: a, b, c, d: 1 mm, e, f, g, h: 50 µm. e-h: Lactophenol cotton blue. g-h: DIC.

*Etymology.* The specific epithet ‘*pseudomucosa*’, refers to its resemblance to the species *W. mucosa* from the northern hemisphere

*Typus.* Falkland Islands, East Falkland, Lafonia, North Arm, west side of inlet, 52°14’56’S 59°37’09’W, on rock. 0-5 m alt. 4 February 2015. *A. Orange* 22698 (Welsh National Herbarium NMW).

*Thallus* epilithic, crustose, continuous, up to 5-6 cm in diam, relatively thick, 175–240 µm (*n*=8, *s*=3) in height; from dark greenish brown to almost black to light greenish brown, matt, waxy appearance. Thallus surface smooth. Hypothallus not present, although sometimes a darker line is visible on lighter thalli. Punctae or ridges not present. Thallus in section paraplectenchymatous. Mycobiont cells irregularly globose-polygonal 4–7 × 3–5 µm (*n*=22, *s*=3). A translucent refringent layer with cells with lipid content, basal layer is present, KI-. Phaenocortex absent, sometimes a thin, discontinuous layer up to 5 µm thick is visible (*n*=5, *s*=3). Brown pigmented layer above photobionts present, K-. Photobiont cells arranged in vertical columns, from irregularly ellipsoidal to globose, (7–)8–12(–14) × 4–7(–8) µm (*n*=18, *s*=3).

*Perithecia* immersed to semi-immersed up to 1/3, up to 0.3 mm in diam, erumpent up to one third of the size of the ascoma, shiny blackish, appearance from globose to slightly conical, mountainous shape. In section, from globose to broadly ellipsoidal, (210–)250–300(–305) × (190–)200–260(–270) µm (*n*=10, *s*=3); ostiolar region non-papillate, typically concolour with the rest of perithecia, ostiolar opening visible at 10×. Involucrellum present, 30–50(–55) µm thick (*n*=7, *s*=3), extending up to the upper third of the perithecium. Excipulum hyaline, slightly brownish near the involucrellum, 12–15(–20) µm thick (*n*=5, *s*=3), composed of c. 6 layers of flattened rectangular cells. Periphyses present, septate, simple, 15–20 × 1 µm (*n*=3, *s*=3). Paraphyses absent. Hamathecial gelatine: K-, I+ red, K/I+ blue. Asci bitunicate, exotunica disappears rapidly with maturity, clavate, 8-spored, located in the lower and lateral parts of the perithecial cavity, (40–)50–80(–85) × (10–)12–15(–20) µm (*n*=13, *s*=3). *Ascospores* simple, hyaline, from ellipsoid to slightly oval, with two large lipid guttules giving the appearance of a septum, (11–)13–15 × (–5)6–7 µm (*n*=40, *s*=3), without halonate perispore.

*Pycnidia* common, immersed, distributed in clusters of brown spots, sometimes forming groups of up to 4 pycnidia, ostiolar region typically concolour with the rest of pycnidia, visible at 10×; unilocular, multilocular in when old; from narrowly ellipsoid to globose, (80–)90–120(–125) × (50–)60–90(–95) µm (*n*=3, *s*=3), pycnidial wall hyaline. Conidiospores bacillar straight to slightly curved, hyaline, 5–6 × 1 µm (*n*=38, s=3).

##### Additional specimens examined

Falkland Islands, West Falkland, Port Stephens, 52°14’25’S 60°85’42’W, on rock low of sea shore. 0-5 m alt. 29 January 2015. *A. Orange* 22582 (Welsh National Herbarium NMW) – Falkland Islands, West Falkland, Port Howard, NE of settlement, 51°61’61’S 59°51’66’W, on rock low of sea shore. 0-5 m alt. 25 January 2015. *A. Orange* 22445 (Welsh National Herbarium NMW)

#### Verrucaria chonorum

Fernández-Costas & Pérez-Ort.; Fig. 18. sp. nov. — **(sp. 16)**

**Figure 18.**
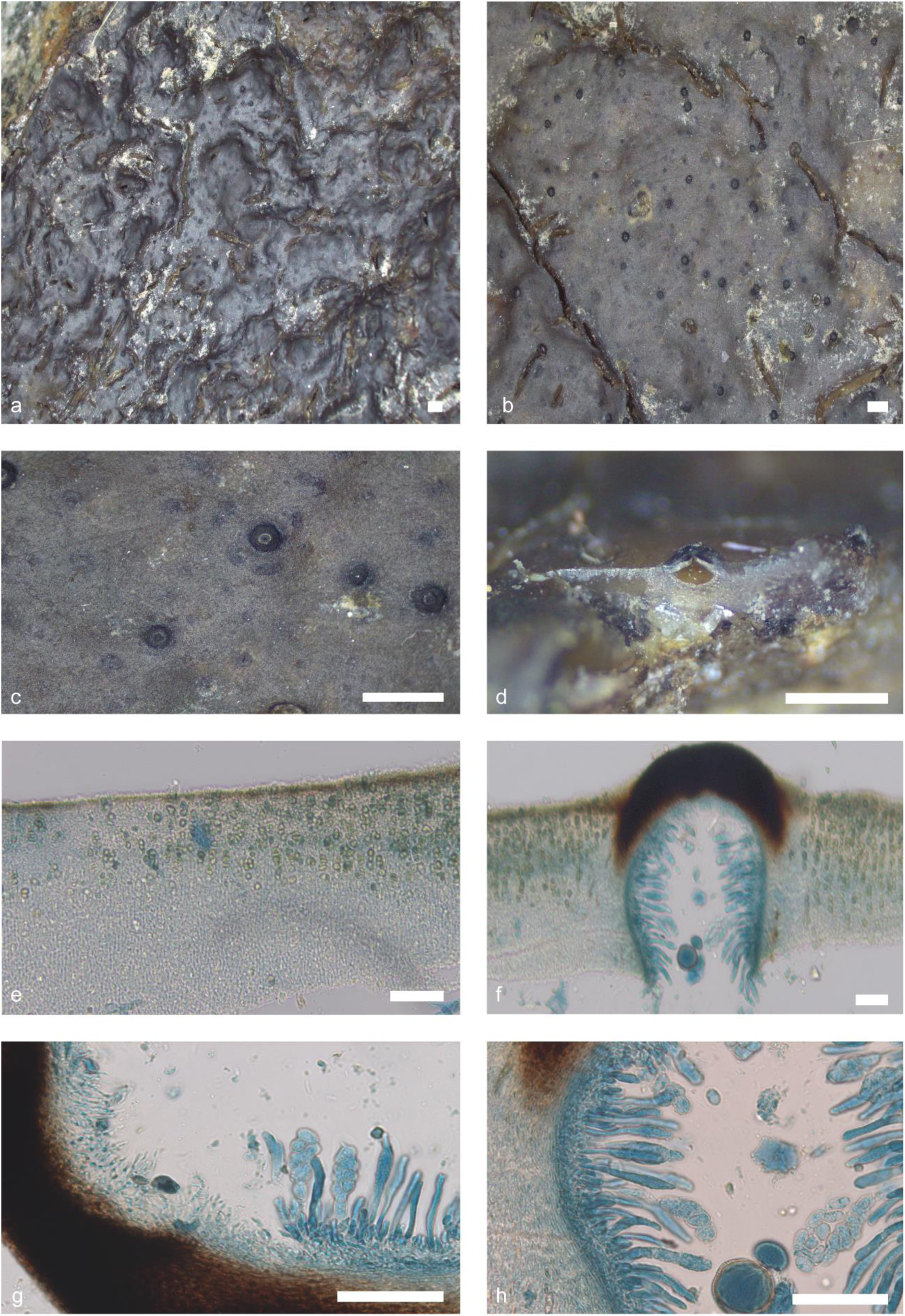
*Verrucaria chonorum* (S. Pérez-Ortega & A. de los Ríos 3305). Thallus habitus and microscopic features. a-b. Habitus, c. Detail of the perithecia; d. Section of a perithecium, e. Thallus section; f. Thallus section with perithecia; g. Asci and periphyses; h. Asci and excipulum. Scale bars: a, b, c, d: 1 mm, e, f, g, h: 50 µm. e-h: Lactophenol cotton blue. g-h: DIC.

*Etymology.* The specific epithet *chonorum* is a reference to the Chonos people, who were an extinct nomadic indigenous people or group of peoples of the archipelagos of Chiloé.

*Typus.* Chile, Region Los Lagos, Province of Palena, Huinay, Comau Fjord, Intertidal rocks below the biological station., 42°22’46’S, 72°24’54’W, on rock. 0 m alt. 19 December 2014. *S. Pérez-Ortega & A. de los Ríos* 3305.

*Thallus* epilithic, crustose, continuous, up to 5 cm in diam, relatively thick, 210–280 µm (*n*=8, *s*=4) in height; from dark ochraceous brown to very dark blackish brownish-brown, matt, waxy appearance. Thallus surface smooth, some herbarium specimens with cracks. Hypothallus present in some cases, black. Punctae or ridges not present. Thallus in section paraplectenchymatous. Mycobiont cells from irregularly polygonal to globose, (4–)5–6(–7) × 3–4 µm (*n*=28, *s*=4). A translucent refringent layer with cells with lipid content, basal layer is present, KI-. Phaenocortex absent, sometimes a thin, discontinuous layer 5–10 µm thick is visible (*n*=5, *s*=4). Brown pigmented layer above the photobionts present, K-. Photobiont cells arranged in vertical columns, from irregularly ellipsoidal to globose, (9–)10–14(–16) × 6–9(–10) µm (*n*=20, *s*=4).

*Perithecia* immersed to semi-immersed up to 1/4, shiny and blackish in colour. Globose in shape, (190–)200–300(– 305) × (240–)250–300(–310) µm (*n*=8, *s*=4); ostiolar region non-papillate, typically concolour with the rest of perithecia, ostiolar opening visible at 10×. Involucrellum present, 30–50(–60) µm thick (*n*=6, *s*=4), extending to the upper third of the perithecium. Excipulum hyaline, 14–17 µm thick (*n*=7, *s*=4) composed of c. 6 layers of flattened rectangular cells. Periphyses present, septate, simple, 15–20(–25) × 1-2 µm (*n*=4, *s*=4). Paraphyses absent. Hamathecial gelatine: K-, I+ red, K/I+ blue. Asci bitunicate, exotunica disappears rapidly with maturity, clavate, 8-spored, located in the lower and lateral parts of the perithecial cavity, (35–)40–55(–60) × 8–12(–17) µm (*n*=12, *s*=4). *Ascospores* simple, hyaline, from ellipsoidal to slightly oval, (9–)10–14(–15) × (5–)6–7(–8) µm (*n*=42, *s*=4), without halonate perispore.

*Pycnidia* common, immersed, randomly distributed on the thallus, inconspicuous, seen as lighter spots in the thallus, from narrowly ellipsoid to globose, unilocular, but multilocular when very mature (75–)90–175(–185) × (45–)50– 90(–95) µm (n=9, s=4), pycnidial wall hyaline. Conidiospores, bacillary, hyaline, and measure 4–5 × 1 µm (n=32, s=4).

##### Additional specimens examined

Chile, Region Los Lagos, Province of Palena, Huinay, Comau Fjord, Intertidal rocks below the biological station., 42°22’46’S 72°24’54’W, on rock. 0 m alt. 19 December 2014. *S. Pérez-Ortega & A. de los Ríos* 3408 – Chile, Region Los Lagos, Province of Palena, Huinay, Comau Fjord, Intertidal rocks at Punta Llonco, 42°20’36”S 72°27’24”W, on rock. 0 m alt. 19 December 2014. *S. Pérez-Ortega & A. de los Ríos* 3384 – Chile, Region Los Lagos, Province of Palena, Huinay, Comau Fjord, Intertidal rocks at Punta Llonco, 42°20’36”S 72°27’24”W, on rock. 0 m alt. 19 December 2014. *S. Pérez-Ortega & A. de los Ríos* 3368.

#### Verrucaria papillata

Fernández-Costas, Schiefelbein & Pérez-Ort.; Fig. 19. sp. nov. — **(sp. 17)**

**Figure 19.**
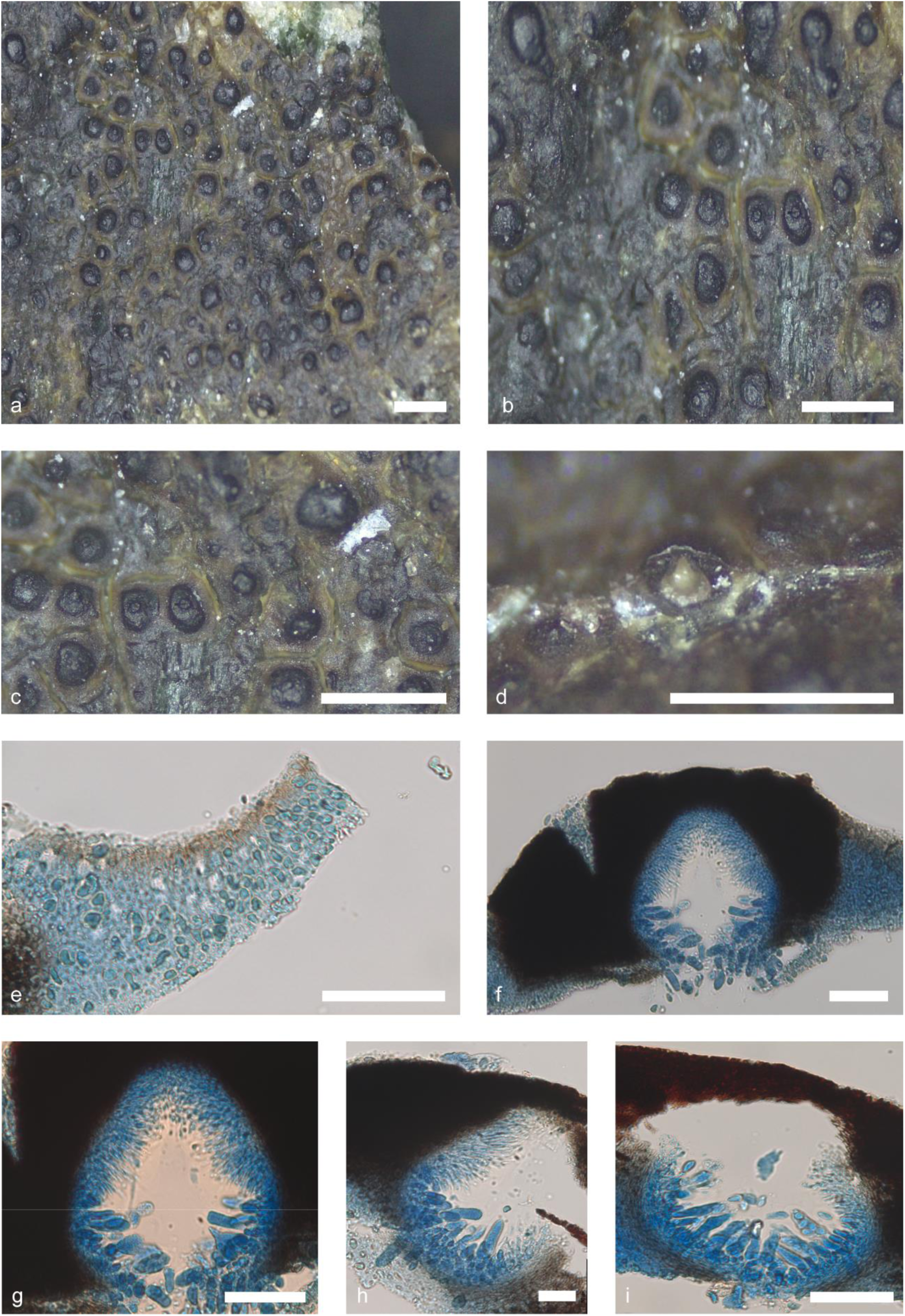
*Verrucaria papillata* (Pérez-Ortega *s. n.*). Thallus habitus and microscopic features. a-b. Habitus, c. Detail of the perithecia; d. Section of a perithecium; e. Thallus section; f. Thallus section with perithecia; g. Perithecia section detail; h. Asci and periphyses; i. Asci and excipulum. Scale bars: a, b, c, d: 1 mm, e, f, g, h, i: 50 µm. e-i: Lactophenol cotton blue. g-i: DIC.

*Etymology.* The specific epithet refers to the characteristic papillose ostiole of this species.

*Typus.* – Chile, Los Lagos Region, Quemchi, Chiloé Island, Pinquén beach. Intertidal zone, 42°8’35”S 73°28’15”W, on rock. 1-5 m alt. 2 December 2009. *S. Pérez-Ortega s. n.* (MA-Lich).

*Thallus* epilithic, crustose, continuous to areolate, up to 1.5 cm in diam, and 80–130 µm in height (*n*=10, *s*=3); from brownish brown to olive green brownish, shiny. Thallus surface smooth, some cracks around the perithecia form polygonal areolae, 0.45 × 0.85 mm. Surface flat to slightly convex. Hypothallus not present. Thallus in section paraplectenchymatous. Mycobiont cells from irregularly globose to polygonal, 4–5 × 2–4 µm (*n*=17, *s*=3). A translucent refringent layer absent. Phaenocortex absent, sometimes a thin, discontinuous layer 5 µm thick is visible in some areas (*n*=4, *s*=1). Pigmented layer above photobiont layer present, brownish, K-. Photobiont randomly arranged throughout the thallus, sometimes look like irregular columns, from irregularly ellipsoidal to globose, (6–)7–10(–11) × (4–)5–7(–8) µm (*n*=20, *s*=3).

*Perithecia* semi-immersed, up to 1/2, shiny black colour, round to slightly pyriform in shape with flattened tip. In section, from globose-ellipsoidal to pyriform, (150–)160–190(–200) × (100–)110–155(–160) µm (*n*=12, *s*=3); ostiolar region papillose, typically concolour with the rest of perithecia, ostiolar opening visible at 10×. Involucrellum present, (50–)55–80(–90) µm thick (*n*=14, *s*=3), extends over the entire perithecium and surround it. Excipulum from blackish, 10–15(–18) µm thick (*n*=9, *s*=3), composed of 5 layers aprox. of flattened rectangular cells. Periphyses present, septate, simple, 15–20 × 1–1.5. Paraphyses are absent. Hamathecial gelatine K-, I + red, K/I + blue. Asci bitunicate, exotunica disappears rapidly with maturity, clavate, 8-spored, located in the lower and lateral parts of the perithecial cavity, (30–)35–45(–50) × 10–12 µm (*n*=13, *s*=3). They are located on the lower and lateral part of the ascoma. *Ascospores* simple, hyaline, from ellipsoidal to ovoid, (7,5–) 8–10 (–11) × (4,5–)4–5(–6) µm (*n*=45, *s*=3).

*Pycnidia* present, immersed, irregularly distributed on the thallus, inconspicuous, ostiolar region concolour with the rest of pycnidia; from globose to bacilliform, (30–)35–40(–50) × (18–)20–30(–40) µm (*n*=3, *s*=1), pycnidial wall hyaline. Conidiospores not observed.

##### Additional specimens examined

Chile, Los Ríos Region, Valdivia Province, Corral, pacific coast at Chaihuin, 73°35’13”W, 39°56’22”S, on rocks. 0-5 m alt. 21 February 2016. *U. Schiefelbein* 4355 (Herbarium Misei Britannici) – Chile, Magallanes and Chilean Antarctic Region, Province of Magallanes, Punta Arenas, N of Puerto del Hambre, 70°58’06”W, 53°44’59”S, on sedimentary rock. 0-5 m alt. 7 January 2023. *U. Schiefelbein* 6451 – Falkland Islands, East Falkland, NW of Goose Green, New Haven, Shang Rookery Point, 50°43’56’S 59°12’59’W, on rock. 0-5 m alt. 13 January 2011. *A. Orange* 19769 (Welsh National Herbarium NMW).

#### Verrucaria lambi

Fernández-Costas, Schiefelbein & Pérez-Ort.; Fig. 20. sp. nov. — **(sp. 18)**

**Figure 20.**
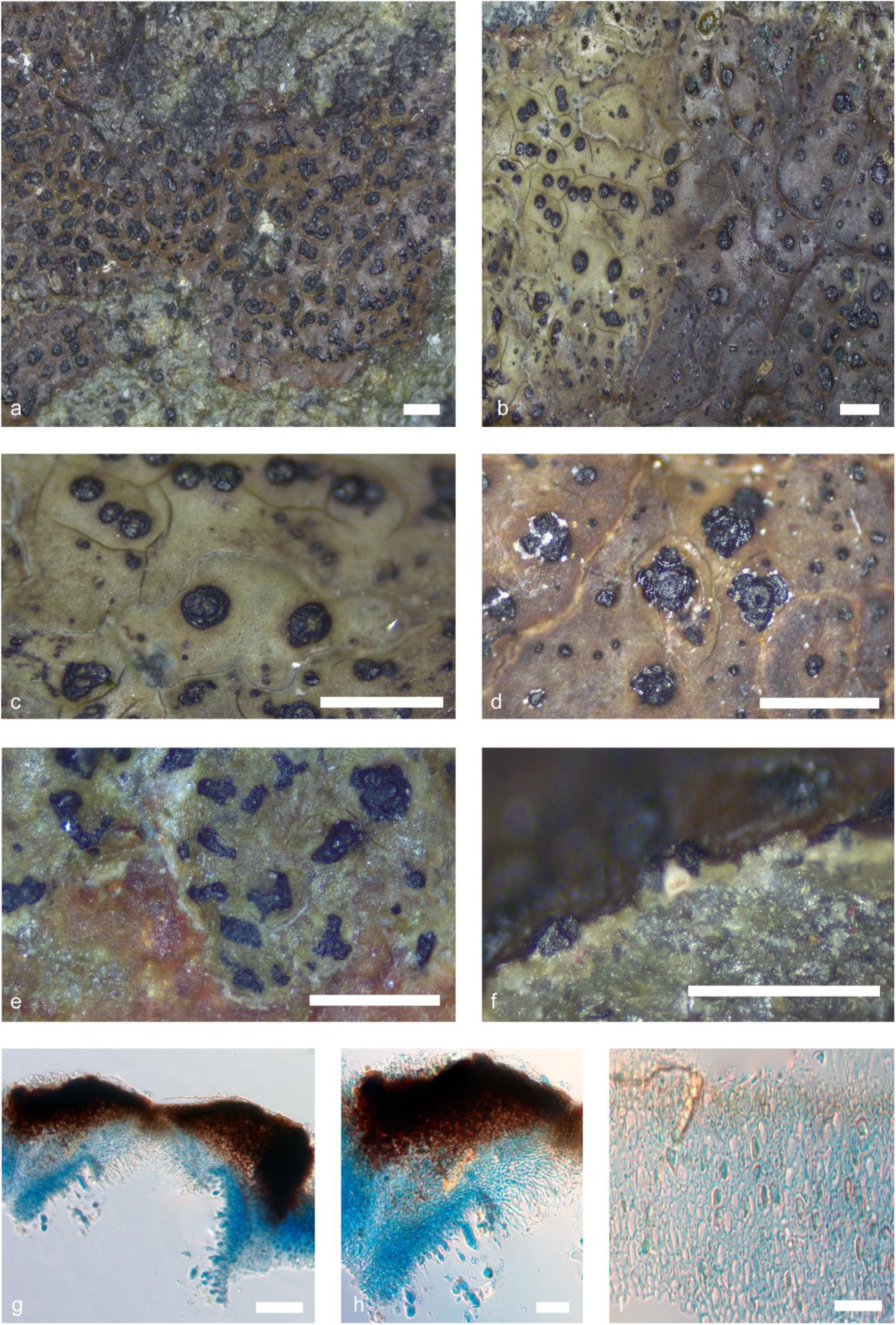
*Verrucaria lambi* (Pérez-Ortega *s. n.*). Thallus habitus and microscopic features. a-b. Habitus, c-d. Detail of the perithecia; e. Thallus ridges detail; f. Section of a perithecium; g. Perithecia section detail; h. Asci and periphyses; i. Thallus section detail. Scale bars: a, b, c, d, e, f: 1 mm, g, h, i: 50 µm. g-i: Lactophenol cotton blue. g-i: DIC.

*Etymology.* This species is named after Dr. Elke Mackenzie, who devoted part of her research to the study of marine Verrucariaceae in Antarctica and Southern South America. The epithet refers to the surname used in her lichenological publications.

*Typus.* Chile, Región de Magallanes y la Antártida Chilena, Tierra del Fuego, Beagle Channel, Chair Island, Darwin Bay, 54°53’59”S, 70°0’48”W, on rock. 0-5 m alt. 16 December 2009. *S. Pérez-Ortega s. n.* (MA-Lich).

*Thallus* epilithic, crustose, continuous to areolate, up to 3 cm in diam, thin, 60–200 µm (*n*=16, *s*=12) in height; from light olive green to brownish brown, pale lime green in shaded positions. Thallus surface smooth. In some specimens friable melanised structures may be observed, similar to punctae, but which may be slightly elongated but do not become true ridges. Hypothallus rarely present, thin whitish. Some specimens show small cracks, concolour with the thallus. Thallus in section paraplectenchymatous. Mycobiont cells from irregularly globose to slightly rectangular, up to 4–6 × 2–4 µm (*n*=43, *s*=12). A translucent refringent layer absent. Phaenocortex absent, occasionally a slight layer up to 5–10 µm thick (*n*=9, *s*=6). Brown pigment layer above photobiont layer present, K-. Photobiont cells randomly arranged throughout the thallus, irregularly ellipsoidal to globose, (5–)6–10(–12) × 4–7(–8) µm (*n*=31, *s*=12).

*Perithecia* semi-immersed up to 1/2 shiny and blackish in colour, rounded to slightly vertically ellipsoidal in shape, (90–)120–190(–195) × (85–)100–170(–180) µm (*n*=17, *s*=12); ostiolar region non-papillate, typically concolour, ostiolar opening visible at 10×. Involucrellum present, (50)65–90 µm thick (*n*=18, *s*=12), occasionally with irregular protuberances, extending to the middle of the perithecium. Excipulum hyaline, 10–15 µm thick (n=16, s=12), composed of c. 7 layers of flattened rectangular cells. Periphyses present, septate, simple, (14–)15–20 × 1(–1.5) µm (*n*=16, *s*=12). Paraphyses absent. Hamathecial gelatine: K-, I+ red, K/I+ blue. Asci bitunicate, exotunica disappears rapidly with maturity, clavate, 8-spored, occurring at the lower and lateral parts of the ascoma, (27–)30–40(–45) × (9–)11–14(–15) µm (*n*=10, *s*=8). *Ascospores* simple, hyaline, ellipsoidal to slightly oval, (7)8–10(11) × (4)5–6 µm (*n*=66, *s*=5), without halonate perispore.

*Pycnidia* common, randomly distributed on the thallus, unilocular, but multilocular when very mature; narrowly ellipsoid, (60–)70–100(–110) × (35–)40-55(–65) µm (*n*=9, *s*=6), pycnidial wall hyaline. Conidiospores bacillar, hyaline, 3–5 × 1(2) µm (*n*=22, *s*=6).

Notes – It should be noted that not all thallus have black melanized friable protuberances, as they are abundant in some thalli and scarcely present or absent in others. Colony of cyanobacteria can usually be found near the perithecia and in superficial hollows of the involucre. These colonies give a strong yellow reaction in the presence of KOH. Despite their external appearance and the visibility of the ostiole opening, most of the perithecia were still immature in many cases.

##### Additional specimens examined

Chile, Aysén Region, Aysén province, Chonos Archipelago, peninsula NE of the village Puerto Aguirre, 73°31’01’W, 45°09’31’S, on coastal beadrocks. 0-5 m alt. 17 February 2016. *U. Schiefelbein* 4334 (Herbarium Musei Britannici) – Chile, Los Ríos Region, Valdivia province, Corral, pacific coast at Chaihuin, 73°35’13’W, 39°56’22’S, on rocky coast. 0-5 m alt. 21 February 2016. *U. Schiefelbein* 4354 (Herbarium Musei Britannici) – Chile, Los Lagos Region, Quillaipe, Supralittoral rocks near Quillaipe, in the Reloncavi Inlet, 41°30’56’S, 72°46’47’W, on rock. 0-15 m alt. 29 March 2009. *S. Pérez-Ortega s. n.* (MA-Lich) – Chile, Magallanes and Chilean Antarctic Region, Navarino Island, Cove near Puerto Navarino in front of Hoste Island, 54°55’48”S, 68°20’45”W, on rock. 0-15 m alt. 27 January 2008. *S. Pérez-Ortega s. n.* (MA-Lich) – Chile, Los Lagos Region, Quemchi, Chiloé Island, Pinquén beach. Intertidal and supralittoral, 42°8’35”S 73°28’15”W, on rock. 1-5 m alt. 2 December 2009. S. Pérez-Ortega *s. n.* (MA-Lich) – Chile, Magallanes and Chilean Antarctic Region, Isla Grande de Tierra de Fuego, Admiralty Sound, Filton Fjord, Area of the Thousand Waterfalls, 54°36’50”S, 70°28’45”W, on rock. 2 m alt. 12 December 2009. *S. Pérez-Ortega s. n.* (MA-Lich) – Chile, Región de Magallanes y la Antártida Chilena, Tierra del Fuego, Beagle Channel, Chair Island, offshore islet, 54°54’02’S, 70°00’30’W, on rock. 0-5 m alt. 16 December 2009. *S. Pérez-Ortega* 429 – Chile, Región de Magallanes y la Antártida Chilena, Tierra del Fuego. Isla Basket. Intertidal, 54°42’13”S, 71°34’53”W, on rock. 0-1 m alt. 17 December 2009. *S. Pérez-Ortega s. n.* (MA-Lich) – Chile, Magallanes and Chilean Antarctic Region, Tierra del Fuego, Basket Island, 54°44’25”S, 71°34’31”W, on rock. 0-5 m alt. 17 December 2009. *S. Pérez-Ortega s. n.* (MA-Lich) – Falkland Islands, East Falkland, East of Stanley, Surf Bay, 51°41’53’S 57°46’01’W, on rock. 0-5 m alt. 10 January 2011. *A. Orange* 19509 (Welsh National Herbarium NMW) – Falkland Islands, West Falkland, Dunbar, 51°24’41’S 60°27’07’W, on rock. 0-5 m alt. 8 November 2015. *A. Orange* 23093 (Welsh National Herbarium NMW)

#### Verrucaria ceuthocarpoides

Fernández-Costas & Pérez-Ort.; Fig. 21. sp. nov. — **(sp. 19)**

**Figure 21.**
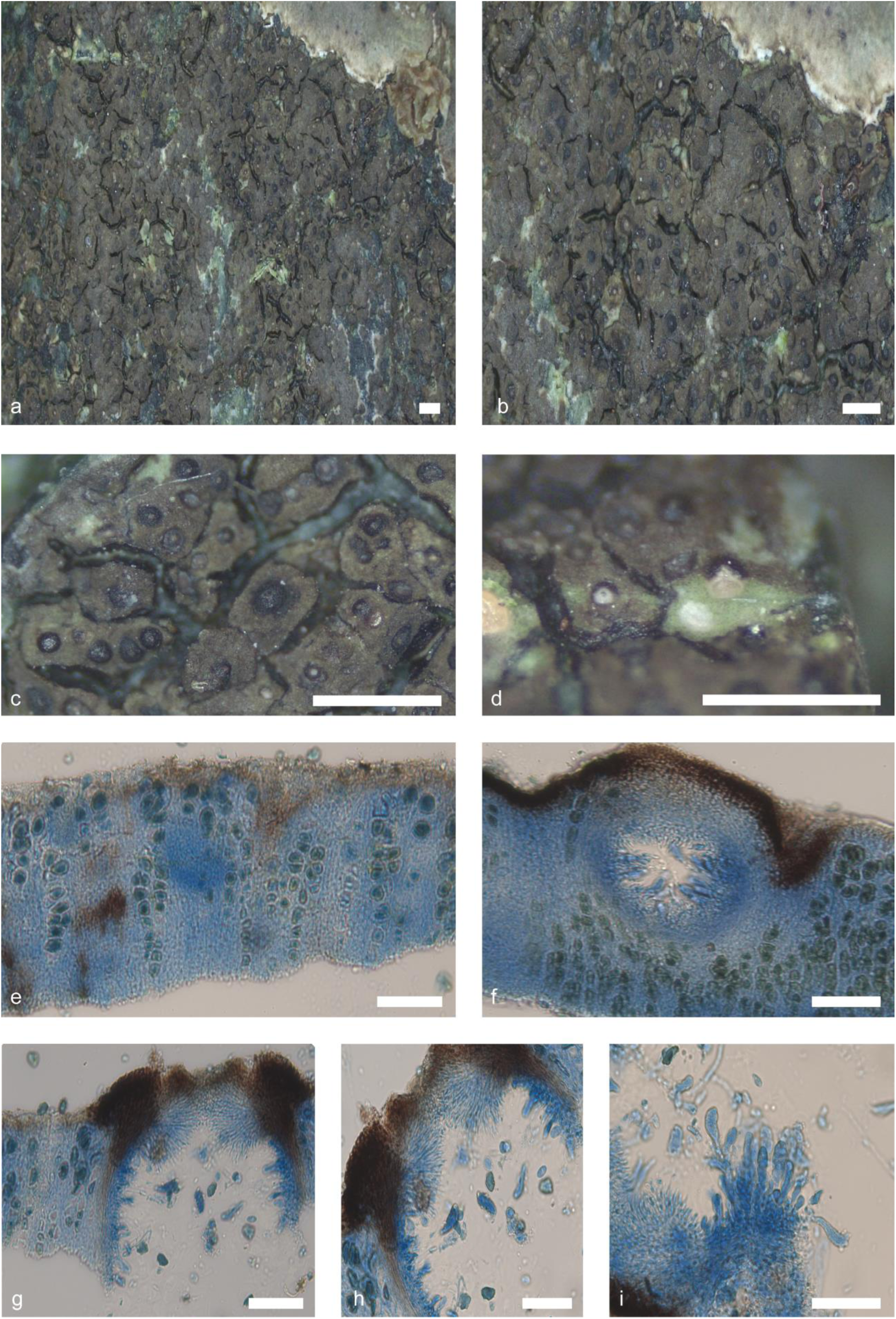
*Verrucaria ceuthocarpoides* (De Los Rios *s. n.*). Thallus habitus and microscopic features. a-b. Habitus, c. Detail of the perithecia; d. Section of a perithecium; e. Thallus section; f. Thallus section with perithecia; g. Perithecia section detail; h. Asci, periphyses and excipulum; i. Asci. Scale bars: a, b, c, d: 1 mm, e, f, g, h, i: 50 µm. e-i: Lactophenol cotton blue. g-i: DIC.

*Etymology.* The specific epithet ‘*ceuthocarpoides*’, refers to its similarity to *V. ceuthocarpa*, a species from the northern hemisphere.

*Typus.* Antartica, Livingston Island, St Kliment Ohridski Base, 62°38’30’S, 60°22’15’W, on rock. 0-5 m alt. 7 March 2024*. A. de los Ríos s. n* (MA-Lichen).

*Thallus* epilithic, crustose, continuous to areolate, up to 4.5 cm in diam, and 100–210 µm in height (*n*=20, *s*=6); from dark brown to blackish-brown to blackish-brown and may turn greenish in fresh specimens, shiny. Thallus surface smooth, cracks form polygonal areolas, 0.4 × 1.6 mm. Areole surface flat to slightly convex, usually delimited by black margins. Hypothallus sometimes visible as a continuous line at the thallus. Thallus in section paraplectenchymatous. Mycobiont cells from irregularly globose to polygonal, 4–6 × 2–4 µm (*n*=24, *s*=6). A translucent refringent layer with cells with lipid content, basal layer is present, KI-. Phaenocortex absent, sometimes a thin, discontinuous layer 10–15 µm thick is visible in some areas (*n*=7, *s*=5). A brown pigment layer, K-, is located above photobiont layer. Photobiont arranged in columns throughout the thallus, from irregularly ellipsoidal to globose, (8–)10–15(–16) × (6–)7–11(–12) µm (*n*=39, *s*=6).

*Perithecia* immersed to semi-immersed, up to 1/4, shiny black colour, round in shape with flattened tip. In section, from globose-ellipsoidal to pyriform, (112–)140–185(–195) × (110–)130–160(–165) µm (*n*=16, *s*=6); ostiolar region typically of a whiter shade than perithecium, ostiolar opening visible at 10×. Involucrellum present, (10–)15–25(– 27) µm thick (*n*=12, *s*=6), extends only in the upper part of the perithecium. Excipulum hyaline, 10–13(–15) µm thick (n=11, s=6), composed of 5–6 layers aprox. of flattened rectangular cells. Periphyses present, septate, simple, 10–15 × 1–1.5. Paraphyses are absent. Hamathecial gelatine K-, I + red, K/I + blue. Asci bitunicate, exotunica disappears rapidly with maturity, clavate, 8-spored, located in the lower and lateral parts of the perithecial cavity, (30–)35–50(–54) × (9–)10–13(–14) µm (*n*=16, *s*=6). They are located on the lower and lateral part of the ascoma. *Ascospores* simple, hyaline, ellipsoidal to ovoid, (8.5–)9–12(–12.5) × (5.5–)6–7(–8) µm (*n*=90, *s*=6).

*Pycnidia* present, immersed, irregularly distributed on the thallus, brownish, inconspicuous, ostiolar region concolour with the rest of pycnidia; from globose to bacilliform, (75–)80–100(–105) × (30–)35–50(–55) µm (*n*=12, *s*=5), pycnidial wall hyaline. Conidiospores from bacillar, hyaline, 3–4 × 1–1.5 µm (*n*=21, *s*=4).

##### Additional specimens examined

Antartica, Livingston Island, Caleta Española, 62°39’24’S, 60°21’57’W, on rock. 0-5 m alt. 21 January 2014. *A. de los Ríos s. n.* – Antartica, Livingston Island, Caleta Argentina, Pingüinera 62°39’57’S 60°23’49’W, on rock. 0-5 m alt. 11 August 2023. *A. de los Ríos s. n.* – Antartica, Livingston Island, Sally Rocks, 62°42’07’S 60°25’44’W, on rock. 0-5 m alt. 25 January 2014. *A. de los Ríos s. n.* – Antartica, Livingston Island, Punta Polaca, 62°39’08’S 60°22’16’W, on rock. 0-5 m alt. 30 January 2014. *A. de los Ríos s. n.* – Chile, Región de Magallanes y la Antártida Chilena, Tierra del Fuego. Isla Basket. Intertidal, 54°42’13”S, 71°34’53”W, on rock. 0-1 m alt. 17 December 2009. *S. Pérez-Ortega s. n.* (MA-Lich).

#### Verrucaria shukakensis

Fernández-Costas & Pérez-Ort.; Fig. 22. sp. nov. — **(sp. 20)**

**Figure 22.**
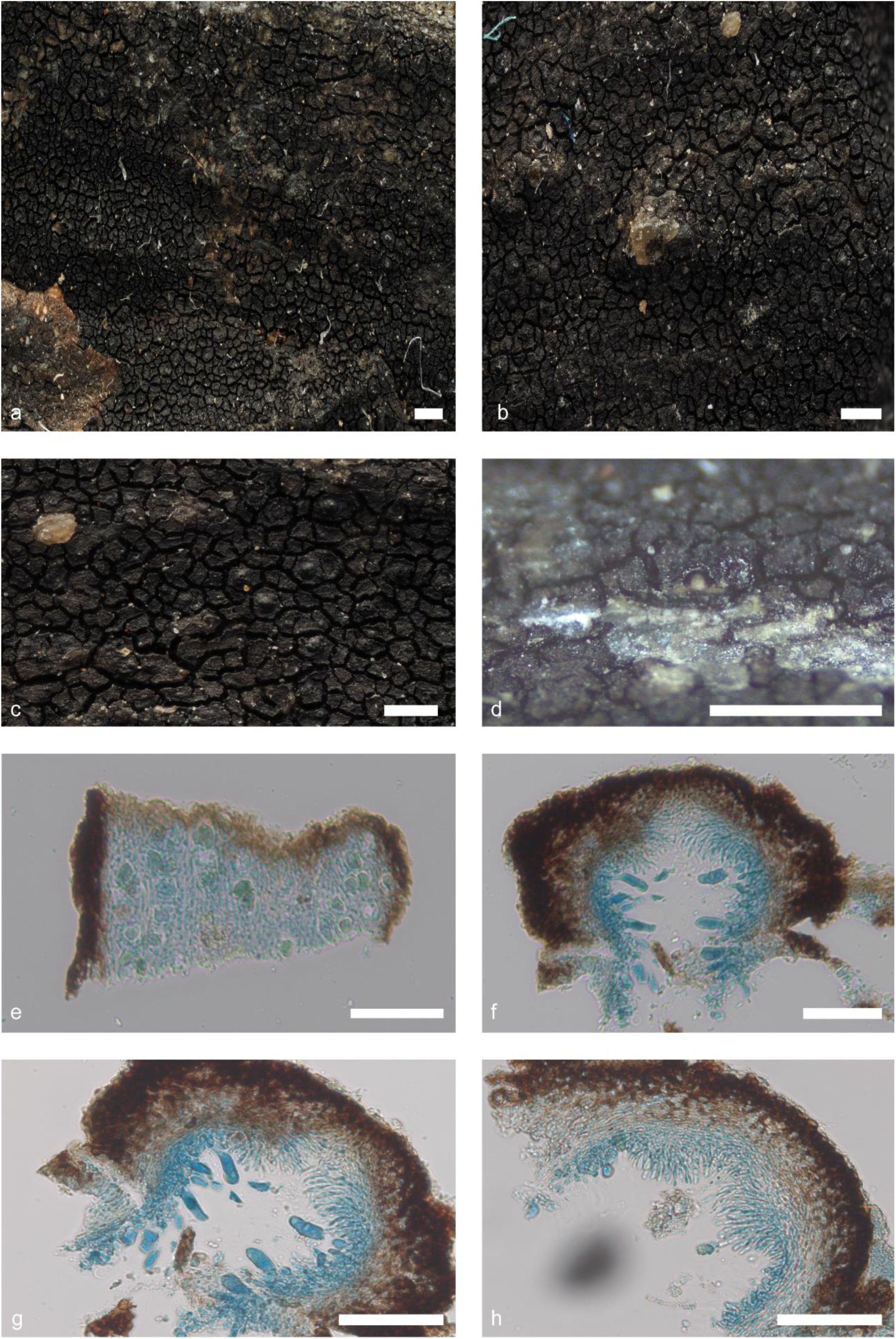
*Verrucaria shukakensis* (Pérez-Ortega *s. n.*). Thallus habitus and microscopic features. a-b. Habitus, c. Detail of the perithecia; d. Section of a perithecium; e. Thallus section; f. Perithecia section; g. Asci and periphyses; h. Periphyses and excipulum. Scale bars: a, b, c, d: 1 mm, e, f, g, h: 50 µm. e-h: Lactophenol cotton blue. g-h: DIC.

*Etymology.* The species epithet refers to the name given by the Yahgan people to the Picton Island, where the type specimen was collected

*Typus.* Chile, Magallanes and Chilean Antarctic Region, Picton Island, Intertidal, 54°59’29”S, 67°3’11”W, on rock. 1-5 m alt. 23 January 2008. *S. Pérez-Ortega s. n.* (MA-Lich)

*Thallus* epilithic, crustose, areolate, up to 3.5 cm in diam, thick, 70–150 µm (*n*=8, *s*=2) in height; from dark brown to blackish-brown, some thinner areas lighter in colour, shiny. Thallus surface smooth, areoles irregularly polygonal, 0.15 × 0.40 mm. Areole surface flat to slightly convex, usually delimited by black margins, when dehydrated, the edge usually rises slightly above the surface of the areole. Hypothallus not present. Thallus in section paraplectenchymatous. Mycobiont cells irregularly globose, 3–5 × 2–5 µm (*n*=11, *s*=2). A translucent refringent layer absent. Phaenocortex absent. Brown pigmented layer above photobiont layer present, K-. Photobiont cells arranged in irregular vertical columns, sometimes appearing not to form true columns, from irregularly ellipsoidal to globose, (8–)10–14(–15) × (7–)8–11 µm (*n*=12, *s*=2).

*Perithecia* immersed to semi-immersed up to 1/4, blackish in colour, shiny, from slightly oval to hemispherical in shape with an irregular surface. In section, from globose-ellipsoidal to pyriform, (100–)110–180(–185) × (90–)100– 160(–165) µm (*n*=10, *s*=2); ostiolar region non-papillate, typically concolour with the rest of the perithecia, ostiolar opening barely visible at 10×. Involucrellum present, (35–)20–40(–45) µm thick, extending to the middle of the perithecium. Excipulum hyaline, 10–15 µm thick (*n*=4, *s*=2), composed of c. 5 layers of flattened rectangular cells. Periphyses present, septate, simple, 15–17(–18) × 1 µm (*n*=4, *s*=2). Paraphyses absent. Hamathecial gelatine K-, I + red, K/I + blue. Asci bitunicate, exotunica quickly disappears as ascus matures, clavate, 8-spored, located in the lower and lateral parts of the perithecial cavity, (30–)35–45(–50) × (8–)10–15(–17) µm (*n*=4, *s*=2). *Ascospores* simple, hyaline, from ellipsoidal to ovoid, (7–)9–12 × 6–7 µm (*n*=57, *s*=2), without halonate perispore.

*Pycnidia* common, immersed, randomly distributed on the thallus, inconspicuous except in light zones, from narrowly ellipsoid to globose, unilocular, (30–)40–60(–65) × (25–)30–40(–43) µm (n=6, s=2), pycnidial wall hyaline. Conidiospores bacillar, hyaline, 3–4 × 1 µm (n=22, s=2).

##### Additional specimens examined

Chile, Magallanes and Chilean Antarctic Region, Navarino Island, cove near Puerto Navarino in front of Hoste Island, 54°55’48”S, 68°20’45”W, on rock. 0-15 m alt. 27 January 2008. *S. Pérez-Ortega s. n.* (MA-Lich).

#### Verrucaria austroamericana

Fernández-Costas, Schiefelbein & Pérez-Ort.; Fig. 23. sp. nov. — **(sp. 21)**

**Figure 23.**
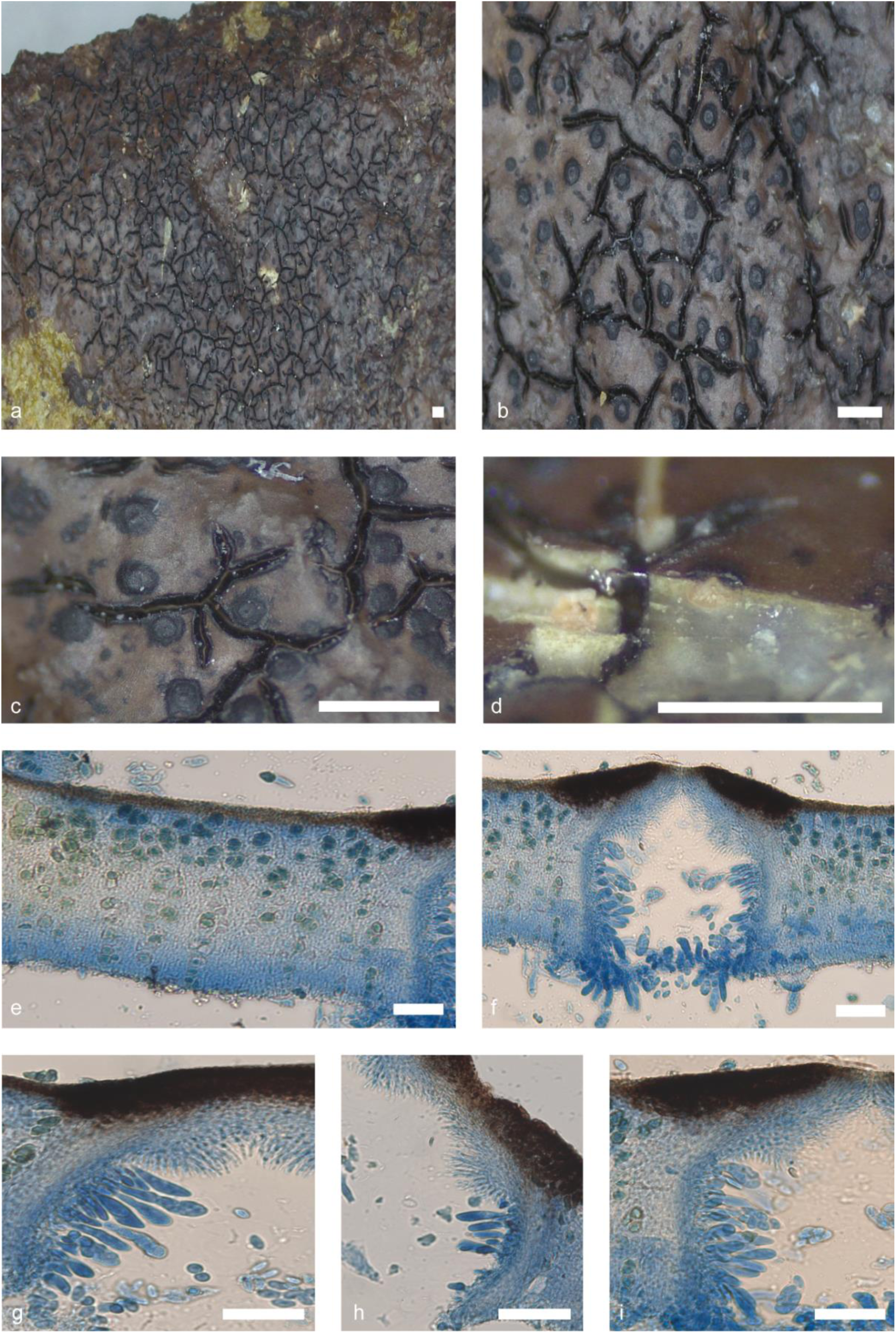
*Verrucaria austroditmarsica* (Orange 22634). Thallus habitus and microscopic features. a-b. Habitus, c. Detail of the perithecia; d. Section of a perithecium, e. Thallus section showing the punctae; f. Thallus section with perithecia; g, h, i. Asci, excipulum and periphyses. Scale bars: a, b, c, d: 1 mm, e, f, g, h, i: 50 µm. e-i: Lactophenol cotton blue. g-i: DIC.

*Etymology.* The epithet is a geographical term denoting the area in which the species has been colleted, specifically the southernmost region of the South American continent.

*Typus.* Chile, Magallanes and Chilean Antarctic Region, Isla Grande de Tierra del Fuego, Seno d’Agostini, Angelitos Cove, Peat bogs, 54°24’12”S, 70°26’24”W, on rock. 20 m alt. 13 December 2009. *S. Pérez-Ortega s. n.* (MA-Lich).

*Thallus* epilithic, crustose, continuous to areolate, up to 5.7 cm in diam, and 120–180 µm in height (*n*=20, *s*=6); from brownish brown to light ochraceous brown, shiny. Thallus surface smooth, cracks form polygonal areolas, 0.2 × 1.2 mm. Areole surface flat to slightly convex, usually delimited by black margins. Hypothallus not present. Thallus in section paraplectenchymatous. Mycobiont cells from irregularly globose to polygonal, 3–6 × 2–4 µm (*n*=25, *s*=6). A translucent refringent layer with cells with lipid content, basal layer is present, KI+ magenta blue. Phaenocortex absent, sometimes a thin, discontinuous layer 10–20 µm thick is visible in some areas (*n*=5, *s*=3). A brown pigment layer, K-, is located above photobiont layer. Photobiont arranged in columns throughout the thallus, from irregularly ellipsoidal to globose, (10–)11–16(–17) × (6–)7–12(–13) µm (*n*=43, *s*=6).

*Perithecia* immersed to semi-immersed, up to 1/4, shiny black colour, round in shape with flattened tip. In section, from globose-ellipsoidal to pyriform, (120–)150–250(–260) × (110–)140–225(–230) µm (*n*=20, *s*=5); ostiolar region slightly papillose, typically concolour with the rest of perithecia, ostiolar opening visible at 10×. Involucrellum present, (20–)25–40(–45) µm thick (*n*=11, *s*=5), extends only in the upper part of the perithecium. Excipulum hyaline, 10–18(–20) µm thick (*n*=12, *s*=5), composed of c. 5–6 layers. of flattened rectangular cells. Periphyses present, septate, simple, (10–)15–20 × 1–1.5. Paraphyses are absent. Hamathecial gelatine K-, I + red, K/I + blue. Asci bitunicate, exotunica disappears rapidly with maturity, clavate, 8-spored, located in the lower and lateral parts of the perithecial cavity, (30–)35–50(–55) × (7–)10–12(–13) µm (*n*=14, *s*=5). They are located on the lower and lateral part of the ascoma. *Ascospores* simple, hyaline, ellipsoidal, (9–)10–11(–12) × (4.5–)5–6(–7) µm (*n*=104, *s*=4).

*Pycnidia* present, immersed, irregularly distributed on the thallus, brownish, inconspicuous, ostiolar region concolour with the rest of pycnidia; from globose to bacilliform, (80–)85–110(–112) × (45–)50–70(–75) µm (*n*=10, *s*=3), pycnidial wall hyaline. Conidiospores from bacillar to some curved, hyaline, 3–5 × 1 µm (*n*=54, *s*=3).

##### Additional specimens examined

Chile, Aysén Region, Aysén province, Chonos Archipelago, coast NE of village Puerto Aguirre, 73°31’01’W, 45°09’31’S, on rocks. 0-5 m alt. 17 February 2016. *U. Schiefelbein* 4336 (Herbarium Musei Britannici) – Chile, Los Rios Region, Valdivia Province, Corral, Playa Pelche, 73°40’28’W, 39°58’43’S, on rocks. 0-5 m alt. 22 February 2016. *U. Schiefelbein* 4358 (Herbarium Musei Britannici) – Chile, Region Los Lagos, Province of Palena, Huinay, Comau Fjord, Intertidal rocks at Punta Llonco, 42°20’36”S 72°27’24”W, on rock. 0 m alt. 19 December 2014. *S. Pérez-Ortega* 3538 – Chile, Magallanes and Chilean Antarctic Region, Province of Última Esperanza, Natales, Seno Última Esperanza, coast S of Puerto Natales, 72°29’41”W, 51°48’45”S, on sedimentary rock. 0 m alt. 17 January 2023. *U. Schiefelbein* 6567 – Chile, Los Lagos Region, Quemchi, Chiloé Island, Pinquén beach. Intertidal and supralittoral, 42°8’35”S 73°28’15”W, on rock. 1-5 m alt. 2 December 2009. *S. Pérez-Ortega* 4 *s. n.* (MA-Lich) – Falkland Islands, East Falkland, Darwin Harbour, 51°48’68’S 58°57’60’W, on rock. 0-5 m alt. 1 February 2015. *A. Orange* 22634 (Welsh National Herbarium NMW).

The following key is presented to identify the species and undescribed lineages in the southern hemisphere *marine Verrucarias* group:

1a. Thallus growing on rock 2

1b. Thallus growing on mussel shells (*Mytilus sp.*) ***V. orangei***

2a. (1a) Thallus continuous to sparingly rimose 3

2b. Thallus richly and deeply rimose to areolate 7

3a. (2a) Thallus with numerous punctae ***V. austroditmarsica***

3b. Thallus lacking punctae 4

4a. (3 b) Thallus living completely submerged for his entire life ***V. serpuloides***

4b. Thallus living in the intertidal zone, periodically submerged. 5

5a. (4b) Thallus thin (60–100 µm), semi-immersed to sessile perithecia. ***V. yagani***

5b. Thallus thick (150–280 µm), semi-immersed to immersed perithecia. 6

6a. (5b) Perithecia with papillose ostiolar region, hyaline excipulum somewhat brownish near involucrellum. Conidiospores bacillary to slightly curved (5–6 × 1 µm) ***V. pseudomucosa***

6b. Perithecia with non-papillose ostiolar region, hyaline excipulum. Conidiospores bacillary (4–5 × 1 µm) ***V. chonorum***

7a. (2b) Involucrellum scabrid 8

7b. Involucrellum smooth 11

8a. (7a) Perithecia big (200–250 × 180–275 µm). Spores of 8–10 × 5–6 µm 9

8b. Perithecia small (120–180 × 110–140 µm). Spores of 8–9 × 4–5 µm 10

9a. (8a) Perithecia 200–250 × 200–275 µm. Ostiolar opening barely visible at 10×. Photobiont cells irregularly ellipsoidal to globose, 8–10 × 5–9 µm ***V. pseudodispartita***

9b. Perithecia 235–250 × 180–200 µm. Ostiolar opening no visible at 10×. Perithecia can form groups. Photobiont cells irregularly ellipsoidal to globose, 5–8 × 4–6 µm ***V. delosriosii***

10a. (8b) Thallus 25–30 µm in height. Photobiont cells randomly arranged, irregularly ellipsoidal to globose, 5–8× 4–5 µm. Asci 25–30 × 10–11 µm ***V. cuncorum***

10b. Thallus 60–85 µm in height. Photobiont cells randomly arranged but sometimes look like irregular columns, irregularly ellipsoidal to globose, 9–12 × 4–7 µm. Asci 30–45 × 12–14 µm ***V. dispartita***

11a. (7b) Cracks black 14

11b. Cracks between areolae ± concolour with rest of light thallus, 12

12a. (11b) Ostiolar region papillose ***V. papillata***

12b. Ostiolar region not papillose 13

13a. (12b) Perithecia immersed ***V. psychrophila***

13b. Perithecia sessile to semi-immersed, with the involucrellum in extravagant shapes on many occasions…. V. lambi

14a. (11a) Excipulum brown 15

14b. Excipulum hyaline 17

15a. (14a) Perithecia immature ***V. hydropunctarioides***

15b. Perithecia mature 16

16a. (15b) Thallus with numerous punctae. Ascospores simple, hyaline, ellipsoidal, 11–14 × 5–6 µm ***V.*** labyrinthica

16b. Thallus lacking punctae. Ascospores simple, hyaline, ellipsoidal, 13–16 × 6–7 µm ***V. rimosoareolata***

17a. (14b) Thallus with a basal layer of carbonised melanised mycobiont cells ***V. durietzii***

17b. Thallus lacking a basal layer of carbonised melanised mycobiont cells 18

18a. (17b) Thallus from pale brown to yellowish cream, occasionally slightly dark brownish or even slightly maroon. V. tessellatula

18b. Thallus of other colours 19

19a. (18b) Basal layer of the thallus, in section KI+ magenta blue. ***V. austroamericana***

19b. Basal layer of the thallus, in section KI 20

20a. (19b) Thallus slightly areolate, 100–210 µm in height. Phaenocortex discontinuous layer 10–15 µm. Photobiont cells 10–15 × 7–11 µm. Ostiolar opening visible at 10×. *Perithecia* immersed to semi-immersed, up to 1/3. V. ceuthocarpoides

20b. Thallus strongly areolate, 70–150 µm in height. Phaenocortex absent. Photobiont cells 10–14 × 8–11. Ostiolar opening barely visible at 10×. Perithecia immersed to semi-immersed, up to 1/4. ***V. shukakensis***

## DISCUSSION

The integrative approach used in this taxonomic revision have demonstrated that the taxonomic diversity of the marine *Verrucariaceae* in the Southern Hemisphere has been significantly underestimated. Previous to this study, c. 21 species of marine *Verrucaria* were known from the Southern Hemisphere (Dodge 1948, 1973; Galloway 2007; Lamb 1948a; McCarthy 2023; Santesson 1939). The combination of extensive sampling in Southern South America and Antarctic Peninsula, comprehensive morpho-anatomical studies and the utilisation of molecular data has enabled the recognition of 28 species-level lineages, 21 of which are presented here as new species.

In addition to the species typically restricted to the Southern Hemisphere, several species typically thriving in the European coasts have been reported occurring austral coasts (Galloway 2007; Lamb 1948a; Santesson 1939), showing typical bipolar distribution ranges (Garrido-Benavent and Pérez-Ortega 2017). These species, namely *V. ceuthocarpa, V. degelii, W. mucosa* and *W. striatula,* have not been recorded during our study. However, morphological similar species have been found in the region, which could have been led to confusion in the past. For instance, *V. sp. 15* and *16* (Fig. 17 and 18) can be readily mistaken for *W. mucosa*, given their strikingly similar thallus and perithecia. However, a key differentiating factor is the length of their spores, which are slightly longer (13–15/10–14) than those of *W. mucosa* (11–13). Another species that can be readily mistaken for another is *W. striatula* with *V. sp 18* (Fig. 20). Both possess identical thallus morphology and the distinctive ridges. However, the presence of *W. striatula*, *V. ceuthocarpa* and *W.mucosa* cannot still be discarded, and it would be prudent to check the cited material in old literature to completely reject this possibility.

Considering the candidate new taxa proposed in this revision, it should be noted that some of the species are described based in a single or low number of specimens. For example, this is exemplified by *V. sp 11, 1, 2* or *3*. In the past, this approach was discouraged and regarded as a suboptimal scientific practice (Cheek *et al*. 2020). However, new conceptualizations recently brought by Cazabonne *et al*. 2024 encourage the publishing these ‘sigleton-based species’ in the fungal kingdom. Members of the family Verrucariaceae are often minute, typically challenging to observe, collect, and identify in the field. This is not different in the case of marine Verrucariaceae. In our case, we can also add on top of that the remote area surveyed during this study, in which sampling often entail long and expensive logistically complex field campaigns. Consequently, we believe that the description of new taxa based on a small number of specimens or singletons is fully justified, given the broad sampling and thorough morphological and molecular investigations.

The lack of abundant morphoanatomical characters, together with the high plasticity found in some of them, has complicated and challenged the correct delimitation of species in many groups of lichen-forming fungi (Lumbsch and Leavitt 2011), including marine species (Orange 2012). The interpretation of the variation of subtle differences between some specimens and its significance at different taxonomic, ecological or geographic scales is often impossible without the help of molecular data to support and test taxonomist’s species hypotheses.

The use of a barcoding approach analysing a large number of samples, as used in our study, is a recommended strategy in groups such as lichen-forming fungi (Crespo and Pérez-Ortega 2009; Dal Forno 2022; Divakar *et al*. 2016; Pino-Bodas *et al*. 2012; Savić and Tibell 2009; Spribille *et al*. 2011; Vondrák 2020, 2022). and, particularly in groups with a scarce number of taxonomic characters and high plasticity such as marine Verrucariaceae. Thanks to this approach, together with the use of species delimitation algorithms, we have been able to unearth hidden diversity that could have very likely be overlooked if only selected specimens would have been chosen for multi-locus analyses. This is the case of the *V. dispartita* group, a complex of species with highly similar morpho-anatomical characteristics. The differences between the species of this group are both subtle and complex to interpret. However, the use of barcoding has demonstrated that there are, in fact, four distinct species (*V. sp. 1, 2, 3* and *7*) that diverge from the *V. dispartita* known from the northern hemisphere. Moreover, some of these species are not even related to the rest, as is the case of *V. sp 7*.

With regard to the delimitation algorithms, both ASAP and mPTP produced significantly more conservative results and were more similar to the species delimited by morphological data than GMYC, which produced a higher number of candidate species, especially multiple GMYC, which is known to split clades into candidate species, as has been observed in other groups of organisms, including lichen-forming fungi (Blázquez *et al*. 2024; Dellicour and Flot 2015; Pentinsaari *et al*. 2017). Our dataset, the number of representatives of each species in our phylogeny highly differs. This is a situation where it has previously stated that may cause significative errors in the delimitation of candidate species using single-locus algorithms (Ahrens *et al*. 2016; Blair and Bryson 2017). In our case, we have not encountered any issues with the singletons, as illustrated in Fig. 2. This figure demonstrates that the majority of algorithms have distinguished these species, which only have one representative, from the rest. It is also significantly influenced by taxonomic rank, showing lower efficiency in genus-level datasets compared to higher taxonomic levels (Guo and Kong 2022). The analyses were conducted on the *Mastodia* and *Turgidosculum* clades collectively and subsequently on an individual basis. The results obtained were consistent across both approaches.

Generic delimitations in Verrucariaceae has been shown to be problematic under the light of molecular data, with morphologically delimited groups often discovered to be be polyphyletic (Gueidan *et al*. 2007, 2009; Prieto *et al*. 2010; Savić *et al*. 2008). The marine species of the family have so far been separated into four different genera, all of which are well-defined phylogenetically, but with little morphological support, except in the case of *Hydropunctaria* (Gueidan *et al*. 2009, 2022; Orange 2012, Pérez-Ortega *et al*. 2018). The species studied and described during our study are placed in two distinct clades within the Verrucariceae, which an obvious lack of morphological homogenity. A small group group of species are placed in the *Turgidosculum* clade (Pérez-Ortega *et al*. 2018). Most species form a single clade around the well-known taxon *Mastodia tessellata*. Species within these two clades showed limited similarities in terms of their morphology.

The species placed in the *Turgidosculum* clade (*Verrucaria* sp. 1, *Verrucaria* sp. 2, *Verrucaria* sp. 3, and *Verrucaria* sp. 4) showed some similarities. For instance, these species exhibit a scabrid involucrellum, accompanied by the presence of punctae in very thin thalli the thallus to varying degrees. These characters are shared by the European species *Verrucaria ditmarsica*, but not by the North American *T. ulvae*, both members of this clade (Pérez-Ortega *et al*. 2018), neither by new proposed taxa *Verrucaria* sp. 6.

Regarding the *Mastodia* clade, some smaller subclades showed same synapomorphies. Thus, *Verrucaria* sp. 12 and *Verrucaria* sp. 14, and the phylogenetically related *V. durietzii* share a large and prominent involucrellum that may, on occasion, envelop the perithecia. On the other hand, *Verrucaria* sp. 15 and *Verrucaria* sp. 16, both closely related with *V. serpuloides*, exhibit comparable characteristics in their thallus and perithecia. Whether those smaller clades should be treated as different genera or whether all the taxa in this group should be placed within the genus *Mastodia* is, for the moment, an open debate. Considering the size, lack of morphological congruence and branch length of the genera proposed so far for the European taxa, one could advocate for a large number of genera to accommodate Southern species. The use of a temporal banding approach (Divakar *et al*. 2017; but see Lücking 2019) could serve, in the absence of shared synapomorphies, as a guide to propose genera with similar criteria.

The topology of the deep relationships between genera and large clades in the Verrucariaceae are still poorly known (Gueidan *et al*. 2007, 2009). Lack of sufficient molecular characters hinders the achievement of supported relationships at this level. In our phylogenetic hypothesis, the species from Southern South America and Antarctica fall into two different lineages (Fig. 1). On the one hand, *Verrucaria* sp. 1 to sp. 6 were situated, in a clade sister to *Turgidosculum ulvae* from the northern hemisphere. The whole clade is well supported by both phylogenetic inference methods and it is separated from the rest of Verrucariaceae by a characteristic long branch, already showed by Pérez-Ortega *et al*. (2018). This long branch may be a sign of a past massive extinction in the group (Crisp & Cook 2009).

In the case of the remaining species, they are all placed in a clade containing the species *Mastodia tessellata*. Pérez-Ortega *et al*. (2010) already showed the phylogenetic affinities of *Mastodia tessellata* with other species of marine Verrucariaceae from Southern Chile. This clade was not well-supported by any of the two inference methods, likely due to the low number of loci obtained for many species, especially the nrSSU and the very informative RPB1, a region has been shown to enhance phylogenetic tree resolution in lichen-forming fungi (Hofstetter *et al*. 2007; Reeb *et al*. 2004; Truong *et al*. 2013). This fact diminishes the amount of information available to produce reliable phylogenetic hypotheses. The Southern clade formed a well-supported clade together with *Verrucariopsis*, a recently described genus from European coasts, which shows a sister relationship with the species in the genus *Wahlenbergiella*.

Our results also showed that two European species, currently in Verrucaria, namely to *V. ceuthocarpa* and V*. degelii*, appeared related to other species in the genus *Wahlenbergiella*, so consequently, they should be combined into that genus.

## CONCLUSIONS

1. Extensive sampling coupled with a barcoding strategy has facilitated a notable advancement in our understanding of the family Verrucariaceae within the marine ecosystems of southern South America and Antarctica and to overcome the problems inherent to a problematic group due to the scarcity of characters and their high plasticity. A total of 28 lineages have been identified, 21 of which may represent new species for scientific research. It seems probable that the diversity of the group in the area is considerably greater than that indicated by our survey, which covered only a small fraction of the territory.
2. The results demonstrated that marine Verrucariaceae species from the southern hemisphere are placed in two distinct and non-related lineages within the family. Further studies incorporating additional molecular data and a temporal banding approach are required to ascertain the nomenclatural consequences of these findings.

